# Mice carrying nonsense mutant p53 develop frequent multicentric or metastatic tumors

**DOI:** 10.1101/2025.03.12.642796

**Authors:** Charlotte Strandgren, Veronica Rondahl, Ann-Sophie Oppelt, Susanne Öhlin, Angelos Heldin, Klas G. Wiman

## Abstract

The *TP53* tumor suppressor gene is mutated in a large fraction of human tumors. Close to 11% of *TP53* mutations are nonsense mutations, causing premature termination of protein synthesis and expression of truncated inactive p53 protein. The most common *TP53* nonsense mutation in human cancer is R213X. To study the impact of *TP53* nonsense mutations *in vivo*, we generated mice harboring the *Trp53* nonsense mutation R210X that corresponds to human *TP53*-R213X. Initially, *Trp53*^R210X^ mice appear phenotypically normal, although the proportion of female *Trp53*^R210X/R210X^ mice is dramatically reduced. Female homozygous mice are poor breeders and remain smaller and lighter than female heterozygous and wildtype littermates. *Trp53*^R210X/R210X^ mice start to show tumors at 2.5 months of age, and their maximal lifespan is 8.5 months. *Trp53*^R210X/+^ mice present tumors from 9 months of age, and by 16.5 months of age 50% of all heterozygous mice have developed overt tumors. 71% of tumors from *Trp53*^R210X/+^ mice show loss of heterozygosity (LOH). Homozygous mice develop hematopoietic and mesenchymal tumors, most commonly T-cell lymphoma and leiomyosarcoma, and heterozygous mice develop hematopoietic, mesenchymal, epithelial and sex cord tumors, most commonly osteosarcoma and leiomyosarcoma. The tumor phenotype is similar to that of *Trp53-null* and *Trp53-*missense knock-in mice, although the *Trp53*^R210X/R210X^ mice have a high rate of multicentric or metastatic tumors, and *Trp53*^R210X/+^ mice have a longer overall survival than *Trp53*^R172H/+^ missense mutant knock-in mice. Treatment of T-cell lymphoma cells from *Trp53*^R210X/R210X^ mice with aminoglycoside G418 induces expression of full-length functional p53 and apoptotic cell death. Our new unique mouse model will allow further studies of the effects of *Trp53* nonsense mutation in a multi-organ system and serve as a model for the Li-Fraumeni syndrome (LFS). It will also be valuable for preclinical evaluation of novel therapeutic strategies for targeting *TP53* nonsense mutations in cancer.

## Introduction

The transcription factor p53 is activated by cellular stress (1). Activated p53 binds to specific DNA motifs as a tetramer, transactivates multiple target genes (2, 3), and controls multiple processes, including cell cycle progression, DNA repair, metabolism, senescence, and apoptosis (1, 4). The *TP53* gene is often inactivated in human tumors, most frequently by missense mutations that disrupt p53 DNA binding and transactivation of target genes (5, 6). Missense mutant p53 can inhibit co-expressed wildtype (WT) p53 through a dominant-negative effect that can increase resistance to genotoxic stress and drive clonal selection (7). Some forms of missense mutant p53 may exert oncogenic gain-of-function (GOF) activity, associated with a wider tumor spectrum and enhanced invasion and metastasis (8–10). However, other studies have not found evidence for mutant p53 GOF activity (7, 11, 12) and it is likely that mutant p53 GOF depends on cell background. For instance, myeloid malignancies may behave differently than solid tumors (7). A recent detailed characterization of *TP53* variants in HCT116 cells demonstrated R175H mutant p53 GOF only after long-term *in vitro* passaging, associated with increased mutant p53 levels and manifested as enhanced migration, invasion and metastasis (13). Around 11% of *TP53* mutations are nonsense mutations that result in a premature stop codon (6) and expression of truncated p53 that lacks critical functional domains and therefore fails to transactivate p53 target genes. Nevertheless, certain truncated p53 proteins generated by *TP53* nonsense mutations may have oncogenic properties (14).

Mouse models that either completely lack p53 or carry missense mutant p53 (*Trp53*^Mis^) support a key role of p53 as tumor suppressor. Homozygous *Trp53*-null or *Trp53*^Mis^ mice show a striking cancer phenotype with a high rate of spontaneous tumors and reduced survival compared to WT mice (9, 15–20). Both *Trp53*^-/-^ and *Trp53*^Mis/Mis^ mice exhibit tumors from 3-6 months of age, whereas *Trp53*^+/-^ and *Trp53*^Mis/+^ mice display tumors at 9-10 months of age, often associated with loss of the WT allele (loss of heterozygosity, LOH). *Trp53*^-/-^ and *Trp53*^Mis/Mis^ mice mainly develop lymphomas and sarcomas, whereas heterozygotes, especially the *Trp53*^Mis/+^ models, also develop carcinomas. *Trp53*^R172H/+^ and *Trp53*^R270H/+^ mice develop more invasive and metastasis prone tumors compared to *Trp53*^+/-^ mice, consistent with GOF activity for these mutants (9, 17). However, a mouse model of colorectal cancer with inactivated *Apc* and activated *Kras* alleles did not show any significant difference between *Trp53*-null and R270H mutant alleles in life span, adenocarcinoma incidence and metastasis (21).

Mouse models designed to investigate the functional implications of *TP53* nonsense mutations *in vivo* have previously not been established. Such models are crucial for a better understanding of the impact of nonsense mutant p53 in a living organism and can also serve as platforms for preclinical development of pharmacological strategies for therapeutic targeting of nonsense mutant *TP53* in cancer. Several compounds can induce translational readthrough of nonsense mutant *TP53* and thus expression of full-length and functional p53 protein in cultured cells, including G418 (22), Clitocine (23), ELX-02 (24), 2,6-Diaminopurine (25), cc-90009 (26), and 5-Fluorouridine (27). Here we present a novel mouse model carrying the *Trp53*-R210X nonsense mutation that corresponds to *TP53*-R213X, the most common *TP53* nonsense mutation in human tumors (6).

## Results

### Generation of a *Trp53*^R210X^ mouse model

To generate a mouse model harboring the *Trp53*-R210X nonsense mutation, we used CRISPR/Cas9 genome editing to introduce two single point mutations resulting in an Arg (CGC) to STOP (TGA) substitution at codon 210 in exon 6 (Fig. 1A). The CRISPR/Cas9 strategy was validated in NIH/3T3 mouse fibroblasts. Single-cell clones of edited NIH/3T3 cells were genotyped as heterozygous using the introduced XbaI cleavage site. PCR amplicons of *Trp53* exons 5-7 from the NIH/3T3 cell clones were subcloned and sequencing of individual plasmid clones identified those carrying the R210X mutation and the two downstream silent mutations. However, the XbaI site was not present in all clones (Fig. 1B). ssODN template-triggered repair can generate short conversion tracts of about 30 nucleotides with Gaussian-like distributions surrounding the Cas9 cut site (28). The mutation producing the intended XbaI site was too far from the Cas9 cut site and therefore outside of the effective conversion zone, explaining its inconsistent introduction.

**Figure 1.**
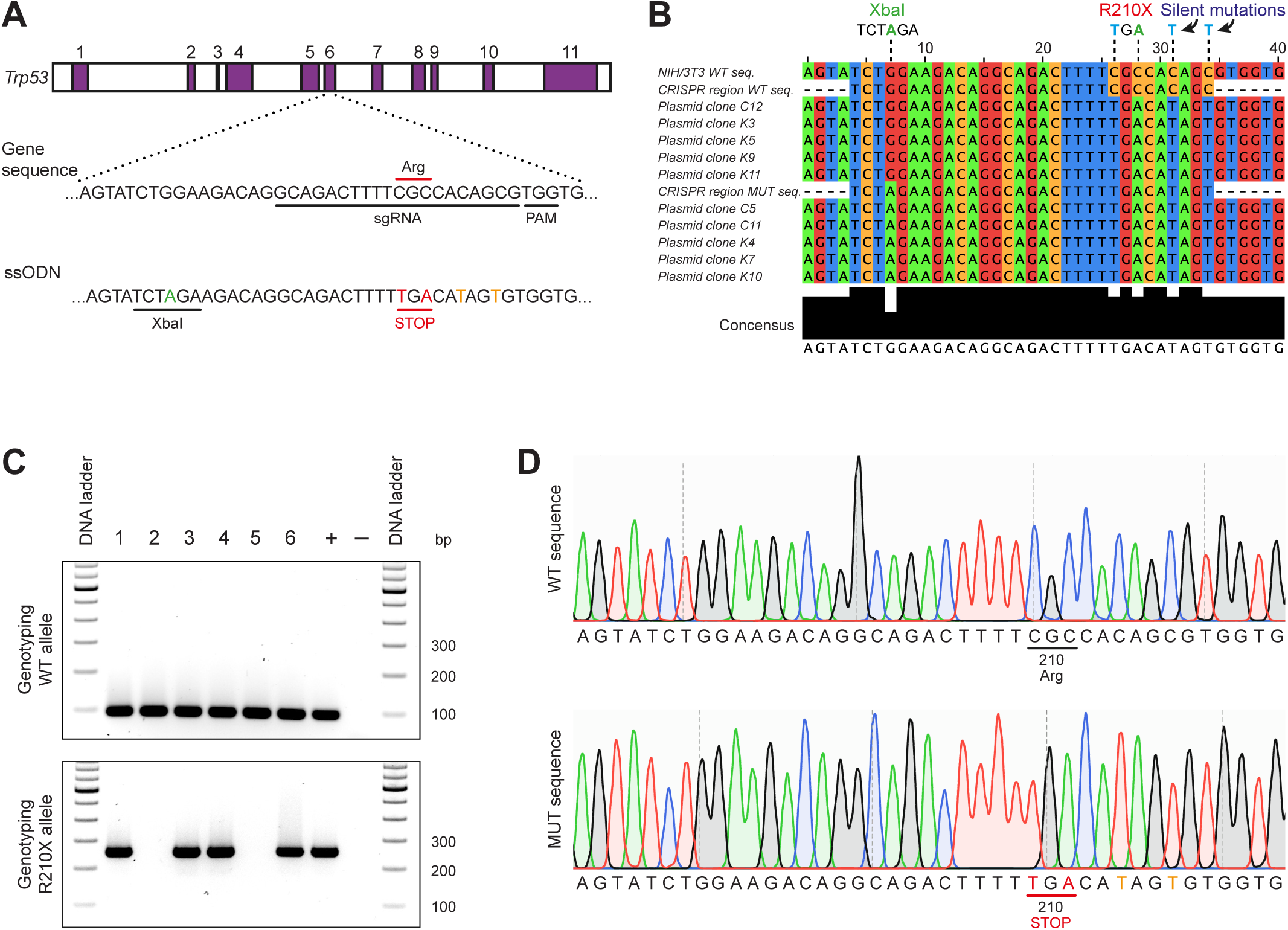
Generation of the *Trp53*^R210X^ knock-in mouse strain. **(A)** Schematic representation of the CRISPR/Cas9 strategy for generation of *Trp53*^R210X^ mutant mice. A single-guide RNA (sgRNA) was designed to target the region encoding the R210 codon in *Trp53* exon 6. The expected cleavage site is 3 nucleotides upstream of the Protospacer Adjacent Motif (PAM) sequence. The donor single-stranded oligodeoxynucleotide (ssODN) template was designed to introduce nucleotide replacements resulting in an Arg (CGC) to STOP (TGA) substitution (indicated in red) at codon 210 (R210X), and an XbaI cut site to facilitate genotyping by restriction enzyme digestion (introduced nucleotide shown in green). Silent mutations introduced to prevent recutting of the sgRNA are shown in orange. The figure is not to scale. **(B)** Representative results from sequence analysis of plasmid clones from edited NIH/3T3 cell clones, showing a correct R210X sequence but inconsistent introduction of the XbaI site. **(C)** Genotyping strategy using wildtype (WT) and mutation specific primers for PCR amplification. PCR products were run on a 2% agarose gel. WT allele is identified by a 95 bp PCR fragment (top panel), and mutant allele by a 252 bp PCR fragment (bottom panel). **(D)** Representative chromatograms showing clean WT (upper panel) and mutated (lower panel) *Trp53* sequences from heterozygous F1 mice, indicating successful knock-in of desired mutations and germline transmission of the R210X allele. Altered nucleotides producing the STOP codon (TGA) at amino acid position 210 are shown in red, silent mutations are shown in orange.

Genotyping the F0 offspring with PCR revealed that 6/17 pups were positive for the R210X mutation. Sanger sequencing confirmed that two of the six pups also had the XbaI site, whereas the remaining four lacked it (not shown). The pups were also genotyped with restriction enzyme digestion, which confirmed the two pups carrying the Xbal site, and also identified one pup carrying the Xbal site but not the correctly mutated R210X sequence. The six F0 offspring carrying the correct R210X mutation were set up for backcross breeding to WT C57BL/6J mice to eliminate possible off-target events. Two founder animals, both lacking the XbaI site, generated F1 offspring, and PCR genotyping verified germline transmission of the R210X allele (Fig. 1C). Sequencing of 1049 bp surrounding the edited *Trp53* exon 6 locus confirmed clean WT and R210X alleles in all F1 mice identified as heterozygous (Fig. 1D).

### Intercrosses of *Trp53*^R210X^ mice yield fewer and smaller female homozygous offspring

The *Trp53*^R210X^ strain was maintained by backcrossing *Trp53*^R210X/+^ mice to WT C57BL/6J mice. Backcross litter sizes at 2 weeks of age were normal compared to the reference litter size of 7.0 pups for C57BL/6J mice (https://informatics.jax.org) (Supplementary Table S1), and genotype and sex distribution (309 offspring) was consistent with expected Mendelian inheritance but with significantly more WT males (Supplementary Table S2). Different combinations of intercrosses were set up to assess the viability and reproductive capacity of *Trp53*^R210X/R210X^ mice (Supplementary Table S1). Intercrossing only *Trp53*^R210X/R210X^ mice resulted in litters with significantly fewer pups at 2 weeks of age compared to backcrosses, whilst intercrosses with at least one *Trp53*^R210X/+^ mouse yielded similar litter sizes as the backcrosses (Supplementary Table S1; Supplementary Fig. S1). Among the 13 intercrosses between *Trp53*^R210X/R210X^ and *Trp53*^R210X/+^ mice, only one litter was derived using a *Trp53*^R210X/R210X^ female, yielding a single male *Trp53*^R210X/+^ pup. All other 12 litters were spawned from *Trp53*^R210X/+^ females. Thus, *Trp53*^R210X/R210X^ female mice are fertile but have a reduced breeding capacity.

Intercrosses between *Trp53*^R210X/+^ and/or *Trp53*^R210X/R210X^ mice gave fewer female *Trp53*^R210X/R210X^ offspring (Supplementary Table S2), whereas male offspring distribution followed expected Mendelian inheritance regardless of genotype. Females remained fewer when all *Trp53*^R210X/R210X^ offspring were analyzed. Aside from the poor breeding capacity of female *Trp53*^R210X/R210X^ mice, heterozygous and homozygous mice of both genders appeared phenotypically normal at young age and were fertile.

*Trp53*^R210X^ mouse bodyweights were recorded weekly from postnatal week 2 and until euthanasia for all offspring from homozygous-generating intercrosses. Female *Trp53*^R210X/R210X^ mice had smaller body size (not shown) and were lighter than both WT and *Trp53*^R210X/+^ littermates at 2 weeks of age and remained lighter throughout life (Fig. 2A). In contrast, male mice had almost identical weight development regardless of genotype until 34 weeks of age, when the remaining *Trp53*^R210X/R210X^ males were markedly lighter than their *Trp53*^R210X/+^ and WT littermates, most likely due to emaciation secondary to tumor growth (Fig. 2B).

**Figure 2.**
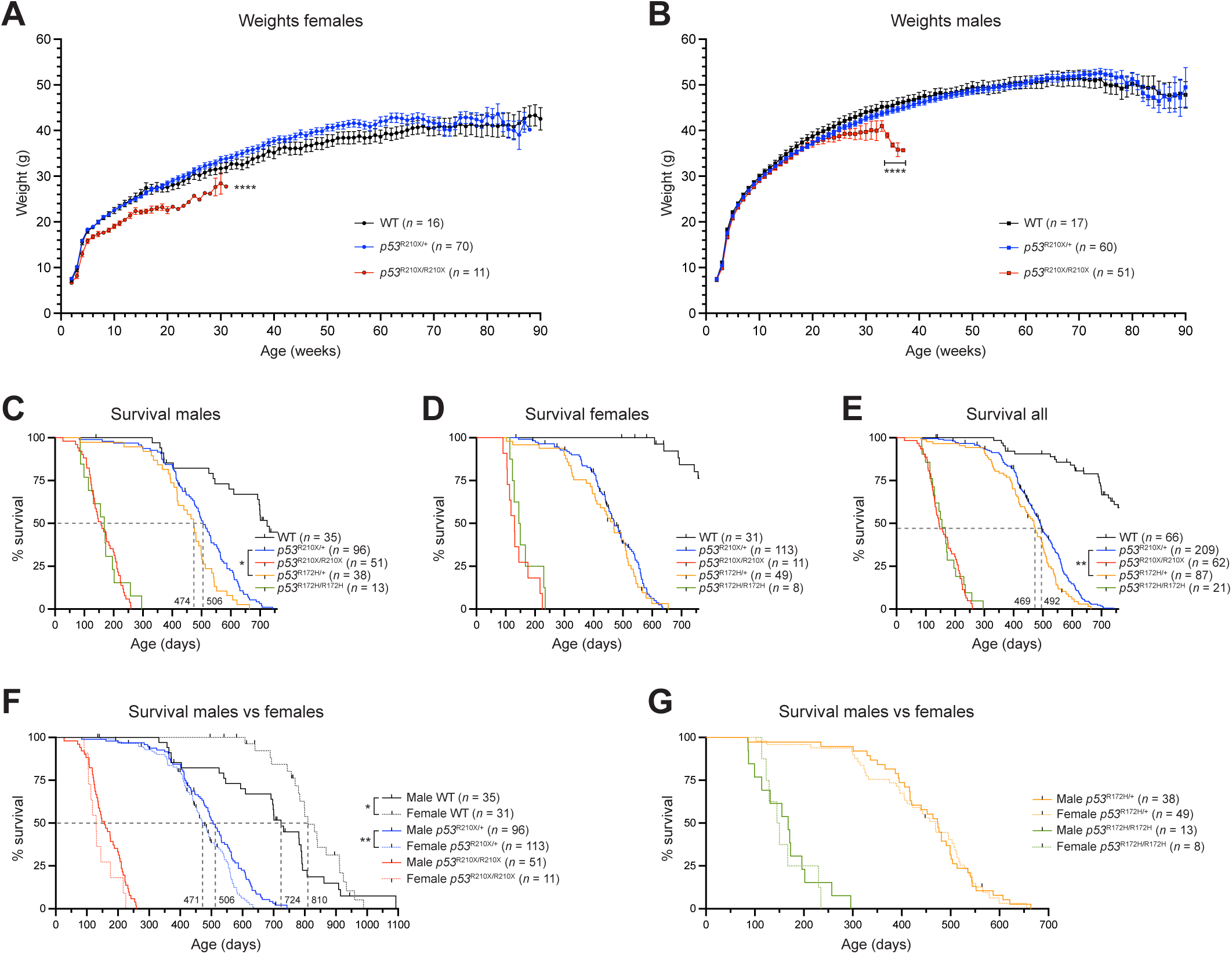
Female *Trp53*^R210X/R210X^ mice are significantly lighter and *Trp53*^R210X^ mice have a reduced lifespan. Bodyweight development of female **(A)** and male **(B)** *Trp53*^R210X/+^ and *Trp53*^R210X/R210X^ mice as compared to wildtype (WT) littermates. **(A)** Female *Trp53*^R210X/R210X^ mice were significantly lighter during their whole lifespan as compared to *Trp53*^R210X/+^ and WT littermates (\*\*\*\**p* < 0.0001). **(B)** Male mice of the different genotypes showed no bodyweight differences until 34 weeks of age. From this point, at the end of their maximum lifespan, male *Trp53*^R210X/R210X^ mice were significantly lighter than *Trp53*^R210X/+^ and WT littermates (\*\*\*\**p* < 0.0001). Indicated *n*-values are the total number of mice monitored from 2 weeks of age. Only weights from mice scored as healthy according to general health assessment are shown in **A-B**, as a mouse with tumor burden tends to either dramatically lose weight or gain weight. **(C-E)** Kaplan-Meier curves illustrating the effects of the *Trp53*^R210X^ mutation on survival. WT littermates and *Trp53*^R172H^ missense mutant knock-in mice are shown for comparison. All inter-strain comparisons between different genotypes of both *Trp53*^R210X^ and *Trp53*^R172H^ strains were highly significant (adj. P-values: *p* = 0.0008) (WT vs. heterozygous; WT vs. homozygous; heterozygous vs. homozygous); only P-values for comparison of *Trp53*^R210X/+^ and *Trp53*^R172H/+^ mice are indicated in the figure. **(C)** Male *Trp53*^R210X/+^ mice showed significantly longer survival compared to *Trp53*^R172H/+^ mice (adj. P-value: *p* = 0.013), but the difference between the female heterozygous mice of the two strains was not statistically significant **(D)**. **(E)** When male and female mouse survival data were combined, *Trp53*^R210X/+^ mice lived significantly longer than *Trp53*^R172H/+^ mice (adj. P-value: *p* = 0.01). There was no difference in the survival between *Trp53*^R210X/R210X^ and *Trp53*^R172H/R172H^ mice of either sex, or when data for male and female mice were combined. **(F-G)** Comparison of male and female survival of the different genotypes showed significant differences for *Trp53*^R210X/+^ mice (*p* = 0.003) and WT mice (*p* = 0.043). Black ticks in panels **C-G** indicate mice that have been censored (i.e. euthanized due to non-cancer related reasons such as wounds from scratching/over-grooming or malocclusion). Bodyweight growth curves in **A** and **B** were analyzed by Mixed-effects analysis with Tukey’s multiple comparisons test. Values represent the mean ± SEM. Survival curves in **C-G** were analyzed by log-rank (Mantel-Cox) test followed by correction for multiple comparisons using the Holm-Šídák method when applicable (**C-E**). P-values: \**p* < 0.05, \*\**p* < 0.01, \*\*\**p* < 0.001, \*\*\*\**p* < 0.0001.

### *Trp53*^R210X^ mice have a reduced lifespan

The *Trp53*^R210X/R210X^ mice had significantly shorter survival than *Trp53*^R210X/+^ and WT mice (adj. P-values: \*\*\**p* < 0.001), as did *Trp53*^R210X/+^ mice compared to WT (adj. P-value: \*\*\**p* < 0.001) (Fig. 2C-E). Female and male *Trp53*^R210X/R210X^ mice had similar survival, whilst female *Trp53*^R210X/+^ mice had reduced survival compared to male (*p* = 0.003) (Fig. 2F). Male WT mice showed a significantly shorter mean survival compared to female WT mice (*p* = 0.043), which may be related to the fact that 11 male WT mice <1.5 years old were found dead or were euthanized due to wounds or abdominal masses. Only one of these had abdominal neoplasia; all others either showed no gross abnormalities or had enlarged seminal vesicles, a normal phenomenon in aging male mice (29).

We also compared *Trp53*^R210X/R210X^ survival to that of *Trp53*^R172H/R172H^ missense knock-in mice maintained in the same animal facility. Survival of both mouse strains showed a similar rapid decline from 3-8.5 months of age, and their mean survival showed no statistically significant difference: 145.5 days for *Trp53*^R210X/R210X^ mice and 155 days for *Trp53*^R172H/R172H^ mice (Fig. 2C-E). In contrast, *Trp53*^R210X/+^ mice survived significantly longer than *Trp53*^R172H/+^ mice (adj. P-value: *p* = 0.01), with a mean survival of 492 days vs. 469 days (Fig. 2E). There were no differences in survival between female and male *Trp53*^R172H^ mice (Fig. 2G).

### Tumor development in *Trp53*^R210X^ mice is highly penetrant

All animals, both euthanized and found dead, were necropsied, and histopathological analysis was performed on available tissues from all *Trp53*^R210X/R210X^ mice and a subset of *Trp53*^R210X/+^ and WT mice. Grossly visible tumors were found in 59 of 62 (95%) *Trp53*^R210X/R210X^ mice, 152 of 209 (73%) *Trp53*^R210X/+^ mice, and 30 of 66 (45%) WT mice. This confirms a highly penetrant tumor phenotype in *Trp53*^R210X^ mice. Histopathological examination was performed on tissues from 49 *Trp53*^R210X/R210X^ mice (those with formalin-fixed tissues available) and 79 (those with formalin-fixed tissues available) of the 85 *Trp53*^R210X/+^ mice with suspected tumors that were randomly selected for LOH analysis (see below). To identify a baseline for common tumors in the WT parents and siblings of the *Trp53*^R210X^ mutant mice, we also examined tumors and tissues from the oldest of the available necropsied WT mice (12 males, 12 females). Among the analyzed mice, 48 (98%) of the *Trp53*^R210X/R210X^ mice and 79 (100%) of the *Trp53*^R210X/+^ mice had histopathologically diagnosable tumors. Twenty (83%) of the selected WT mice had tumors and were thus considered a good representation of the background tumor panorama (Supplementary Table S3).

### Different tumor spectra in *Trp53*^R210X/R210X^ and *Trp53*^R210X/+^ mice

*Trp53*^R210X/R210X^ mice only developed hematopoietic tumors and soft-tissue mesenchymal tumors (Fig. 3; Supplementary Fig. S2A-C; Supplementary Table S4A), whereas *Trp53*^R210X/*+*^ mice showed a broader tumor spectrum with hard-tissue mesenchymal tumors, soft-tissue mesenchymal tumors, hematopoietic tumors, epithelial tumors, and a single sex cord tumor (Fig. 4A-D; Supplementary Fig. S3A-C; Supplementary Table S4B), which included both malignant and benign tumors (Supplementary Table S3). WT mice had a broad spectrum of both malignant and benign tumors (Supplementary Table S4C; Supplementary Fig. S4A-C). Malignant tumors were classified as solitary when they were found in a single organ or tissue, hematopoietic tumors found in at least two organs or tissues were classified as multicentric, and mesenchymal and epithelial tumors found in at least two organs and tissues were classified as metastatic. To assess loss of the WT *Trp53* allele (LOH) in tumors from *Trp53*^R210X/+^ mice, we performed semi-quantitative PCR with the primers for genotyping (Fig. 4E). Out of 132 tumors analyzed from *Trp53*^R210X/+^ mice, 94 (71%) showed LOH, consistent with a strong selection pressure to eliminate the remaining WT allele.

**Figure 3.**
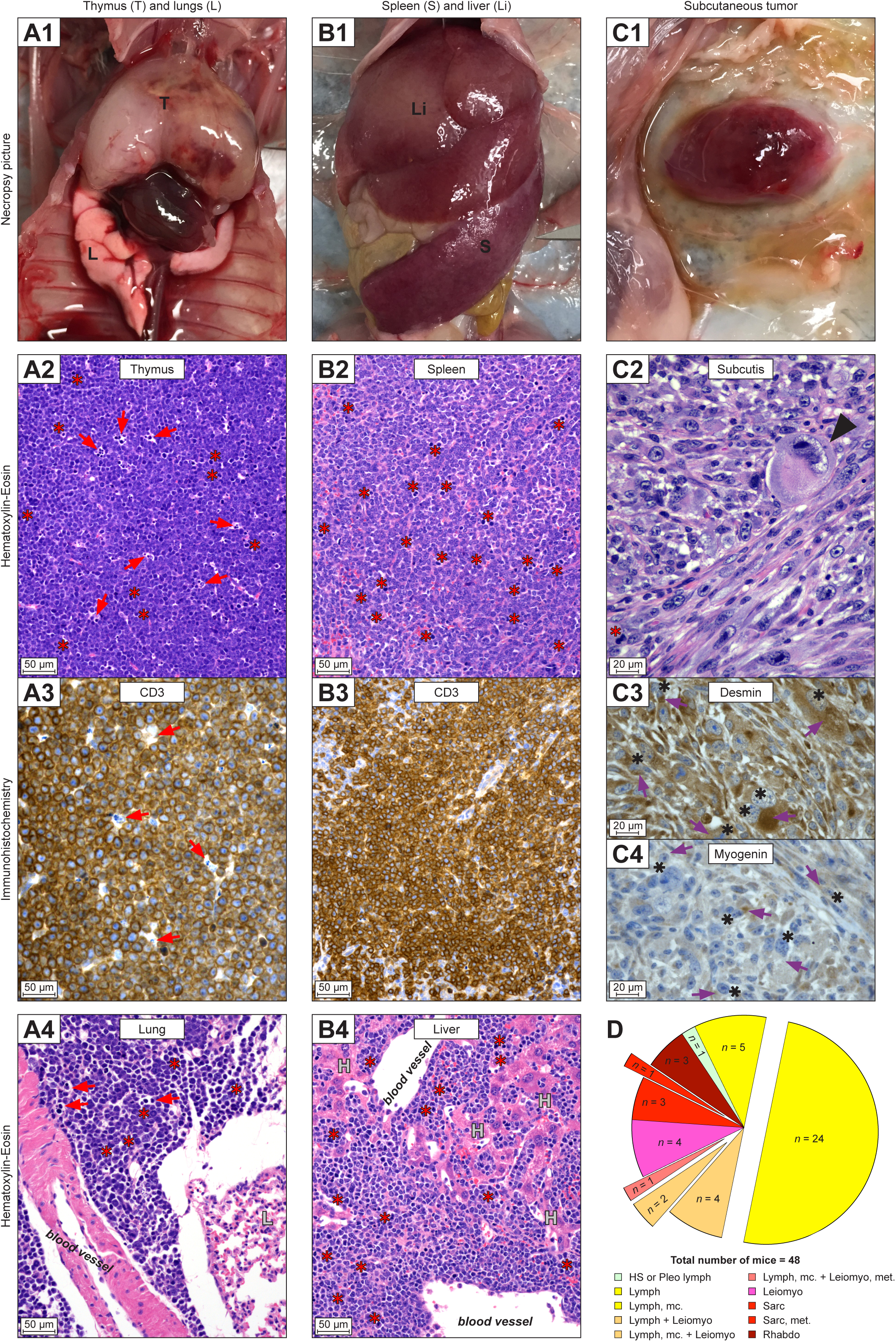
*Trp53*^R210X/R210X^ mice develop multicentric lymphomas and sarcomas. Examples of tumors observed at necropsy and their histopathological diagnosis. All necropsy photographs are oriented with the head at the top. All histomicrographs are stained with hematoxylin-eosin (HE) or immunohistochemistry (IHC) as indicated. Scale bars = 20µm. Red stars = mitotic figures; red arrows = single-cell death; black arrows = tingible body macrophage; purple arrows = cytoplasmic staining; black stars = nuclear staining. **(A1)** A male *Trp53*^R210X/R210X^ mouse with a severely enlarged thymus (T). **(A2)** Histomicrograph of the thymus neoplasia in A1. The normal thymus architecture is completely effaced by mats of neoplastic round cells with dispersed tingible body macrophages phagocytosing dead neoplastic cells (arrows). **(A3)** CD3 IHC confirming that the neoplastic lymphocytes are T-cells. **(A4)** Histomicrograph of lung shown in A1 with perivascular neoplastic lymphocyte infiltrates, L indicates unaffected lung tissue. **(B1)** A male *Trp53*^R210X/R210X^ mouse with a severely enlarged spleen (S) and liver (Li). **(B2)** Histomicrograph of spleen shown in B1. The normal splenic architecture is completely effaced by mats of neoplastic round cells. **(B3)** CD3 IHC confirming that the neoplastic lymphocytes are T-cells. **(B4)** Histomicrograph of liver shown in B1, with severe perivascular and intrasinusoidal neoplastic lymphocyte infiltrates, H indicates hepatocytes. **(C1)** A male *Trp53*^R210X/R210X^ mouse with a subcutaneous tumor on the left side of thorax and abdomen. **(C2)** Histomicrograph of the subcutaneous tumor in C1, with a dense proliferation of pleomorphic neoplastic mesenchymal cells forming streams and bundles. Black arrowhead indicates a giant cell with a bizarre, giant nucleus. **(C3 and C4)** Desmin and myogenin IHC confirming that the neoplastic cells originate from skeletal muscle. **(D)** Distribution of tumors and multicentric or metastasizing tumors in *Trp53*^R210X/R210X^ mice. The total number of mice includes all mice with at least one tumor. To increase clarity, pleomorphic lymphoma and histiocytic sarcoma were only included when they were the only diagnosed tumor(s), not when they occurred concurrently with other tumor types. Therefore, the number of multicentric tumors are lower than in Supplementary Table S3. The full data is presented in Supplementary Tables S4A and S4D. HS = histiocytic sarcoma; Leiomyo = leiomyosarcoma; Lymph = lymphoma; mc = multicentric; met = metastatic; Pleo lymph = pleomorphic lymphoma; Rhabdo = rhabdomyosarcoma; Sarc = sarcoma.

**Figure 4.**
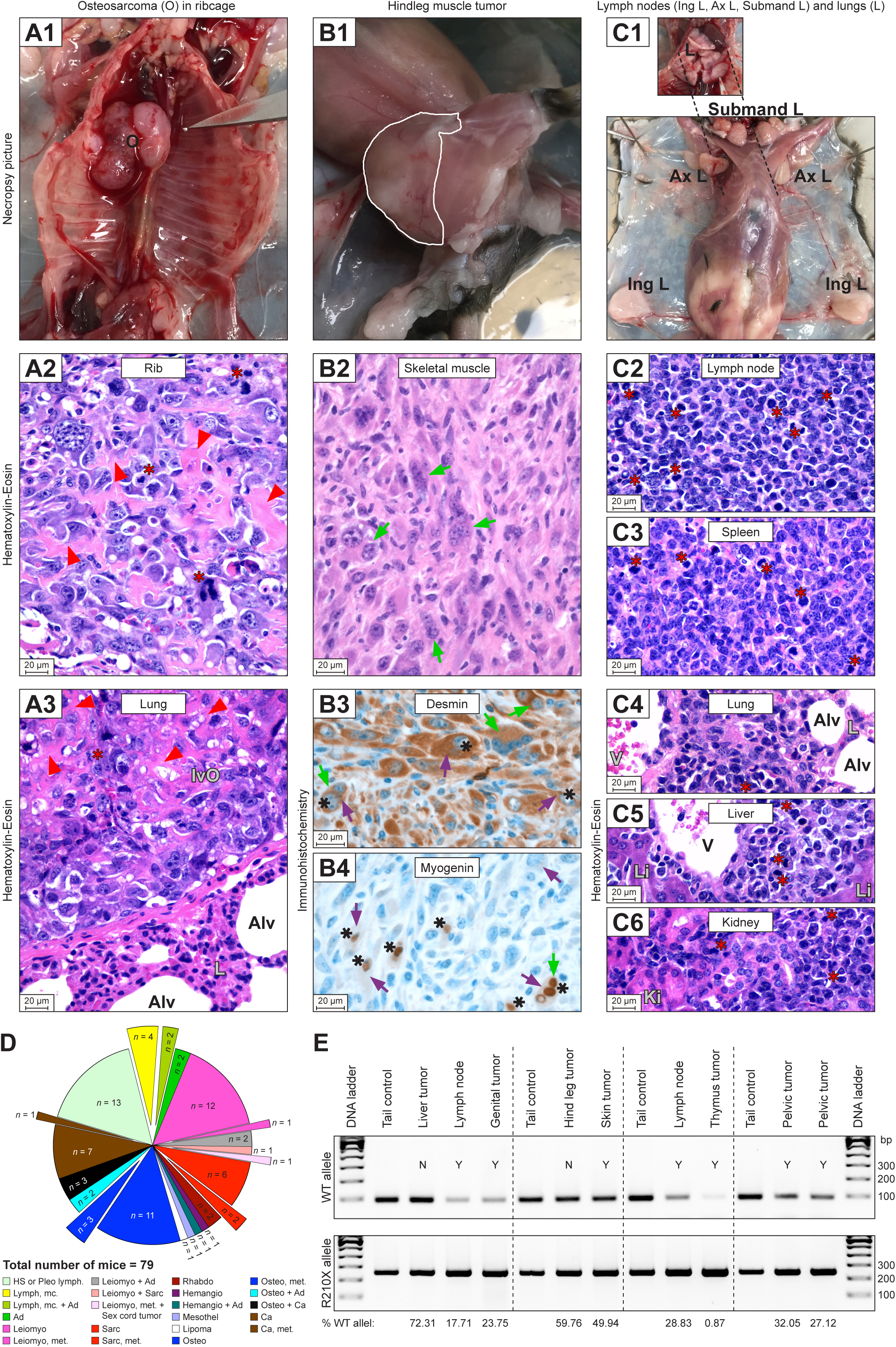
*Trp53*^R210X/+^ mice develop a wide spectrum of tumors with frequent LOH and high rate of multicentric and metastatic tumors. Examples of tumors observed at necropsy and their histopathological diagnosis. All necropsy photographs are oriented with the head at the top. All histomicrographs are stained with hematoxylin-eosin (HE). Scale bars = 20µm. Red stars = mitotic figures, red arrowheads = neoplastic osteoid; green arrows = multinucleate neoplastic cells; purple arrows = cytoplasmic staining; black stars = nuclear staining. **(A1)** A female *Trp53*^R210X/+^ mouse with a hard tumor (O = osteosarcoma) on the thorax wall, involving several ribs. **(A2)** Histomicrograph of the hard tumor in A1, with dense proliferation of very pleomorphic, neoplastic round to spindle-shaped mesenchymal cells that produce variable amounts of neoplastic osteoid, confirming osteosarcoma diagnosis. **(A3)** Histomicrograph of lung from the animal shown in A1, with a large intravascular osteosarcoma metastasis (IvO) adhering to the vessel wall (L = lung parenchyma, Alv = alveolus). **(B1)** A male *Trp53*^R210X/+^ mouse with a hindleg muscle tumor (indicated by the white line). **(B2)** Histomicrograph of the tumor from the animal shown in B1. Short streams and bundles of highly pleomorphic, spindle-shaped to polygonal neoplastic mesenchymal cells in a minimal stroma, with numerous multinucleate cells. **(B3 and B4)** Positive cytoplasmic desmin and nuclear myogenin immunohistochemistry confirming that the neoplastic cells originate from skeletal muscle. **(C1)** A male *Trp53*^R210X/+^ mouse with numerous severely enlarged peripheral lymph nodes (Submand L = submandibular lymph nodes; Ax L = axillar lymph nodes; Ing L = inguinal lymph nodes). Inset shows lungs *in situ.* **(C2 and C3)** Histomicrographs of a lymph node and the spleen from the animal shown in C1, with dense proliferation of neoplastic round cells variably effacing the normal tissue architecture i.e. lymphoma. **(C4-C6)** Histomicrographs of lung, liver and kidney from the animal shown in C1, with perivascular and interstitial neoplastic infiltrates, i.e. metastatic lymphoma (**C4**: Alv = alveolus, V = blood vessel lumen, L = lung parenchyma; **C5**: Li = liver parenchyma; **C6**: Ki = kidney parenchyma). **(D)** Distribution and combinations of tumors and multicentric or metastasizing tumors found in *Trp53*^R210X/+^ mice. The total number of mice includes all mice with at least one tumor. To increase clarity, pleomorphic lymphoma and histiocytic sarcoma were only included when they were the only diagnosed tumor(s), not when they occurred concurrently with other tumor types. Therefore, the number of multicentric tumors are lower than in Supplementary Table S3. The full data is presented in Supplementary Tables S4B and S4D. Ad = adenoma; Ca = carcinoma; Hemangio = hemangiosarcoma; HS = histiocytic sarcoma; Leiomyo = leiomyosarcoma; Lymph = lymphoma; mc = multicentric; Mesothel = mesothelioma; met = metastatic; Osteo = osteosarcoma; Pleo lymph = pleomorphic lymphoma; Rhabdo = rhabdomyosarcoma; Sarc = sarcoma. **(E)** Representative agarose gel for evaluation of LOH in tumors collected from four different euthanized mice. Each animal’s tail was used as a control tissue. WT (upper panel) and R210X (lower panel) gel band intensities were measured to quantify the percentage of WT allele retained in each tumor compared to the tail; percentage of retained WT allele in each sample is indicated at the bottom of the panel. LOH was defined as <50% WT allele retained. N = no LOH; Y = LOH.

T-cell lymphomas, confirmed by positive CD3 staining, were only found in homozygous (66.7 %) and heterozygous (5.1 %) mice (Fig. 5A; Supplementary Fig. S2A-B, S3A; Supplementary Table S4A-D). Only *Trp53*^R210X/R210X^ mice had solitary thymic T-cell lymphoma, but the most common type of lymphoma in homozygous mice was multicentric thymic T-cell lymphoma, which was less common in *Trp53*^R210X/+^ mice and absent in WT mice (Fig. 5B; Supplementary Table S4A-C). In *Trp53*^R210X/+^ and in WT mice the most common type of lymphoma was multicentric pleomorphic lymphoma. Pleomorphic lymphomas, confirmed by immunohistochemistry (IHC) (Supplementary Table S4A-C) were most frequent in WT mice and lowest in homozygous mice (adj. P-value: *p* = 0.0048) (Fig. 5A).

**Figure 5.**
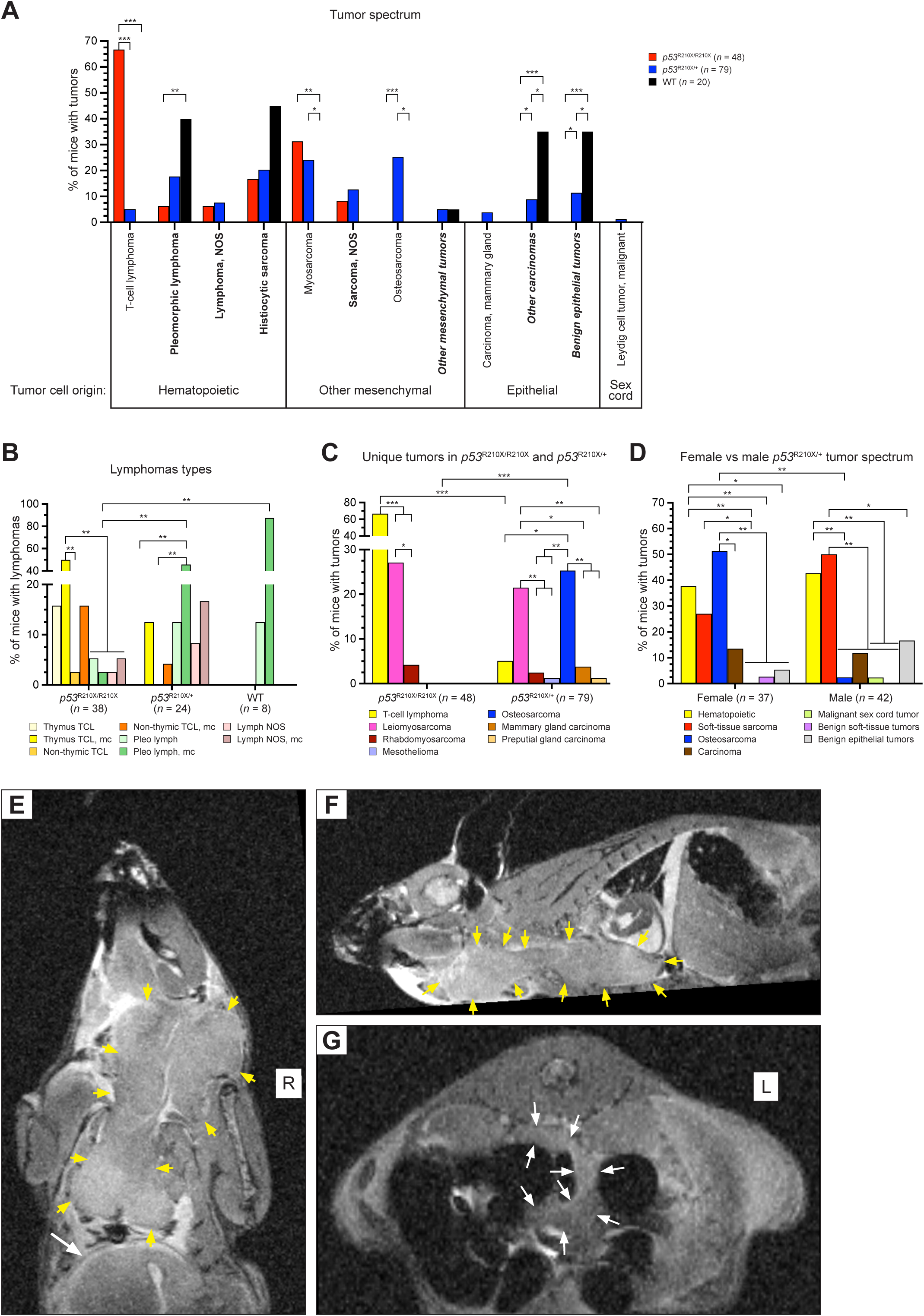
Comparison of tumor spectra of *Trp53*^R210X/R210X^, *Trp53*^R210X/+^ and WT mice. **(A)** Most frequently found tumor types of hematopoietic, mesenchymal, epithelial and sex cord origin in the three genotypes. *Trp53*^R210X/R210X^ mice showed a high rate of T-cell lymphoma whereas *Trp53*^R210X/+^ was the only genotype that developed osteosarcomas and malignant sex cord tumors. Only *Trp53*^R210X/R210X^ and *Trp53*^R210X/-^ developed soft-tissue sarcomas. Tumor type in bold = found in C57BL/6J WT (i.e.: pleomorphic lymphoma; lymphoma, NOS; histiocytic sarcoma; sarcoma, NOS; other mesenchymal tumors; other carcinomas; benign epithelial tumors) (36); tumor type in italics = several tumor types grouped together; full tumor spectrum is shown in Supplementary Fig. S5A. **(B)** Predominant lymphoma subtypes in *Trp53*^R210X/R210X^, *Trp53*^R210X/+^ and WT mice. TCL = T-cell lymphoma; mc = multicentric; NOS = not otherwise specified; Pleo lymph = pleomorphic lymphoma. *Trp53*^R210X/R210X^ had frequent solitary and multicentric thymic T-cell lymphoma while *Trp53*^R210X/+^ and WT mice mainly developed solitary and multicentric pleomorphic lymphomas. **(C)** Distribution of the tumor types unique to the *Trp53*^R210X^ mice, i.e. never found in the WT mice, and never or rarely reported as spontaneous tumors in C57BL6/J mice (36). T-cell lymphoma was more frequent in *Trp53*^R210X/R210X^ than in *Trp53*^R210X/+^ mice, and osteosarcoma was more frequent in *Trp53*^R210X/+^ mice. In both genotypes, the most common myosarcoma was leiomyosarcoma. In *Trp53*^R210X/+^ mice, leiomyosarcomas were as common as osteosarcomas, and both these tumors were more common than all other unique tumors. **(D)** Comparison of tumor spectra in female and male *Trp53*^R210X/+^ mice. Female *Trp53*^R210X/+^ had more frequent osteosarcomas than males. Comparisons of tumor spectra in female and male *Trp53*^R210X/R210X^ and WT mice are shown in Supplementary Fig. S5B-C. **(E-G)** A male *Trp53*^R210X/R210X^ mouse was sacrificed at 234 days of age due to weight loss, strained breath and a neoplasia on the throat, and subjected to an *ex vivo* MRI analysis without prior necropsy. **(E and F)** The throat neoplasia was part of a large neoplastic thymus (outlined by yellow arrows, size approx. 2.7x1.3x0.9 cm) that intrathoracally extended caudally to the diaphragm (indicated by a white arrow in **E**), cranially through the thoracic inlet along the ventral neck to the level of the ramus mandibularis, and extending into the right-side musculature medial to the shoulder, probably involving the prescapular lymph node. In the right cranial mediastinum there was a neoplastically transformed cranial mediastinal lymph node, about 0.8 cm long and 0.4 cm in diameter. The submandibular lymph nodes were not identifiable and appeared to be continuous with the thymus tumor. The lungs had a clear appearance with no signs of neoplastic proliferations or infiltrates. **(G)** In the dorsal abdomen there were multiple enlarged lymph nodes, up to 3 mm in diameter, elongated and convoluted (outlined by white arrows). The spleen was mildly enlarged and rounded (not visible in the figure), and in the normal-sized liver there were two nodules with an appearance best consistent with areas of vacuolar degeneration and extramedullary hyperplasia. Statistical analysis in **A-D** was performed by two-sided Fisher’s exact test, followed by correction for multiple comparisons using the Holm-Šídák method. In **A**, comparisons were made between genotypes within each tumor type. In **B-C**, comparisons were made between all tumor types within one genotype, and between the different genotypes for each individual tumor type. In **D**, comparisons were made between female and male for each tumor type (i.e. female osteosarcoma vs. male osteosarcoma), and between all tumor types within one sex (i.e. all tumor types found in females). Adjusted P-values: \**p* < 0.05, \*\**p* < 0.01; \*\*\**p* < 0.001.

Similarly, malignant soft-tissue sarcomas were only found in homozygous and heterozygous mice (Fig. 5A and C; Supplementary Fig. S5A; Supplementary Table S4A-C). Of the other mesenchymal tumor types, osteosarcoma was only found in heterozygous mice (25.3 %), and both genotypes had similar proportions of soft-tissue sarcomas (Fig. 5A and C; Supplementary Fig. S5A; Supplementary Table S4A-B). *Trp53*^R210X/R210X^ mice developed primary soft-tissue sarcomas in subcutis, skin, and skeletal muscle (Fig. 3; Supplementary Fig. S2C; Supplementary Table S4A), whereas *Trp53*^R210X/+^ mice developed soft-tissue sarcomas in these sites as well as in numerous other organs, including urogenitalia (Fig. 4; Supplementary Fig. S3B; Supplementary Table S4B). The majority of soft-tissue sarcomas were myosarcomas – primarily leiomyosarcomas that were confirmed by positive desmin staining and negative myogenin staining, and rhabdomyosarcomas confirmed by positive desmin and negative myogenin staining (Fig. 3 and 4; Supplementary Fig. S2C, S3B, S5A; Supplementary Table S4A-B). Female *Trp53*^R210X/+^ had more frequent osteosarcomas than males (adj. P-value: *p* = 0.0028) (Fig. 5D). *Trp53*^R210X/R210X^ and WT mice showed no significant differences between sexes (Supplementary Fig. S5B-C).

To visualize the *in situ* distribution of lymphoma in a *Trp53*^R210X/R210X^ mouse, we performed post-mortem MRI on a 234-days old male that was euthanized due to a large tumor on the ventral neck and respiratory distress. The MRI showed extensive neoplasias originating and extending from the thymus and several lymph nodes (Fig. 5E-G).

### Frequent multicentric/metastatic tumors with variable impact on survival

Nearly two-thirds of the *Trp53*^R210X/R210X^ mice (29 of 48; 60%) had multicentric or metastatic tumors (Supplementary Tables S3 and S4A). Of these, 25 were T-cell lymphomas (of 32 T-cell lymphomas in total), three were other hematopoietic tumors, and one a soft-tissue sarcoma. The most common lymphoma sites were thymus, spleen, lung and liver, followed by kidney, heart, and submandibular salivary gland (Fig. 3A-B; Supplementary Fig. S2A-B; Supplementary Table S4A). In comparison, only a little more than one-third of the heterozygous mice (29 of 79; 37%) had multicentric or metastatic tumors (Supplementary Tables S3 and S4B). Twenty-two were hematopoietic tumors, three were soft-tissue sarcomas, three were osteosarcomas, and one a carcinoma. Half of the tumor-bearing WT mice (10 of 20, 50%) had multicentric or metastatic tumors, and all these were hematopoietic tumors (Supplementary Tables S3 and S4C). The number of organs and tissues with neoplasia are probably underestimated, since not all tissues were sampled from all animals.

There was no significant effect of multicentric or metastatic disease on survival of *Trp53*^R210X/R210X^ mice (*p* = 0.0595), whereas *Trp53*^R210X/+^ mice with multicentric or metastatic disease had significantly longer survival than mice with only solitary tumors (*p* = 0.0167) (Fig. 6A). To elucidate if tumor type was prognostic of survival in mice with solitary malignant tumors, we extracted survival data for the most common tumor type in each genotype and compared to the remaining mice with solitary tumors. We found no differences in survival between solitary T-cell lymphomas and other malignant solitary tumors in homozygous mice, or between solitary osteosarcomas and other malignant, solitary tumors in heterozygous mice (Fig. 6B). However, *Trp53*^R210X/R210X^ mice with solitary T-cell lymphoma, but not *Trp53*^R210X/R210X^ mice with solitary soft-tissue sarcomas, had significantly shorter survival than mice with multicentric or metastatic tumors (Fig. 6C). Furthermore, *Trp53*^R210X/R210X^ mice with thymic lymphoma showed decreased survival compared to those with both thymic and splenic lymphoma (Fig. 6D). Since thymic tumors can interfere with heart and lung function, we also investigated thymic tumor size. *Trp53*^R210X/R210X^ mice with only thymic lymphoma had the largest thymus tumors compared to mice with splenic lymphoma or both thymic and splenic lymphoma (Fig. 6E).

**Figure 6.**
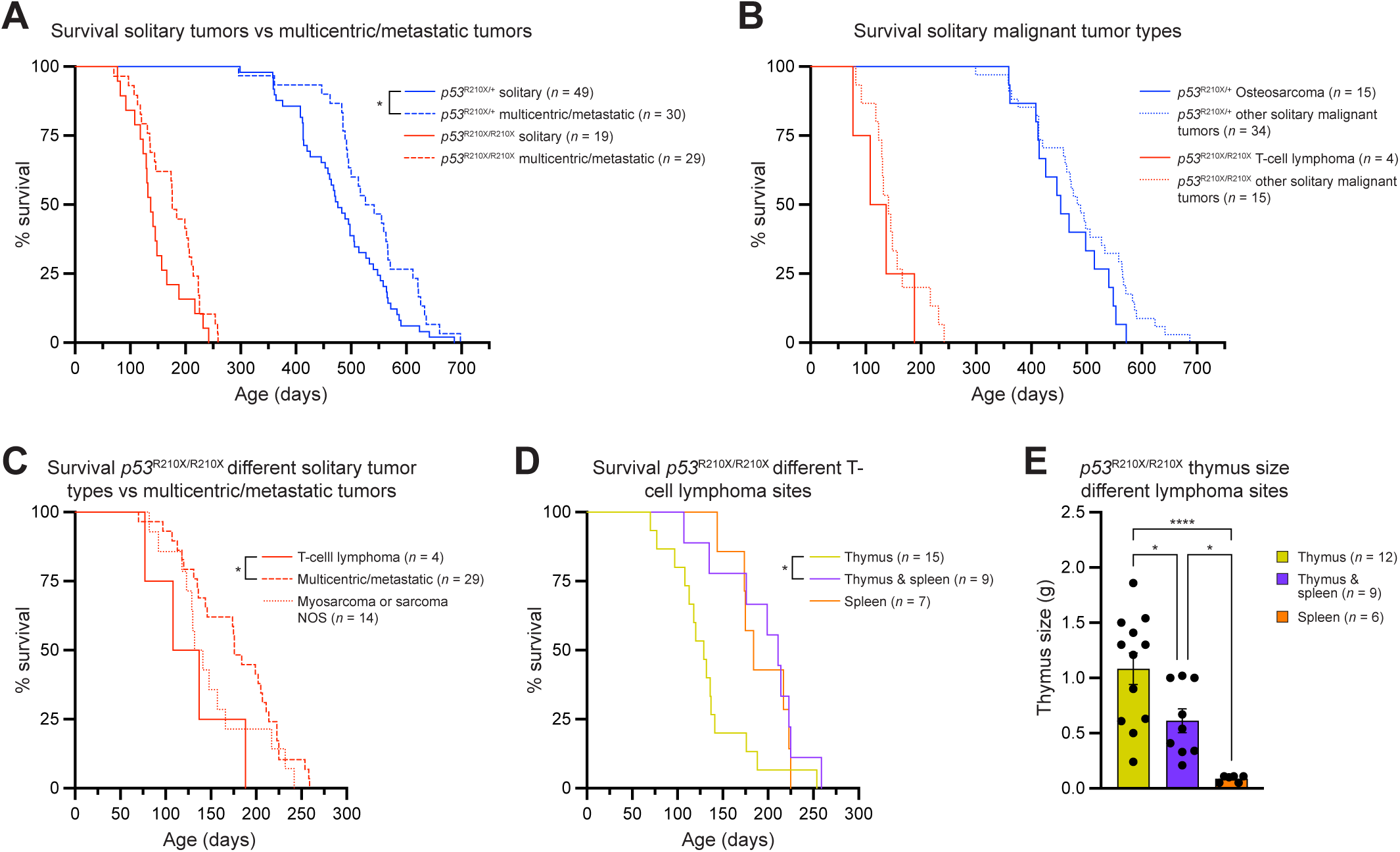
Impact of multicentricity, metastasis and thymus size on overall survival of *Trp53*^R210X/R210X^ and *Trp53*^R210X/+^ mice. **(A)** Kaplan-Meier curves showing that *Trp53*^R210X/+^ mice with multicentric or metastatic disease had significantly better survival compared to mice with solitary tumors (\**p* = 0.0167), whereas *Trp53*^R210X/R210X^ mice did not (*p* = 0.0595). **(B)** *Trp53*^R210X/R210X^ and *Trp53*^R210X/+^ mice with the indicated malignant solitary tumors (most common solitary tumor types for respective strain) did not show any differences in survival compared to mice of the same genotype with other solitary tumors. Note that two *Trp53*^R210X/R210X^ mice that were euthanized due to solitary leiomyosarcoma also had solitary thymic T-cell lymphoma and were included as solitary thymic T-cell lymphoma in Fig. 5B. Here, however, they are included as “other solitary malignant tumors” since that was the cause for euthanasia. **(C)**. *Trp53*^R210X/R210X^ mice with solitary T-cell lymphomas showed significantly shorter survival than *Trp53*^R210X/R210X^ mice with multicentric or metastatic disease (*p* = 0.0424) but not compared to mice with sarcomas. Note that two *Trp53*^R210X/R210X^ mice that were euthanised due to solitary leiomyosarcoma also had solitary thymic T-cell lymphoma and were included as solitary thymic T-cell lymphoma in Fig. 5B. Here, however, they are included as “myosarcoma or sarcoma NOS” since that was the cause for euthanasia. **(D)** *Trp53*^R210X/R210X^ mice with thymic T-cell lymphoma had a shorter overall survival compared to *Trp53*^R210X/R210X^ mice with T-cell lymphoma in both thymus and spleen (adj. P-value: *p* = 0.038). **(E)** The thymi of *Trp53*^R210X/R210X^ mice with solitary thymic T-cell lymphoma were in general larger than the thymi from *Trp53*^R210X/R210X^ mice with T-cell lymphoma in the spleen or in both thymus and spleen. Mean ± SEM are indicated. Survival curves in **A-D** were analyzed by log-rank (Mantel-Cox) test followed by correction for multiple comparisons using the Holm-Šídák method when applicable. In **D,** comparisons were made to the group with solitary thymic lymphoma. In **E**, statistical analysis was performed using One-way ANOVA followed by Tukey’s multiple comparisons tests. P-values: \**p* < 0.05, \*\**p* < 0.01; \*\*\**p* < 0.001.

### Translational readthrough in T-cell lymphoma cells derived from *Trp53*^R210X/R210X^ mice

To investigate if translational readthrough of R210X nonsense mutant *Trp53* can generate full-length active p53, we established T-cell lymphoma lines from thymic lymphomas of *Trp53*^R210X/R210X^ mice (X405, X491 and X547). As controls, we used T-cell lymphoma lines established from thymic lymphomas of *Trp53*^R172H/R172H^ mice (H661 and H671). Fig. 7A shows HE and CD3 IHC staining of thymus from mouse X405, verifying T-cell origin of the tumor. Supplementary Fig. S6A shows corresponding histological images for thymi of *Trp53*^R210X/R210X^ mice X491 and X547 as well as *Trp53*^R172H/R172H^ mice H661 and H671. All cell lines were analyzed with PCR to confirm the correct genotype (Supplementary Fig. S6B). Flow cytometry with CD3, CD4 and CD8 antibodies verified T-cell origin, with 97.5% CD3^+^ cells and >99% among those CD4^+^CD8^+^ (Fig. 7B). To examine translational readthrough, we treated T-cell lymphoma cells with aminoglycoside G418 for 72 h. Immunofluorescence staining with anti-p53 C-terminal antibody (280aa C-term) that detects full-length p53 but not C-terminally truncated p53, and anti-p53 N-terminal antibody N1 that detects both full-length and C-terminally truncated p53, showed that 100µM G418 induced full-length p53 (Fig. 7C), and that treatment with 200µM G418 caused substantial cell death (Supplementary Fig. S6C). Western blotting with anti-p53 antibody 1C12 that recognizes an N-terminal epitope confirmed a dose-dependent induction of translational readthrough and expression of full-length p53 (Fig. 7D; Supplementary Fig. S7A). We also found a slight increase in truncated p53, most likely due to inhibition of nonsense-mediated decay following translational readthrough (30). Moreover, qRT-PCR revealed a dose-dependent upregulation of p53 target genes *Puma* and *Zmat3* and to a lesser extent *Cdkn1a* (*p21*) and *Bax* in all *Trp53*^R210X/R210X^ T-lymphoma cell lines but not in T-lymphoma cell lines derived from *Trp53*^R172H/R172H^ mice upon treatment with G418 (Fig. 7E; Supplementary Fig. S7B), confirming that full-length p53 induced by G418 is functional as transcription factor. G418 also inhibited cell proliferation in a dose-dependent manner in T-lymphoma cells derived from both *Trp53*^R210X/R210X^ and *Trp53*^R172H/R172H^ mice as shown by the WST-1 assay, but with significantly higher efficiency in the *Trp53*^R210X/R210X^-derived T-lymphoma lines (Fig. 7F; Supplementary Fig. S7C). Finally, G418 at 150 and 200µM induced apoptotic cell death in the T-lymphoma cells as shown by Annexin V flow cytometry (Fig. 7G; Supplementary Fig. S8). This cell death was significantly more efficient in the *Trp53*^R210X/R210X^ T-lymphoma cells than in the *Trp53*^R172H/R172H^ T-lymphoma cells at 200µM G418.

**Figure 7.**
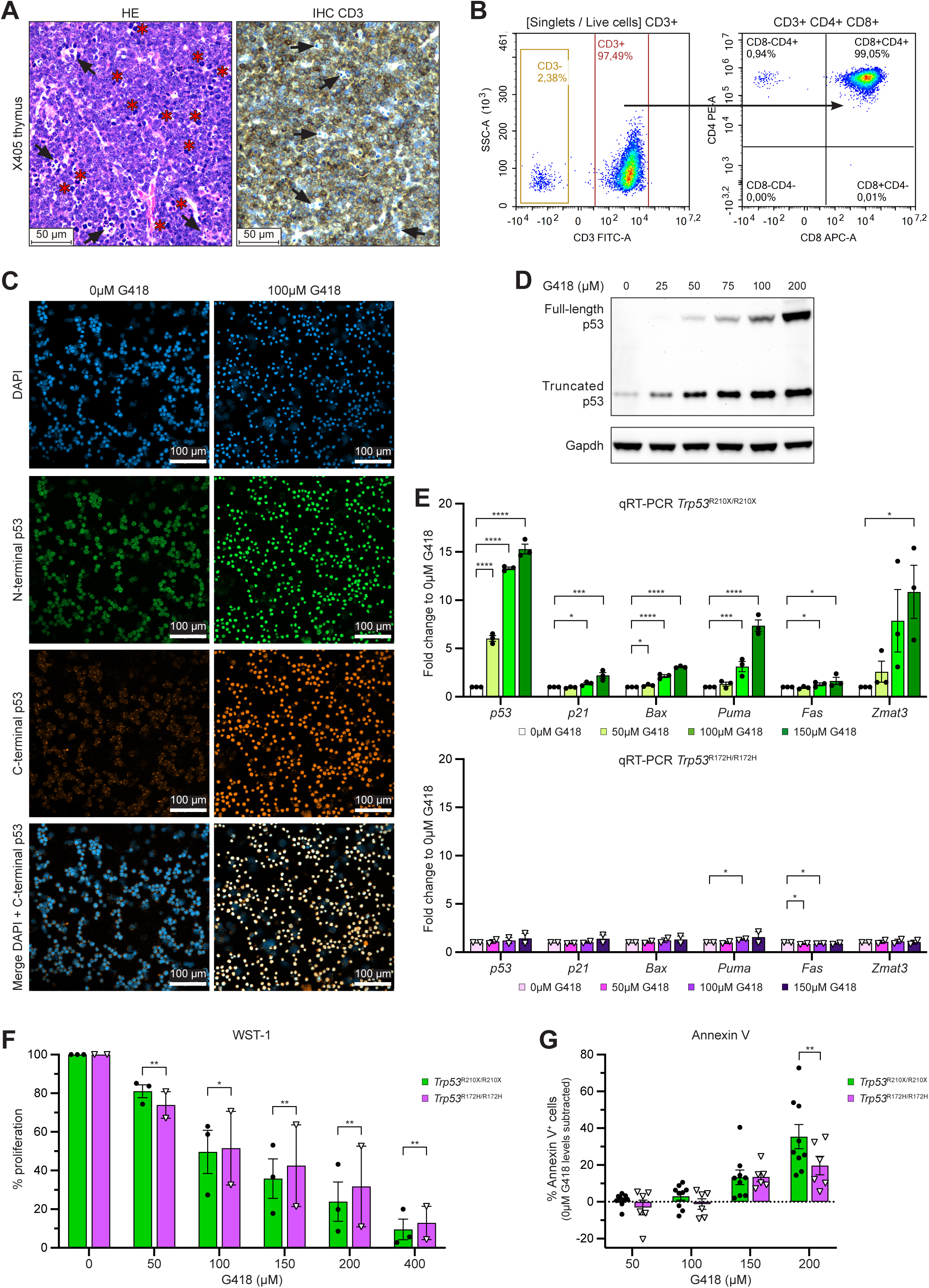
G418 induces translational readthrough and full-length functional p53 in *Trp53*^R210X/R210X^ T-lymphoma cells. **(A)** Representative histomicrographs of the thymus from the *Trp53*^R210X/R210X^ mouse X405 stained with HE (left) and CD3 antibody (IHC, right). Mats of CD3-positive, i.e. T-cell origin, pleomorphic neoplastic lymphocytes with numerous mitoses (asterisks), and dispersed, moderate numbers of tingible body macrophages (arrows) efface the normal thymic architecture. Scale bars = 50µm. **(B)** Flow cytometry to verify T-cell origin of X405 cells derived from a *Trp53*^R210X/R210X^ thymic lymphoma by staining with antibodies for CD3 (left panel), and CD4 and CD8 (right panel). **(C)** Immunofluorescence staining of X405 T-lymphoma cells, either untreated (0µM G418, left panels) or treated with 100µM G418 (right panels) for 72 hours. p53 was detected using N1 (N-terminal epitope) and 280aa C-term (C-terminal epitope) antibodies. Bottom panels show merged DAPI and 280aa C-term p53 staining, highlighting translational readthrough of the *Trp53*-R210X nonsense mutation and production of full-length p53 protein upon treatment with G418. Scale bars = 100µm. **(D)** Western blot analysis showing dose-dependent induction of full-length p53 in X405 T-lymphoma cells following 72h treatment with G418 at indicated concentrations. p53 was visualized using the anti-p53 antibody 1C12 that recognizes an N-terminal epitope and therefore detects both full-length and C-terminally truncated p53. Gapdh was used as a loading control. **(E)** qRT-PCR analysis showing dose-dependent induction of *p53* mRNA and p53 target gene mRNA levels in the three *Trp53*^R210X/R210X^ T-lymphoma cell lines X405, X491 and X547 after 72h treatment with G418 (upper panel). Two independently established *Trp53*^R172H/R172H^ T-lymphoma cell lines were used as negative control (lower panel). Gene expression values were normalized to *Gapdh* expression and compared to the untreated (0µM G418) negative control for each gene. Each dot represents the mean from three independent experiments per cell line. **(F)** WST-1 assay to determine proliferation of T-lymphoma cells from *Trp53*^R210X/R210X^ (*n* = 3 cell lines) and *Trp53*^R172H/R172H^ (*n* = 2 cell lines) mice upon 72h treatment with indicated concentrations of G418. Each dot represents the mean from eight independent experiments per cell line. **(G)** Flow cytometry of Annexin V staining to assess apoptotic cell death in *Trp53*^R210X/R210X^ (*n* = 3 cell lines) and *Trp53*^R172H/R172H^ (*n* = 2 cell lines) T-lymphoma cells following 72h treatment with G418. All quantifications from three independent experiments per cell line are shown. Statistical analysis was performed by repeated measures two-way ANOVA followed by Dunnett’s multiple comparisons test **(E)**, and by two-way Mixed-effects analysis **(F**-**G)**, respectively. All individual cell line values from all experiments were used for statistical analysis but mean ± SEM are indicated in figures **E** and **F**. Adjusted P-values: \**p* < 0.05, \*\**p* < 0.01, \*\*\**p* < 0.001, \*\*\*\**p* < 0.0001.

## Discussion

We present here the first mouse model carrying a *Trp53* nonsense mutation. *Trp53*-R210X corresponds to human *TP53*-R213X, one of the 10 most common *TP53* mutations in human tumors. The *Trp53*^R210X^ allele was transferred at Mendelian frequency in all crosses except intercrosses with female *Trp53*^R210X/R210X^ mice that resulted in smaller litters than the reference number for the C57BL/6J WT background strain. These results are consistent with data from *Trp53*-null mice, where significant decreases in embryonic implantation, pregnancy rate and litter size were observed in matings with female *Trp53*^-/-^ mice (31). This effect in *Trp53*-null mice is most likely caused by reduced levels of leukemia inhibitory factor (LIF), which may also be pertinent to our new *Trp53*^R210X^ strain.

We observed a substantial reduction in the frequency of female *Trp53*^R210X/R210X^ pups regardless of breeding setup, consistent with previous data for *Trp53*-null mice (32–35). Exencephaly and craniofacial malformations in *Trp53*^-/-^ female embryos (32, 33, 35) may also explain the skewed gender distribution observed in the *Trp53*^R210X/R210X^ mice. Apart from the poor breeding capacity of female *Trp53*^R210X/R210X^ mice, heterozygous and homozygous mice of both genders appeared phenotypically normal initially and were fertile, as reported for *Trp53*-null mice (15, 16).

The reduced bodyweights of female *Trp53*^R210X/R210X^ mice compared to *Trp53*^R210X/+^ and WT mice was unexpected. To our knowledge, similar findings have not been reported for any genetically manipulated *Trp53*-model; in one study, female and male *Trp53*^+/-^ and *Trp53*^-/-^ mice showed no difference in bodyweight (34), and two recently published *Trp53* missense knock-in mouse models displayed normal male and female bodyweights (19). In our study, male mice of all three genotypes had almost identical weight curves up until 34 weeks of age, very similar to those reported for male *Trp53*-null mice (34). At that point, the weight curve for male *Trp53*^R210X/R210X^ mice began to deviate downward from those of *Trp53*^R210X/+^ and WT mice, presumably a direct consequence of tumor development.

*Trp53*^R210X/R210X^ mice had similar survival curves as the control *Trp53*^R172H/R172H^ mice, as well as several other published *Trp53*-null and *Trp*53^Mis^ mouse models (9, 15–20, 34). Interestingly, *Trp53*^R210X/+^ mice showed significantly longer survival than the *Trp53*^R172H/+^ mice in our colony. One plausible explanation for this difference is that the truncated p53 protein encoded by the *Trp53*^R210X^ allele exerts a less pronounced dominant-negative effect by oligomerization with co-expressed WT p53 as compared to R172H missense mutant p53, or completely lacks such effect, allowing more potent WT p53-dependent tumor suppression in *Trp53*^R210X/+^ mice.

As reported for homozygous *Trp53*-null and *Trp53* missense knock-in mice (9, 15–20), homozygous *Trp53*^R210X^ mice showed a remarkable susceptibility to tumor development at an early age, with lymphoma as the predominant tumor type, (79.2% in *Trp53*^R210X/R210X^ mice and 57-78% in homozygous *Trp53*-null and *Trp53*^Mis^ mice) followed by soft-tissue sarcoma (35.4% in *Trp53*^R210X/R210X^ mice and 22-38% in homozygous *Trp53*-null and *Trp53*^Mis^ mice). This confirms that *Trp53* nonsense mutation at codon 210 leads to a dramatically increased tumor incidence, just like complete loss or missense mutation of *Trp53*. The most common lymphoma type in *Trp53*^R210X/R210X^ mice was T-cell lymphoma (84% of lymphoma cases). T-cell lymphomas and soft-tissue sarcomas can be considered as *de novo* tumor types, since they were never found in the WT mice, and are very rare in the background C57BL6/J strain (36). In contrast, pleomorphic lymphoma and histiocytic sarcoma, which were found in 20.8% of the homozygous mice, are common tumor types that are found at all ages, although at increasing incidence with age, and cannot therefore be reliably associated with the homozygous genotype (37, 38).

Whereas all homozygous *Trp53*^R210X^ mice had died or been euthanized by 8.5 months of age, heterozygosity for the *Trp53*^R210X^ mutation did not begin to affect survival until approximately 9 months of age. Like homozygous *Trp53*^R210X^ mice, heterozygous mice showed a striking tumor phenotype, although with larger variation in tumor type and later onset of clinical signs compared to homozygous mice, and with frequent LOH, again in agreement with previous studies of *Trp53*-null and *Trp53* missense knock-in mouse models (9, 15–17, 20, 39). Heterozygous mice also developed the *de novo* soft-tissue sarcomas, and more rarely T-cell lymphoma, as well as osteosarcomas, mammary gland carcinomas, and in one case a malignant Leydig cell tumor, none of which are reported as a spontaneous tumor in C57BL/6J mice (36, 40). The malignant Leydig cell tumor is of particular interest since its presence shows that this mouse model also can develop tumors of sex cord origin, i.e. testicular or ovarian, in addition to the more frequent hematopoietic, mesenchymal and epithelial tumors. Thus, the *Trp53*^R210X/+^ mice, like previously published *Trp53*-null and missense knock-in models, represent a valuable model for the Li-Fraumeni syndrome (LFS), in which *TP53* nonsense mutations occur with a similar frequency as in somatic tumors according to The *TP53* Database collection of germline variants (The *TP53* Database (R21, Jan 2025): https://tp53.cancer.gov) (41). The delayed tumor onset in *Trp53*^R210X/+^ compared to *Trp53*^R210X/R210X^ mice reflects the need to eliminate the remaining WT allele. This provides time for acquisition of additional genetic alterations that might predispose to other tumor types than those reported in C57BL/6J mice (36, 40).

*Trp53*^R210X/R210X^ mice frequently developed multicentric lymphomas, most commonly with tumors in liver and lung, and one leiomyosarcoma metastasis, whereas the *Trp53*^R210X/+^ mice showed a comparatively lower rate of multicentric lymphoma and higher rate of soft-tissue sarcoma metastasis. Unexpectedly, multicentric or metastatic disease did not have a negative effect on survival of *Trp53*^R210X^ mice. H*o*wever, it should be noted that solitary tumors without metastasis can be life-threatening through various mechanisms, such as rapid growth that compromises organ function, e.g. compression and restriction of heart and lungs by thymic lymphomas, or production of substances that affect vital organs or have detrimental systemic effects.

The observed high rate of lymphoma multicentricity in *Trp53*^R210X^ mice is in contrast to data for *Trp53*^R246S/R246S^, *Trp53*^R245W/R245W^ and *Trp53*^R270H/R270H^ mice that only infrequently or never develop multicentric or metastatic disease (18, 20). This could suggest that truncated p53 derived from the R210X nonsense mutant allele can exert some kind of GOF activity. Indeed, GOF features of C-terminally truncated p53 proteins have been demonstrated in cell culture and *in vivo* assays (14). However, we did not observe any significant differences in overall survival, tumor incidence, tumor spectrum or LOH frequency between our *Trp53*^R210X^ mice and data reported for *Trp53*-null mice (15, 16, 39), arguing against a prominent GOF activity of the R210X allele in this setting. A caveat is that specific data on multicentricity or metastasis frequency in homozygous *Trp53*-null or missense knock-in mice are limited in previous studies (9, 15–20). Thus, we do not exclude that further investigation will reveal GOF properties of nonsense mutant *TP53*.

To validate the utility of our model as a platform for the development of nonsense mutant *TP53*-targeted anti-cancer drugs, we showed that the well-known readthrough inducer G418 could restore expression of full-length and functional p53 in T-lymphoma cell lines derived from *Trp53*^R210X/R210X^ mice. We observed upregulation of p53 target genes in the T-lymphoma cell lines derived from *Trp53*^R210X/R210X^ mice but not in *Trp53*^R172H/R172H^ T-lymphoma control cell lines. The observation that G418 inhibited cell growth and induced apoptotic cell death with statistically significant higher efficiency in the *Trp53*^R210X/R210X^ lymphoma cells than in the *Trp53*^R172H/R172H^ lymphoma cells confirms a p53-dependent biological effect, although G418 also has p53-independent general toxicity.

In conclusion, our *Trp*53^R210X^ mouse model should facilitate the development of novel strategies for therapeutic targeting of nonsense mutant *TP53* in cancer. Importantly, we have demonstrated that induction of translational readthrough of the *Trp53-*R210X allele in mouse thymic lymphoma cells results in the expression of full-length and functional p53 protein and cell death by apoptosis. Further studies of downstream effects of p53 activation in these cells are required, and it will also be interesting to examine translational readthrough and p53 activation in cell lines derived from other types of tumors in *Trp53*^R210X^ mice. Induction of translational readthrough is a potential therapeutic strategy for nonsense mutant *TP53*-carrying tumors upon diagnosis, or for prevention of tumors in cancer-prone individuals such as LFS family members carrying an inherited *TP53* nonsense mutation. Given that around 11% of all *TP53* mutations in cancer are nonsense mutations (6), successful clinical implementation of efficient readthrough-inducing therapy would have a significant impact on public health worldwide.

## Materials and methods

### Generation of *Trp53*^R210X^ mutant mice

To generate the *Trp53*^R210X^ mouse model, a chemically modified synthetic single-guide RNA (sgRNA) (5’-gcagacttttCGCcacagcg-3’) with an 80-mer SpCas9 scaffold targeting the region encoding the R210 codon in *Trp53* exon 6 was used (CRISPRevolution sgRNA EZ Kit, Synthego, USA). Single strand oligodeoxynucleotides (ssODNs) of 200 nucleotides were used as repair template (Ultramer^TM^, Integrated DNA Technologies, USA; 5’- ctgacttattcttgctcttaggcctggctcctccccagcatcttatccgggtggaaggaaatttgtatcccgagtatctAgaagacaggcagacttttTgAcaTagTgtggtggtaccttatgagccacccgaggtctgtaattttgttttggtttgtgcgtcttagagacagttgactccagcctagactgatgttgac-3’) for homology-directed repair (HDR). The sequence included silent mutations to exclude recutting of the guide RNA and an XbaI site (tctAga) close to the R210X codon TgA to facilitate genotyping. The CRISPR/Cas9 strategy was verified by editing NIH/3T3 mouse fibroblasts (CRL-1658, ATCC, USA), see below for genotyping strategy using the XbaI site. The edited NIH/3T3 cells were plated as single cells and cultured in Dulbecco’s Modified Eagle’s Medium High Glucose (DMEM) supplemented with 5% FBS.

The edited mice were produced at the Karolinska Center for Transgene Technologies, Karolinska Institutet. Zygotes harvested from superovulated and mated C57BL/6J female mice (Charles River Laboratories, Germany) were microinjected into the pronuclei with injection buffer [10 mM Tris-HCl (pH 7.5), 0.1 mM EDTA] containing a pre-formed Cas9-sgRNA ribonucleoprotein complex at a concentration of 20 ng/µl eSpCas9 (Sigma-Aldrich/Merck, Germany), 20 ng/µl of the *Trp53* sgRNA (Synthego), and 20 ng/µl of the 200 nucleotide ssODN Ultramer (Integrated DNA Technologies). Microinjected zygotes were implanted into oviducts of pseudopregnant SOPF RjOrl:SWISS (Janvier Labs, France) surrogate female mice via microsurgery. Ear punch biopsies from the F0 offspring were lysed (42) and subjected to PCR with primers encompassing the edited region, giving a 1049 bp amplicon (*Trp53* exon 5-7; Forward primer: 5’-ATCGTTACTCGGCTTGTCCC-3’; Reverse primer: 5’- GGGTAGGAACCAAAGAGCGT-3’). PCR products were purified using the QIAquick PCR Purification Kit (Qiagen, Germany) and digested with restriction enzyme XbaI (New England Biolabs, USA) to reveal recombination of the ssODN template. The PCR amplicons from selected candidate F0 mice were subcloned into pMiniT 2.0 (New England Biolabs) and individual plasmid clones were subjected to Sanger sequencing (KIGene, Karolinska Institutet). All F1 mice identified as heterozygous were further confirmed by Sanger sequencing individual plasmid clones using primers described above for *Trp53* exon 5-7. Sequences were analyzed using the SnapGene (version 7.0.1) and Jalview (version 2.11.2.6) softwares.

### Animal studies

Animal care was in accordance with Swedish regulation and Karolinska Institutet guidelines. Experimental mice were housed in enriched individually ventilated cages in 1 hour dawn/dusk and 10h light/dark cycle, at 22°C ± 2°C and 50% ± 20% air humidity at a pathogen-free animal facility at Karolinska Institutet. Mice were supplied with irradiated CRM(P) mouse pellets (SAFE Diets, France) and drinking water *ad libitum*. All animal studies were approved by the Stockholm Regional Animal Ethical Committee, Sweden (Dnr 14188-2019).

*Trp53*^R210X/+^ mice were maintained on a C57BL/6J background by backcrossing to C57BL/6J mice purchased from Charles River Laboratories. Homozygous mice were generated by intercrossing *Trp53*^R210X/+^ and/or *Trp53*^R210X/R210X^ mice. *Trp53*^R172H^ (9) (C57BL/6J background), a kind gift from Dr. Guillermina Lozano, MD Anderson Cancer Center, Houston, were used as a reference strain. DNA for genotyping was extracted from ear biopsies using the Gentra Puregene Tissue Kit (Qiagen). Genotyping *Trp53*^R210X^ mice was performed by two separate touchdown PCR reactions (10 cycles of touchdown from 65°C to 61°C, followed by 25 cycles at 60°C – total 35 cycles) with specific primers to detect mutant and WT alleles, respectively (*Trp53*-R210X allele, 252 bp amplicon, forward primer: 5’-CATCTACAAGAAGTCACAGCAC-3’, reverse primer: 5’-GGTACCACCACACTATGTCA-3’; *Trp53*-R210 WT allele, 95 bp amplicon, forward primer: 5’-GCAGACTTTTCGCCACAGC-3’, reverse primer: 5’- CTGGAGTCAACTGTCTCTAAGAC-3’). PCR products were analyzed on 2% agarose gels using a Gel Doc EZ Imager system (Bio-Rad Laboratories, USA). Genotyping of *Trp53*^R172H^ mice was performed as described(9). Mice presenting a mutant allele band together with a WT allele band in the genotyping PCR were considered heterozygous, and mice showing only a mutant allele band and no WT band were considered homozygous. Mice from *Trp53*^R210X/+^ intercrosses that were negative for the R210X mutation are referred to as WT mice and were used as controls. *Trp53*^R210X^ mouse bodyweights were recorded weekly from postnatal week 2 and until moribund and euthanized. Homozygous mice were palpated weekly for tumor occurrence starting at 10 weeks of age, and heterozygous as well as WT littermate mice were palpated from 12 months of age.

### Tissue collection and processing

Moribund mice were euthanized by cervical dislocation and tissues were directly collected for further analysis. Tissue samples were also collected from mice found dead, unless there was too much autolysis. Tissues for molecular analysis were collected either by immediate snap freezing in liquid nitrogen and stored at -80°C, or in RNAlater™ Stabilizing Solution (Thermo Fisher Scientific, USA). Tissues collected for histological analysis were carefully dissected and fixed in 4% formaldehyde (buffered, pH 6.9, Sigma-Aldrich/Merck) at room temperature on a slow set rocking table for 24-48h, and subsequently transferred to 70% ethanol for storage at 4°C until further use. Skeletal tissues and calcified tumors were decalcified using MolDecal10 Ready to use Solution (pH 7.2-7.4, HistoLab Products, Sweden) for 3-4 weeks in 4°C on a slow set rocking table. Decalcification solution was changed every 3-5 days. For establishment of thymic lymphoma cell lines, enlarged thymuses from homozygous mice were fine dissected and collected in ice-cold PBS (see below).

### Histopathology and immunohistochemistry (IHC)

Formaldehyde-fixed tissues were processed in an automated tissue processing machine (LOGOS, Milestone Medical, Italy) for dehydration, followed by paraffin-embedding and microtome sectioning (4μm). Sections were collected on Superfrost™ Plus Adhesion Microscope Slides (Thermo Scientific, USA). For histological analysis, slides were stained with hematoxylin and eosin (HE) using a standard protocol. Verification of tumor cell origin was done by IHC following standard methods. Staining was performed with antibodies against CD3 (1:900, A0452, DAKO/Technologies, USA), desmin (1:2000, ab32362, Abcam, UK), myogenin (1:100, ab124800, Abcam) (all with EDTA pH 8.5 for antigen retrieval), CD68 (1:800, #97778, Cell Signaling Technology, USA), and vimentin (1:1000, ab92547, Abcam) (both with citrate buffer pH 6.0 for antigen retrieval), and incubation at 4°C overnight. Visualization was performed using ImmPress HRP and ImmPACT DAB kits (MP-7401 and SK-4205, Vector Laboratories, USA), followed by counterstaining in Mayer’s hematoxylin. HE and IHC slides were examined by light microscopy.

### *Ex vivo* MRI data acquisition

MRI data were acquired using a 9.4T MRI scanner (Varian, UK) with a horizontal bore. The scanner was controlled with software VnmrJ v4.0 Revision A. A gradient insert with an inner diameter of 12 cm (Varian, UK) and a millipede RF-coil with an inner diameter of 40 mm (Varian) were used. The scan program contained an overview scan, scanner alignment, diffusion weighted, T1-weighted (fat suppressed and Two-point Dixon) and T2-weighted (fat suppressed and FLAIR) scans. Overview and Two-point Dixon sequences were acquired in sagittal plane, the others in dorsal plane. Images were reviewed in collaboration with a European specialist in veterinary diagnostic imaging on a computer workstation using digital imaging and communications in medicine image viewing software (Horos software vers. 3.3.6, Horos project).

### Assessment of loss of heterozygosity (LOH)

*Trp53*^R210X/+^ tumors collected from euthanized mice were analyzed for LOH by a semi-quantitative method, using the aforementioned genotyping PCR but with only 29 cycles and 100ng DNA per sample. DNA was extracted as described above under Animal studies. For each tumor, tail tissue from the corresponding mouse was used as a control. To quantify the fraction of WT allele left in the tumor, agarose gel band intensities of WT and mutant PCR products were measured for tail and tumor tissue, respectively; analysis was performed using Image Lab Software (Mac Version 6.1, Bio-Rad Laboratories) of gel band signal intensities from gel picture raw data. Tumor intensity ratio was normalized to the respective tail using the following formula:

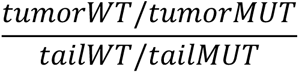

LOH was defined as a value < 0.5, indicating <50% WT allele fraction in tumor compared to tail.

### Establishment of T-cell lymphoma cell lines

Thymic (T-cell) lymphoma cell lines were established and cultured as described (43).

### Flow cytometry

T-cell origin of established cell lines was verified by flow cytometry. 5x10^5^ cells were incubated in staining buffer (3% FBS in PBS) containing the following antibodies: Anti-CD3-FITC (1:100, #11-0031-85), Anti-CD4-APC (1:100, #17-0041-83), and Anti-CD8-PE (1:100, #12-0081-85) (all eBioScience, USA). Alive cells were assessed by FxCycle^TM^ Violet Stain DAPI (Thermo Fisher Scientific) diluted 1:500 in staining buffer. Cells were incubated 20 min at 4^°^C with 100µl of the antibody/DAPI cocktail, followed by adding 1ml staining buffer and centrifugation at 400g for 5 min. Cells were resuspended in 500µl staining buffer, filtered through a 35μm cell strainer and analyzed using a NovoCyte flow cytometer (ACEA Biosciences, USA).

Cell death was assessed by Annexin V staining. 5x10^4^ cells/ml were seeded in 6-well-plates using 4ml/well. Cells were treated with G418 (0-200μM) for 72 hours. Treated cells were harvested, washed with DPBS (Sigma-Aldrich/Merck), resuspended in Annexin V binding buffer, counted using Countess 3 Automated Cell Counter (Thermo Fisher Scientific) and diluted to 1x10^6^ cells/ml. 100µl cell dilution was incubated with 5µl BD HorizonTM V450 Annexin V antibody (BD Biosciences, USA) for 15 minutes in the dark at room temperature (RT). 400µl Annexin V binding buffer was added and cells were analyzed using a NovoCyte flow cytometer (ACEA Biosciences).

### Western blotting

T-lymphoma cells were seeded in 6-well-plates at 1-3x10^5^ cells/ml and treated with G418 (0-200μM) for 72 hours. Western blotting was performed as previously described using 80μg protein (44). Detection of p53 was performed using primary antibody 1C12 anti-p53 (1:1000, #2524, Cell Signaling Technology) and an HRP-conjugated Rabbit-anti-Mouse IgG secondary antibody (1:5000, #61-6520, Invitrogen, USA). Following visualization of p53, HRP-conjugated G-9 anti-GAPDH antibody (1:10 000, sc-365062, Santa Cruz Biotechnology, USA) was used as a loading control. Proteins were visualized by SuperSignal™ West Femto Maximum Sensitivity Substrate and iBright FL1000 imaging system (both Thermo Fisher Scientific). All original Western blots are provided in Supplementary Information – Original Western blots.

### Immunofluorescence staining

1x10^5^ T-lymphoma cells in 100μl PBS were used for Cytospin (Shandon CYTOSPIN 4, Thermo Electron Corporation) at 1500 rpm for 1 min. Cells were fixed in 4% formaldehyde for 10 min at RT, permeabilized using 0.1% Triton in PBS for 5 min at RT, followed by washing in PBS. Primary antibodies rabbit polyclonal p53 N-terminal antibody [N1] (GTX100629, Genetex, USA) and goat polyclonal Anti-TP53 (280aa C-Term) (ASJ-VN4O74-150, Nordic Biosite, Sweden) were diluted together at 1:100 in PBS, incubation 4°C ON. Following PBS wash, secondary antibodies Donkey anti-Rabbit IgG (H+L) Alexa Fluor™ Plus 488 (A32790) and Donkey anti-Goat IgG (H+L) Alexa Fluor™ Plus 555 (A32816) (both Thermo Fisher Scientific) were diluted together at 1:1000 in PBS, incubation 1h in RT. Slides were washed with PBS and mounted with Fluoroshield™ containing DAPI (F6057, Sigma-Aldrich/Merck). An AxioObserver Z1 (Zeiss) microscope was used for imaging.

### Quantitative real-time PCR (qRT-PCR)

T-lymphoma cells were seeded in 6-well-plates at 5x10^4^ cells/ml and treated with G418 (0-150μM) for 72 hours. Cells were harvested and RNA extraction, cDNA synthesis and qRT-PCR were performed as described except that cDNA synthesis was carried out using SuperScript IV (Thermo Fisher Scientific) (27). TaqMan probes used were *p53* (*Trp53*; Mm01731290_g1), *p21* (*Cdkn1a*; Mm00432448_m1), *Bax* (Mm00432051_m1), *Puma* (*Bbc3*; Mm00519268_m1), *Fas* (Mm01204974_m1), *Zmat3* (Mm00494181_m1), and *Gapdh* (Mm99999915_g1).

### WST-1 assay

T-lymphoma cells were seeded in 6-well-plates at 5x10^4^ cells/ml and treated with G418 (0-400μM) for 72 hours. Cells (100µl) were then transferred to a 96-well-plate and 10µl of cell proliferation reagent WST-1 (Roche, Switzerland) were added. Absorbance was determined at 450nm with a Varioskan™ LUX multimode microplate reader (Thermo Scientific) after 30, 45 and 60 minutes incubation in the cell incubator.

### Statistical analysis

Statistical analysis of numerical data was performed using GraphPad Prism 10 (version 10.4.0). Multiple groups were compared using One-way ANOVA, followed by Tukey’s multiple comparisons test when all groups were compared to each other, or by Dunnett’s multiple comparisons test when all groups were compared to only the control group. Chi-square goodness-of-fit test was used to assess Mendelian inheritance patterns of gender and genotypes. Mouse bodyweight growth curves were analyzed by Mixed-effects analysis with Tukey’s multiple comparisons test. Kaplan-Meier mouse survival curves were analyzed by log-rank (Mantel-Cox) test followed by correction for multiple comparisons using the Holm-Šídák method. Categorical data was analyzed using two-sided Fisher’s exact test followed by correction for multiple comparisons using the Holm-Šídák method. Group analysis of data from multiple cell lines was performed using repeated measures two-way ANOVA followed by Dunnett’s multiple comparisons test, or by two-way Mixed-effects analysis. Statistical significance was defined as *p* < 0.05. Values are presented as mean ± SEM.

## Supporting information

Original Western blots Fig. 7D

Original Western blots Suppl. Fig. S7A

## Acknowledgements

We thank Bernhard Schmierer, Karolinska Genome Engineering (KGE) core facility, and Stephan Teglund, Karolinska Center for Transgene Technologies (KCTT), for expert help with generation of the *Trp53*^R210X^ mouse strain; Prof. Guillermina Lozano, MD Anderson Cancer Center, Houston, for kindly sharing the *Trp53*^R172H^ mouse strain used as a reference strain; Anna Malmerfelt at the Histology core facility, Dept. of Oncology-Pathology, Karolinska Institutet, for excellent help with preparation of sections and staining for histopathological analysis; and Alexis Gombert (DVM, IPSAV, ECVDI Diplomate), at the Diagnostic Imaging Clinic, University Animal Hospital, Swedish University of Agricultural Sciences, Uppsala, for expert help with interpreting and describing the MRI results.

## Author contribution

CS, VR, SÖ, AH and KGW designed the experiments. CS, VR, AH, SÖ and ASO performed the experiments. CS created the *Trp53*^R210X^ mouse strain and performed all mouse-related work. VR performed all histopathological assessment and analysis. All authors took part in analysis and interpretation of data. CS, VR, AH and KGW wrote the manuscript. KGW acquired funding for the project.

## Ethics

All animal studies were approved by the Stockholm Regional Animal Ethical Committee, Sweden (Dnr 14188-2019).

## Data availability

All original data are available upon request. This work does not include any datasets, such as DNA sequences or protein structure data, that have been uploaded in publicly available repositories.

## Additional information

Supplementary information is available at Cell Death & Disease’s website.

