## Supplementary material for "Mice carrying nonsense mutant p53 develop frequent multicentric or metastatic tumors": Original Western blots Suppl. Fig. S7A

### Original Western blots, Supplementary Figure S7A

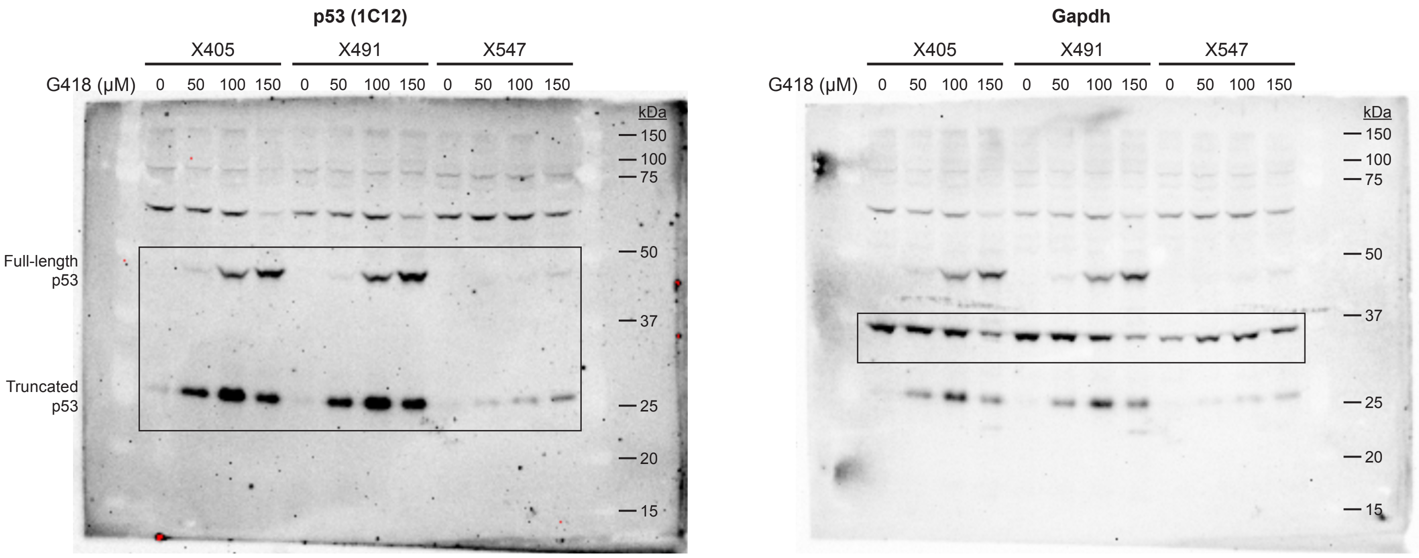


**Original Western blots, Supplementary Figure S7A.**
The Western blot membrane shown in Supplementary Fig. S7A was first blotted with the anti-p53 antibody 1C12 and an HRP-conjugated Rabbit-anti-Mouse IgG secondary antibody. 1C12 recognizes an N-terminal epitope and will therefore detect both full-length and C-terminally truncated p53. Following visualization of p53 and subsequent washing, the membrane was blotted with the HRP-conjugated G-9 anti-Gapdh antibody; Gapdh was used as a loading control. Squares indicate parts of the membrane shown in Supplementary Fig. S7A. Molecular weight markers are shown.
