## Supplementary Figure S1 for "Mice carrying nonsense mutant p53 develop frequent multicentric or metastatic tumors"

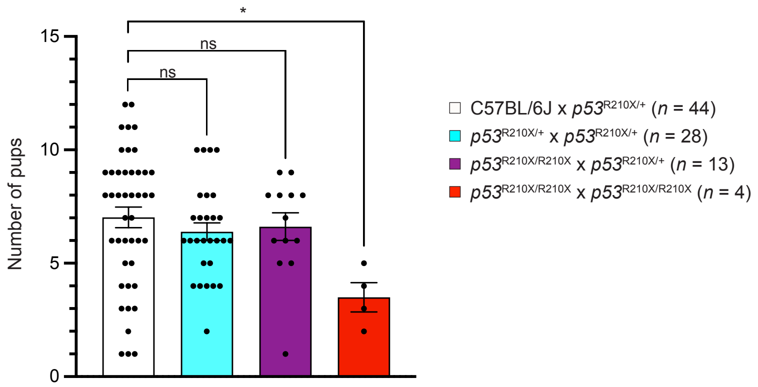


**Supplementary Figure S1. Intercrosses of *Trp53*$^{R210X/R210X}$ mice yield significantly fewer pups, related to Supplementary Table S1.**

Graph showing number of pups in each litter born from different intercrosses of *Trp53*$^{R210X}$ mice. Intercrosses involving *Trp53*$^{R210X/+}$ mice show an average litter size comparable to that of WT backcross breedings, whereas intercrosses of only *Trp53*$^{R210X/R210X}$ mice result in significantly smaller litters, *$p < 0.05$. Comparisons to backcross breeding with C57BL/6J WT mice were performed using one-way ANOVA followed by Dunnett's multiple comparison test. Bar heights indicate average number of pups/litter for the different crosses; number of litters for each intercross is indicated. SEM is shown.
