## Supplementary Figure S2A for "Mice carrying nonsense mutant p53 develop frequent multicentric or metastatic tumors"

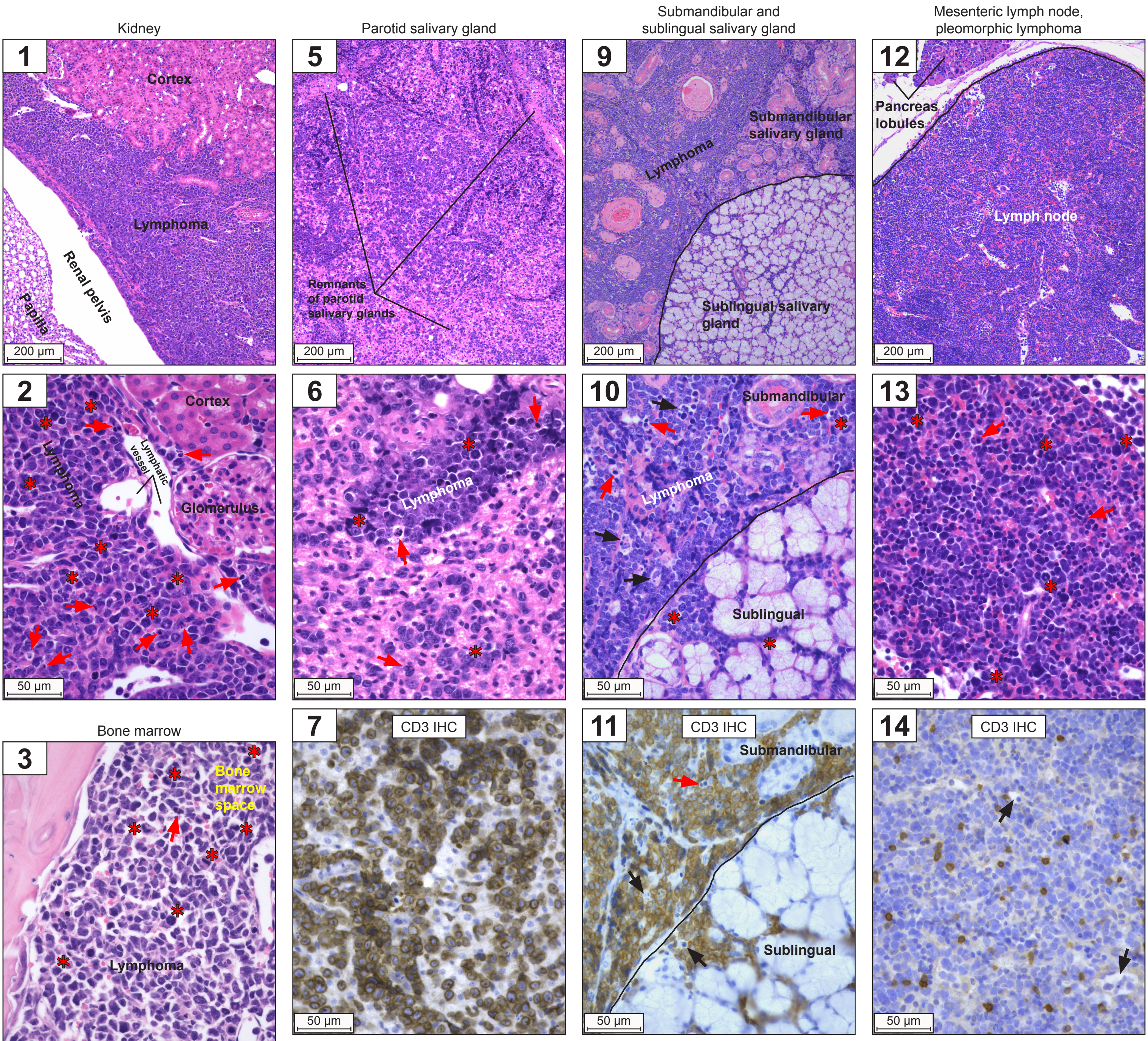


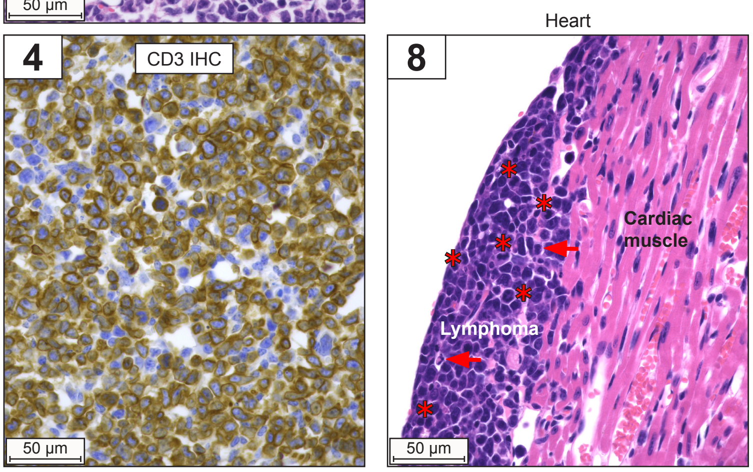


**Supplementary Figure S2A. Histomicrographs of T-cell lymphomas and pleomorphic lymphoma in kidney, bone marrow, heart, salivary glands and mesenteric lymph node from *Trp53*$^{R210X/R210X}$ mice, related to Figure 3.** Representative histomicrographs of lymphomas in different organs. Panel 1-11: T-cell lymphomas in kidney (1-2), bone marrow (3-4), parotid salivary gland (5-7), heart (8), submandibular and sublingual salivary gland (9-11). Panel 12-14: Pleomorphic lymphoma in mesenteric lymph node. Red asterisk: mitotic figure; red arrow: single-cell death; black arrow: tingible body machrophage. Black line in panels 9-12 delineates the expected outline of the organ indicated. Panel 1-3, 5-6, 8-10 and 12-13: hematoxylin-eosin staining. Immunohistochemical (IHC) staining in panel 4, 7, 11 and 14 shows CD3-positive cells with moderate to strong cytoplasmic and membrane staining. Note the ”starry sky” effect of the numerous tingible body macrophages indicated by black arrows in panel 14. Scale bars: 200µm and 50µm as indicated.
