## Supplementary Figure S2B for "Mice carrying nonsense mutant p53 develop frequent multicentric or metastatic tumors"

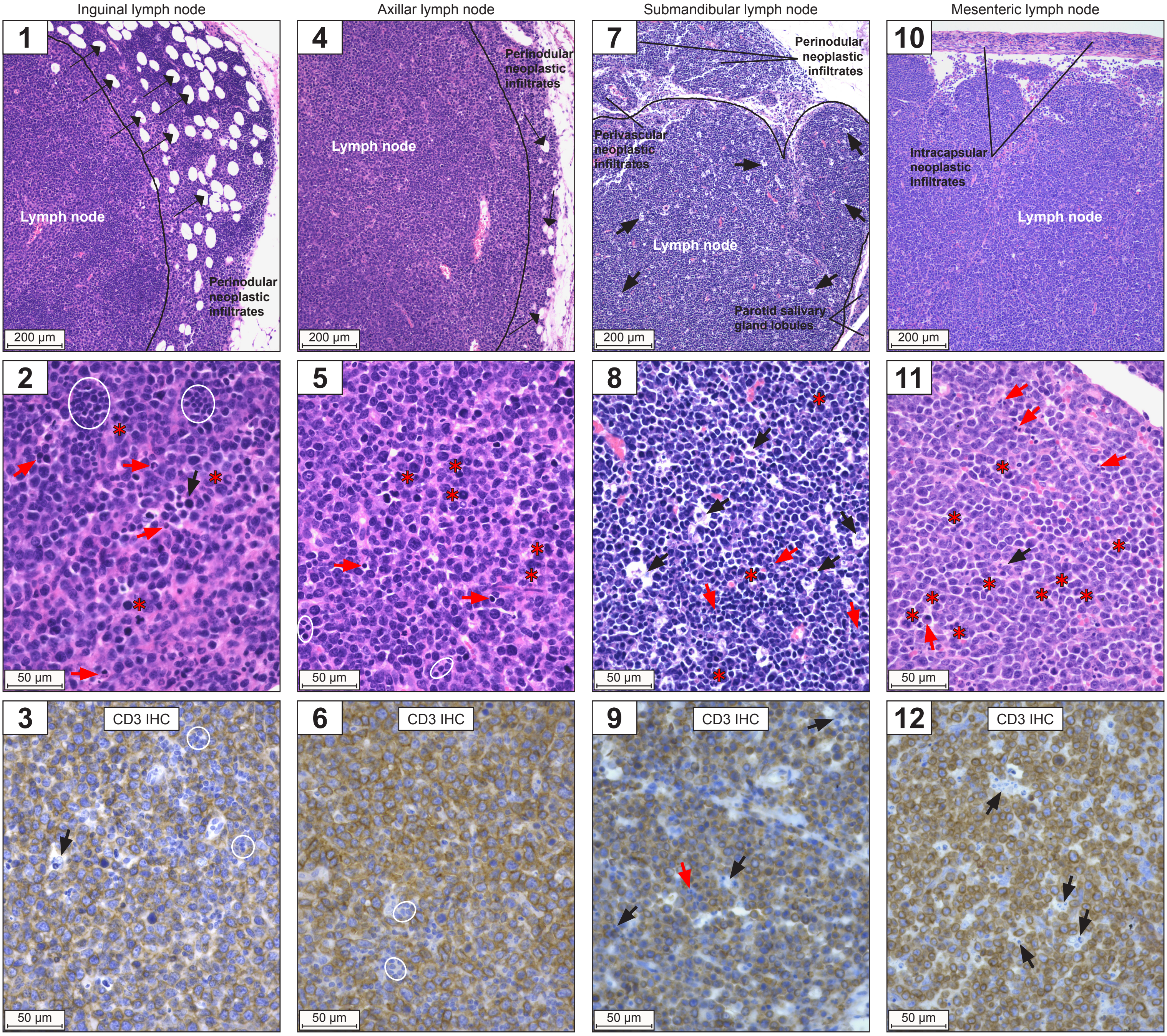


**Supplementary Figure S2B. Histomicrographs of T-cell lymphomas in inguinal, axillar, submandibular and mesenteric lymph nodes from *Trp53*$^{R210X/R210X}$ mice, related to Figure 3.**

Representative histomicrographs of T-cell lymphomas in inguinal (panels 1-3), axillar (4-6), submandibular (7-9) and mesenteric (10-12) lymph nodes from *Trp53*$^{R210X/R210X}$ mice. Red asterisk: mitotic figure; red arrow: single-cell death; black thick arrow: tingible body machrophage; black thin arrow: lipocytes in perinodal connective tissue. Black line in panels 1, 4 and 7 indicates approximate outline of lymph node. Upper and middle rows: hematoxylin-eosin staining. Aggregates of small, mature CD3 negative lymphocytes, i.e. remnants of the normal lymphocyte population in the lymph node, are circled. Note the ”starry sky” effect of the numerous tingible body macrophages indicated by black arrows in submandibular lymph node. Scale bars: 200µm (upper row) and 50µm (middle and bottom rows) as indicated.
