## Supplementary Figure S2C for "Mice carrying nonsense mutant p53 develop frequent multicentric or metastatic tumors"

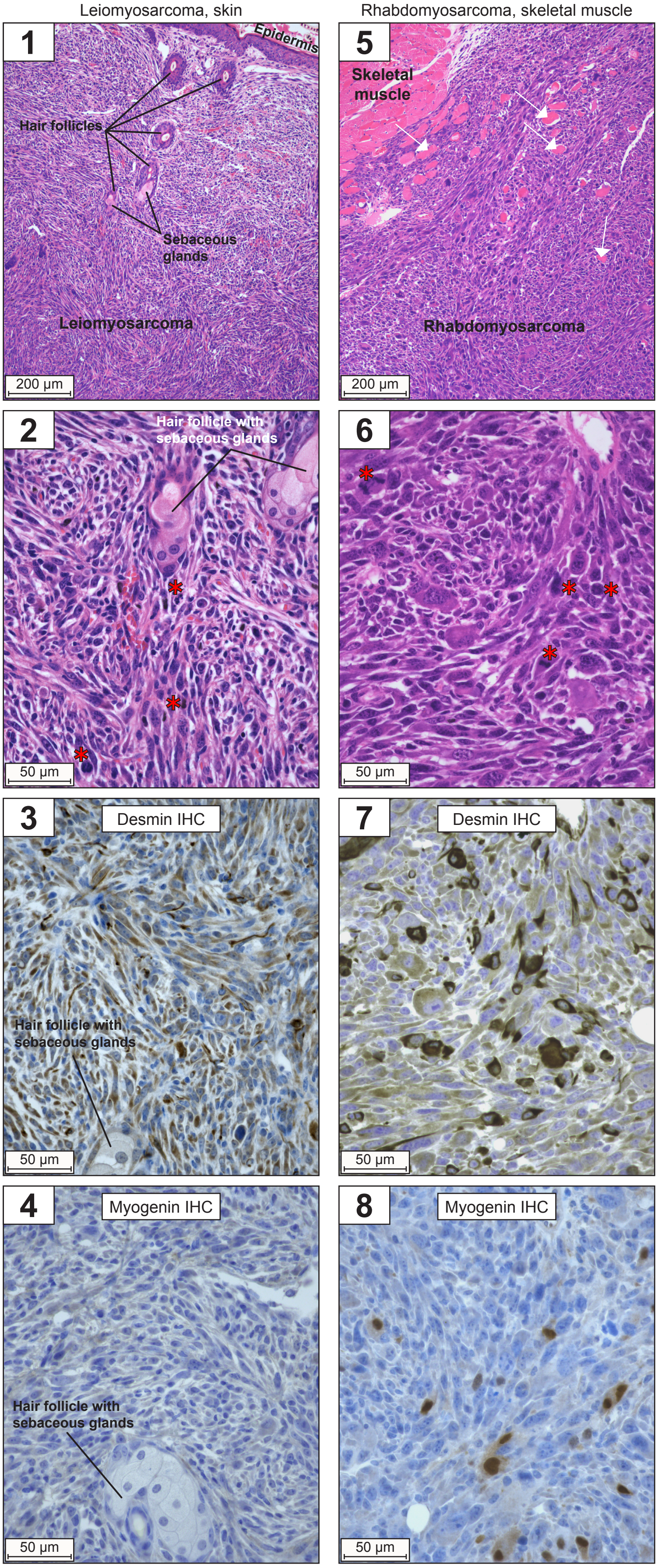


**Supplementary Figure S2C. Histomicrographs of leiomyosarcoma and rhabdomyosarcoma in skin and skeletal muscle, respectively, from *Trp53*$^{R210X/R210X}$ mice, related to Figure 3.**

Representative histomicrographs of leiomyosarcoma and rhabdomyosarcoma from *Trp53*$^{R210X/R210X}$ mice. Panel 1-2 and 5-6: hematoxylin-eosin staining. Red asterisk: mitotic figure; white thin arrow: entrapped well-differentiated myocoytes, found in the infiltrative borders of the tumor (panel 5). Panel 3 and 7: strong to moderate cytoplasmic desmin staining verifies myocyte origin. Panel 4: no staining for myogenin, consistent with leiomyocyte origin. Panel 8: dispersed neoplastic cells with strong nuclear myogenin staining verifies rhabdomyocyte origin. Scale bars: 200μm (upper row) and 50μm as indicated.
