## Supplementary Figure S3A for "Mice carrying nonsense mutant p53 develop frequent multicentric or metastatic tumors"

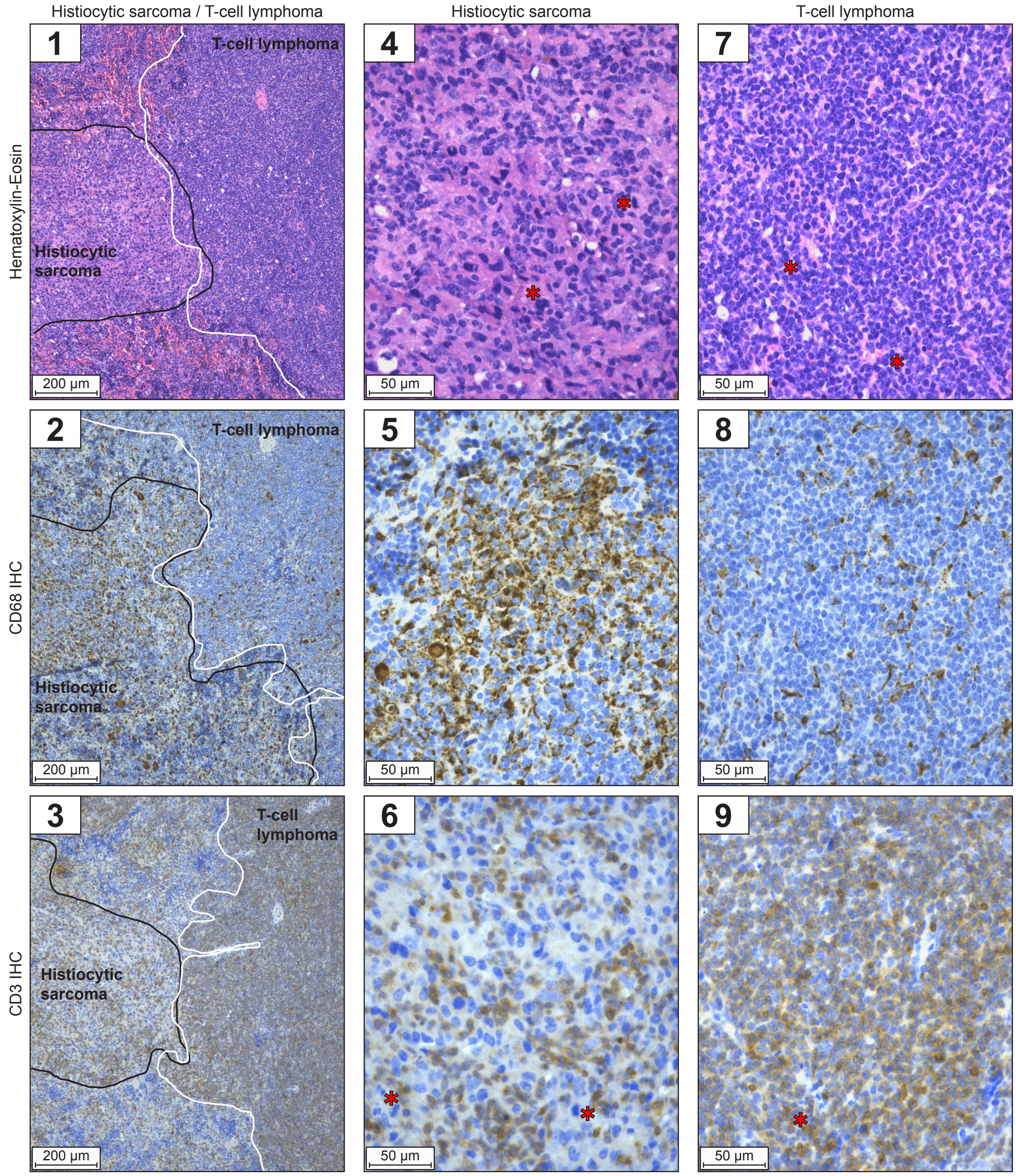


**Supplementary Figure S3A. Histomicrographs of histiocytic sarcoma and T-cell lymphoma in spleen from *Trp53*$^{R210X/+}$ mice, related to Figure 4.**

Panel 1-3: Spleen with histiocytic sarcoma (outlined by black line, left side of each panel) and T-cell lymphoma (outlined by white line, right side of each panel). Histiocytic sarcoma with numerous CD68-positive cells (moderate to strong cytoplasmic and membrane staining), i.e. histiocytes that form infiltrating sheets, with dispersed CD3-positive T-lymphocytes (strong cytoplasmic staining). T-cell lymphoma with numerous CD3-positive cells (moderate to strong cytoplasmic and membrane staining), i.e. T-lymphocytes that form expansive sheets, with few dispersed CD68-positive histiocytes (strong cytoplasmic staining). Scale bars: 200µm.

Panel 4-6: Histiocytic sarcoma with numerous neoplastic CD68-positive cells (moderate to strong cytoplasmic staining) with pleomorphic appearance and dispersed CD3-positive cells (moderate to strong cytoplasmic and membrane staining), i.e. T-lymphocytes.

Panel 7-9: T-cell lymphoma with numerous CD3-positive cells (moderate to strong cytoplasmic and membrane staining), i.e. T-lymphocytes, and few dispersed CD68-positive cells (strong cytoplasmic staining) with uniform appearance.

Note mitotic figures (red asterisks) in CD3-positive cells in the lymphoma and in CD3-negative cells in the histiocytic sarcoma. Scale bars: 50µm
