## Supplementary Figure S3B for "Mice carrying nonsense mutant p53 develop frequent multicentric or metastatic tumors"

**Histomicrographs of leiomyosarcoma in preputial gland and seminal vesicles, and rhabdomyosarcoma in cheek skin from *Trp53*$^{R210X/+}$ mice, related to Figure 4.**

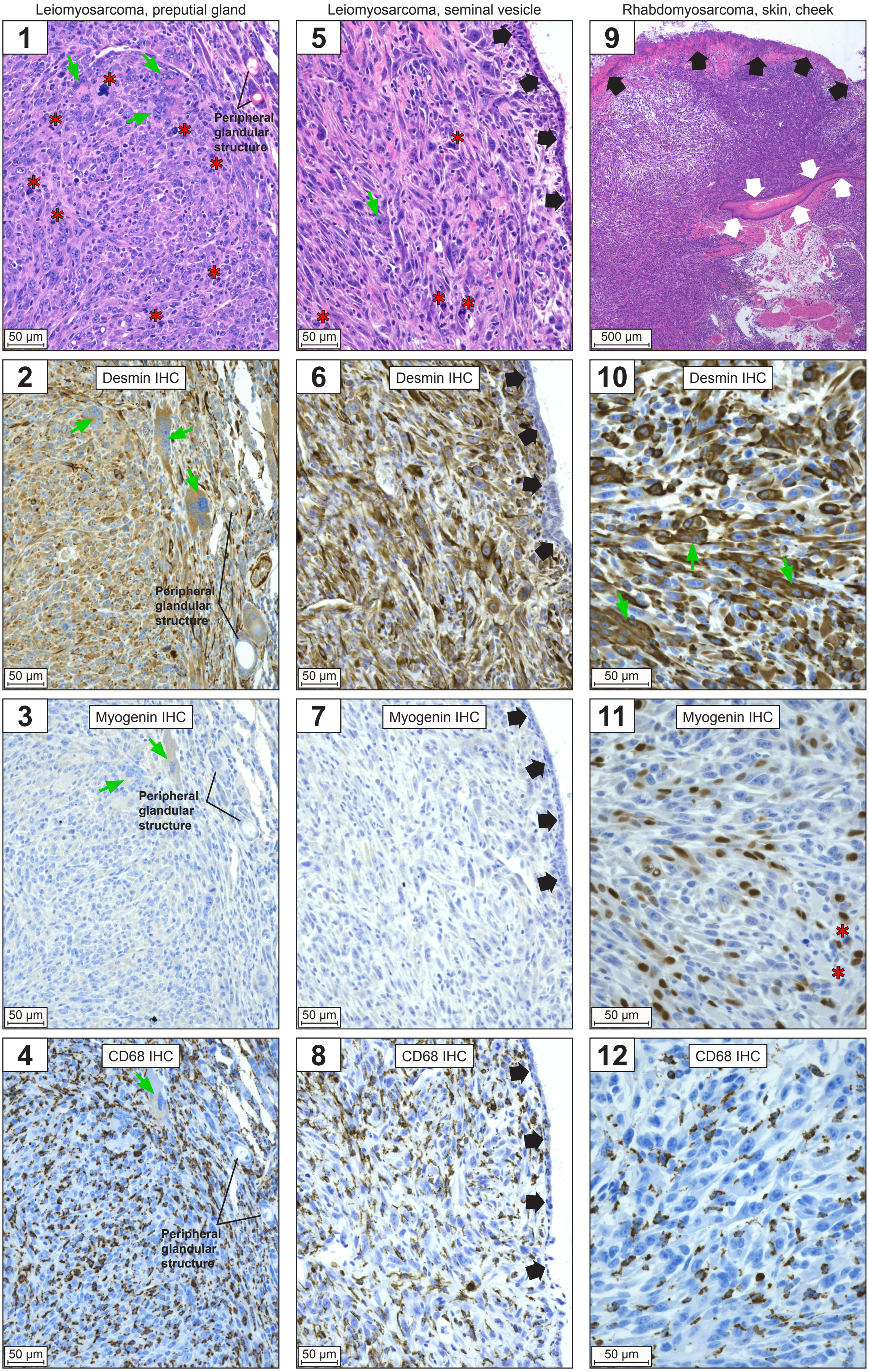


<u>Leiomyosarcoma.</u> Panel 1-4: Preputial gland, almost effaced by a dense proliferation of neoplastic mesenchymal cells. Only few peripheral glandular structures remain.
Panel 5-8: Seminal vesicle lamina propria is severely expanded and effaced by a dense proliferation of neoplastic mesenchymal cells. The epithelial lining is mostly intact (short broad black arrows).
The neoplastic cells in panels 1-8 are spindle-shaped to polygonal and form broad interlacing bundles in a sparse stroma. The cells are highly pleomorphic, with giant multinucleated cells (green arrows) and mitotic figures (red asterisks). Panel 1 and 5: hematoxylin-eosin staining. IHC confirms leiomyocyte origin: all neoplastic cells have moderate to strong cytoplasmic desmin staining, negative nuclear myogenin staining and negative CD68 staining. CD68-positive cells are dispersed, uniform histiocytes infiltrating the tumor. Scale bars: 50µm.

<u>Rhabdomyosarcoma (skin, cheek).</u> Panel 9-12: The epidermis is ulcerated (broad black arrows) with an intact epidermal fold in part of the section (broad white arrows). The dermis and subcutis is severely expanded and effaced by a dense proliferation of neoplastic mesenchymal cells. The cells are highly pleomorphic, with elongated multinucleated cells with the nuclei lining up, i.e. ”strap-cells” (green arrows), and mitotic figures (red asterisks). Panel 9: hematoxylin-eosin staining. IHC confirms rhabdomyocyte origin: all neoplastic cells have moderate to strong cytoplasmic desmin staining, some also have strong nuclear myogenin staining, and all have negative CD68 staining. CD68-positive cells are dispersed, uniform histiocytes infiltrating the tumor. Scale bars: 500µm (panel 9) and 50µm (panel 10-12) as indicated.
