## Supplementary Figure S3C for "Mice carrying nonsense mutant p53 develop frequent multicentric or metastatic tumors"

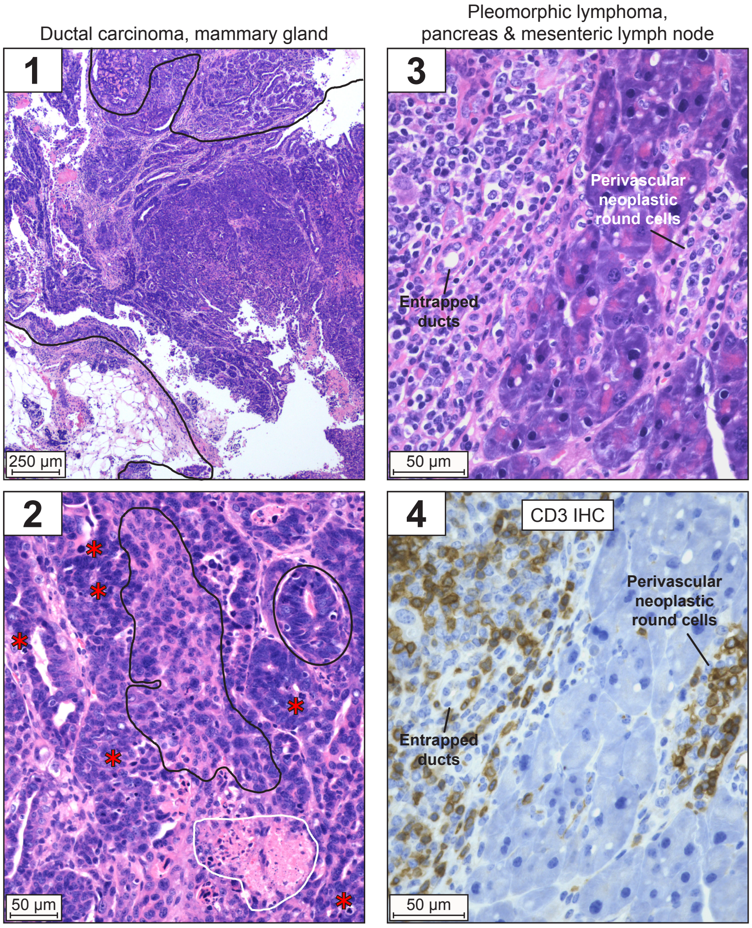


**Supplementary Figure S3C. Histomicrographs of ductal carcinoma and pleomorphic lymphoma in mammary gland and pancreas, respectively, from *Trp53*$^{R210X/+}$ mice, related to Figure 4.**

Ductal carcinoma in mammary gland. Panel 1 shows a malignant epithelial tumor (between black lines) with normal mammary gland tissue in the lower part of the image, and with hyperplastic mammary gland tissue in the upper part. Panel 2 shows dense proliferations of neoplastic epithelial cells that form ductular (circled) and solid proliferations (black solid outline) in a sparse stroma with multifocal necroses (white solid outline). Red asterisks: mitotic figures.

Pleomorphic lymphoma in pancreas and mesenteric lymph node. Panel 3 shows that the pancreas is multifocally effaced by neoplastic round cells (left side of the image) which are also found perivascularly in the more intact parts of the pancreas. In the effaced area, remaining entrapped ducts can be seen. Panel 4 shows that about half, or less than half, of the neoplastic round cells are CD3-positive.

Panel 1-3: hematoxylin-eosin staining. Scale bars: 250µm and 50µm as indicated.
