## Supplementary Figure S4A for "Mice carrying nonsense mutant p53 develop frequent multicentric or metastatic tumors"

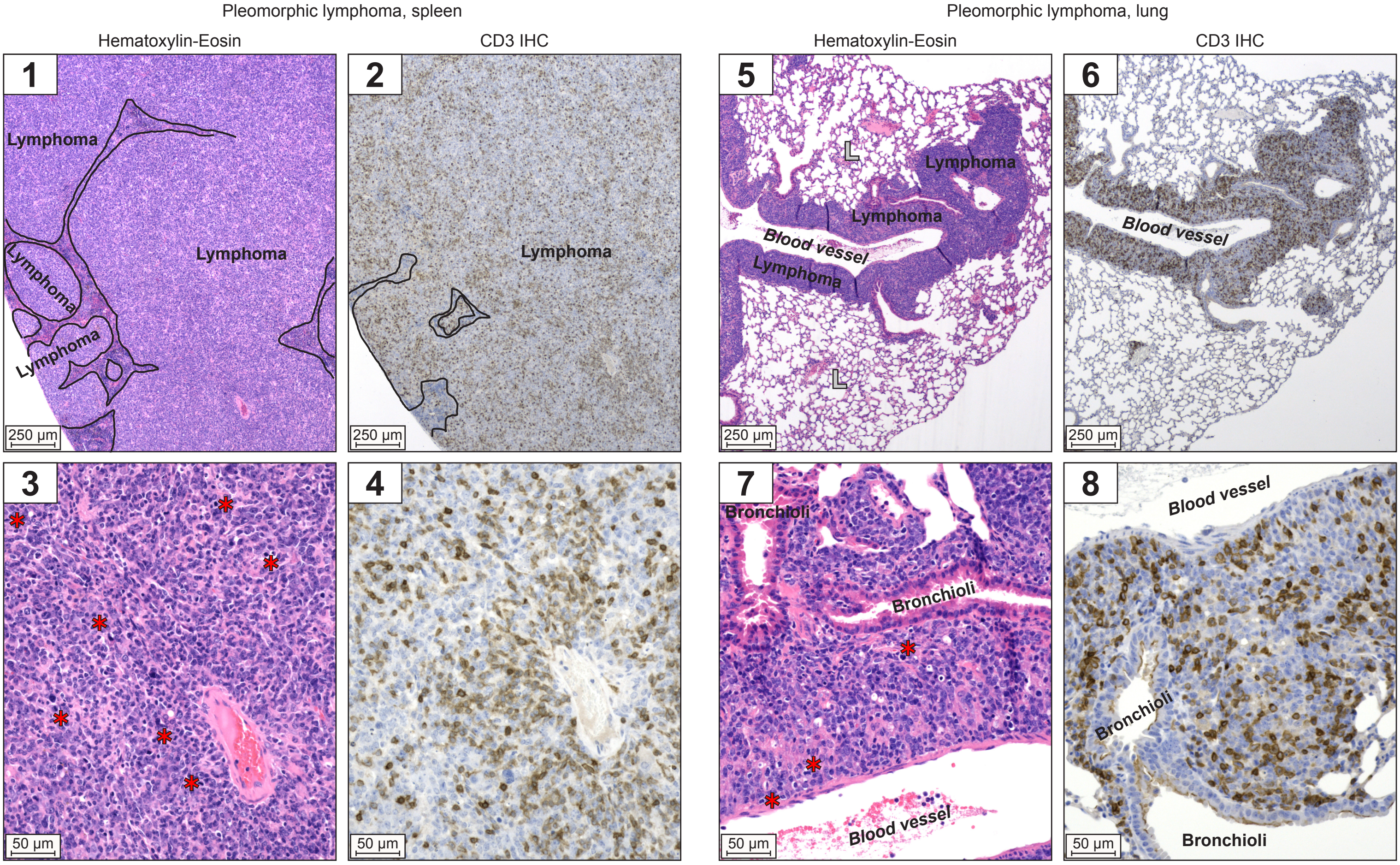


**Supplementary Figure S4A. Histomicrographs of pleomorphic lymphoma in spleen and lung from WT mice, related to Figure 5A.**

Pleomorphic lymphoma in spleen. Panels 1-4 show dense multinodular to coalescing infiltrates (outlined in black) of neoplastic lymphocytes, a subpopulation of which are strongly CD3-positive, that efface almost all of the normal spleen architecture, and diffusely infiltrate the remaining parts. Red asterisks: mitotic figures.

Pleomorphic lymphoma in lung. Panels 5-8 show dense perivascular infiltrates of neoplastic lymphocytes, a subpopulation of which are strongly CD3-positive. Red asterisks: mitotic figures; L: normal lung parenchyma.

Scale bars: 250µm and 50µm as indicated.
