## Supplementary Figure S4B for "Mice carrying nonsense mutant p53 develop frequent multicentric or metastatic tumors"

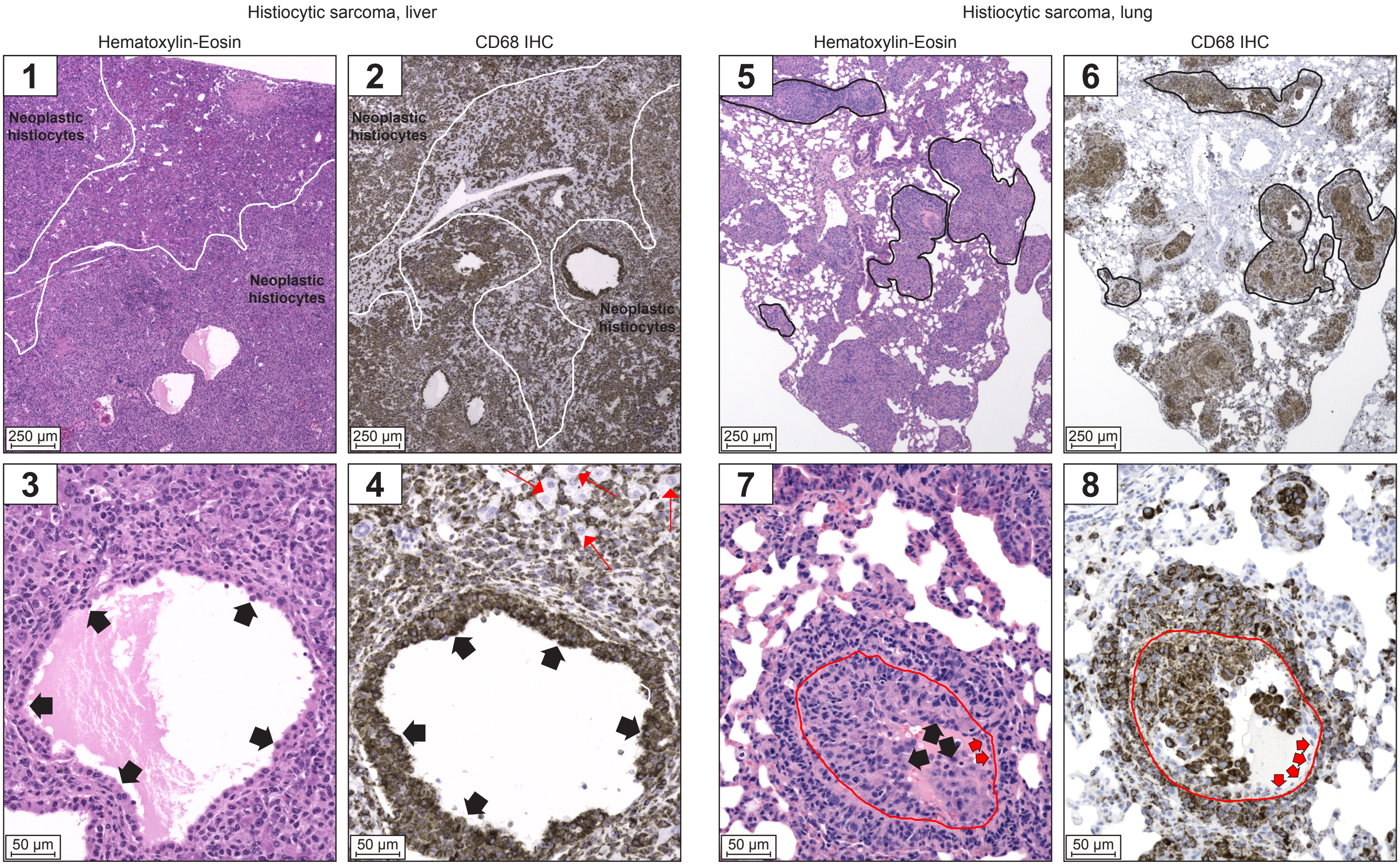


**Supplementary Figure S4B. Histomicrographs of histiocytic sarcoma in liver and lung from WT mice, related to Figure 5A.**

Histiocytic sarcoma in liver. Panels 1-2 show dense infiltrates of neoplastic histiocytes (outlined by white solid line) efface large parts of the liver, and diffusely infiltrate the remaining parts. Panels 3-4 show a very dilated blood vessel surrounded by dense mats of neoplastic histiocytes. Neoplastic histiocytes are piling up along the endothelium (short broad black arrows) and efface the vessel wall. There are few remaining viable hepatocytes (red thin arrows) among the pleomorphic strongly CD68-positive neoplastic cells.

Histiocytic sarcoma in lung. Panels 5-6 show multinodular to coalescing, vessel centered, dense infiltrates of pleomorphic, strongly CD68-positive neoplastic histiocytes (examples outlined in black solid line) efface large parts of the lung. Panels 7-8 show a very dilated blood vessel surrounded by dense mats of neoplastic histiocytes. Neoplastic histiocytes are piling up along the endothelium (short broad black arrows) and efface the vessel wall. Only small rests of normal, flat endothelium (short broad red arrows) remain. Red line demarcates the approximate location of the endothelial basal lamina.

Scale bars: 250µm and 50µm as indicated.
