## Supplementary Figure S4C for "Mice carrying nonsense mutant p53 develop frequent multicentric or metastatic tumors"

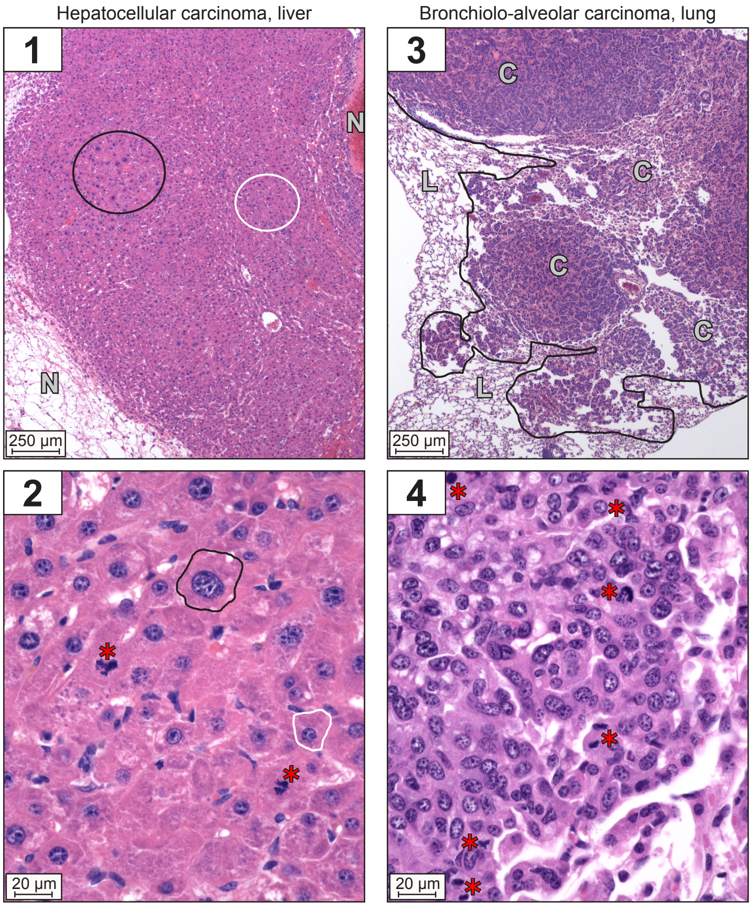


**Supplementary Figure S4C. Histomicrographs of hepatocellular and bronchiolo-alveolar carcinoma in liver and lung, respectively, from WT mice, related to Figure 5A.**

Hepatocellular carcinoma in liver. Panel 1 shows disorganized proliferation of neoplastic hepatocytes in sparse stroma with multifocal necroses (indicated by N). Panel 2 shows the neoplastic hepatocytes are highly pleomorphic with very high anisokaryosis (variation in nuclear size) and anisocytosis (variation in cell size), and a high mitotic rate with numerous bizarre mitotic figures (red asterisks). The neoplastic hepatocytes vary from very enlarged with a very large nucleus (black solid line) to more similar in size to normal hepatocytes (white solid line).

Bronchiolo-alveolar carcinoma in lung. Panel 3 shows poorly circumscribed, multinodular proliferation of neoplastic epithelial cells forming papillary, tubular and solid areas in sparse stroma (outlined in black; C: carcinoma; L: lung). Panel 4 shows the neoplastic epithelial cells are pleomorphic with numerous mitotic figures (red asterisks).

All histomicrographs are stained with hematoxylin-eosin. Scale bars: 250µm and 20µm as indicated.
