## Supplementary Figure S5 for "Mice carrying nonsense mutant p53 develop frequent multicentric or metastatic tumors"

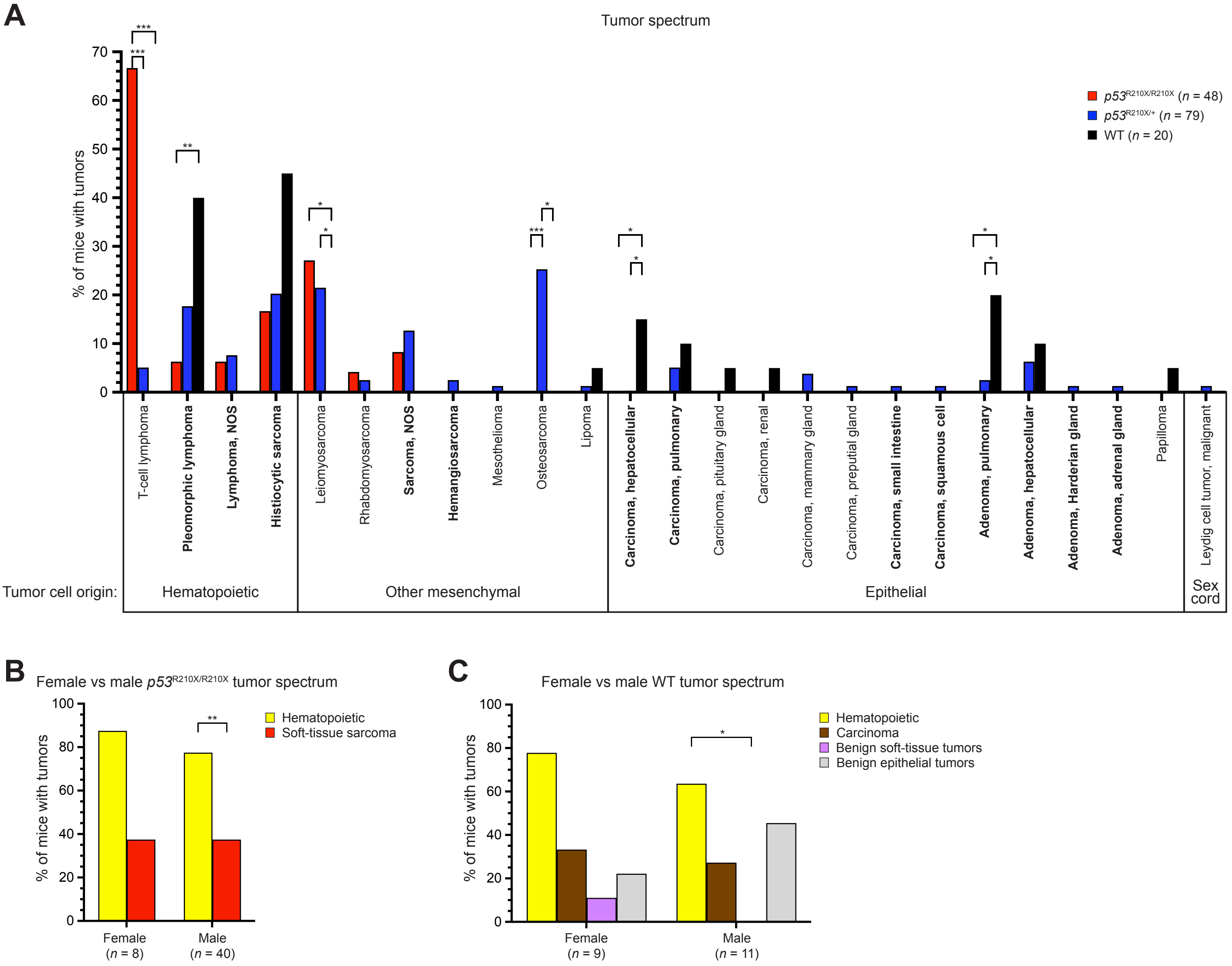


**Supplementary Figure S5. Full tumor spectrum for *Trp53*$^{\mathbf{R210X/R210X}}$, *Trp53*$^{\mathbf{R210X/+}}$ and WT mice, and female vs male tumor type comparisons for *Trp53*$^{\mathbf{R210X/R210X}}$ and WT mice, related to Figure 5A and D.**

**(A)** All identified tumors in the three genotypes, showing the large variation in tumor types found in *Trp53*$^{R210X/+}$ mice, and illustrating the overlap of tumor types with WT mice. Bold tumor type: found in C57BL/6J WT mice as described in Elies *et. al.*, 2024 (36). Comparison of tumor spectra in female and male **(B)** *Trp53*$^{R210X/R210X}$ and **(C)** WT mice. There were no differences between sexes in either genotype. Male *Trp53*$^{R210X/R210X}$ mice had a higher incidence of hematopoietic tumors than soft-tissue sarcomas, and male WT mice had a higher incidence of hematopoietic tumors than benign soft-tissue tumors. All statistical analysis was performed by two-sided Fisher's exact test. In **A**, comparisons were made between genotypes within each tumor type. In **B** and **C**, comparisons were made between female and male outcome for each tumor type (i.e. female hematopoietic tumors vs male hematopoietic tumors, etc.), and between all tumor types within one sex (i.e. all tumor types found in females). Corrections for multiple comparisons were performed using the Holm-Šídák method. Adjusted P-values shown: *$p < 0.05$, **$p < 0.01$, ***$p < 0.001$.
