## Supplementary Figure S6 for "Mice carrying nonsense mutant p53 develop frequent multicentric or metastatic tumors"

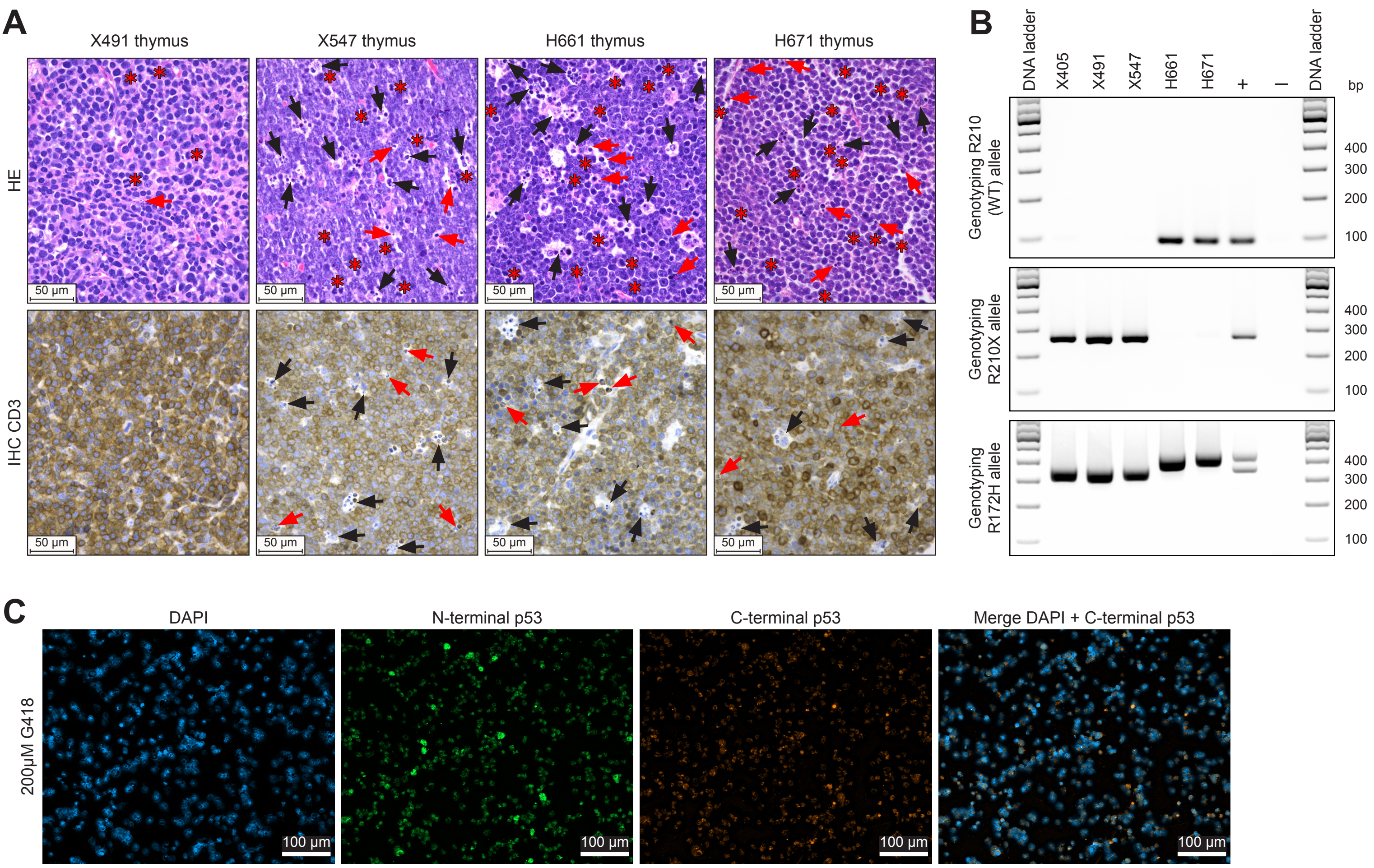


**Supplementary Figure S6. Verification of mouse T-lymphoma cell lines and cell death induced by aminoglycoside G418, related to Figure 7.**

**(A)** Representative histomicrographs of the thymus lymphomas from mice X491, X547, H661 and H671 stained with hematoxylin-eosin (HE, upper) and CD3 immunohistochemistry (IHC, lower). Mats of CD3-positive, i.e. T-cell origin, pleomorphic neoplastic lymphocytes with numerous mitoses (red asterisks), tingible body macrophages (black arrows), and cell fragments consistent with single-cell death (red arows) efface the normal thymic architecture in all thymi. **(B)** Verification of *Trp53*$^{R210X/R210X}$ (X405, X491 and X547) and *Trp53*$^{R172H/R172H}$ (H661 and H671) T-lymphoma cell line genotypes using the same PCR genotyping strategies as for mouse genotyping of the respective strain. All cell lines were genotyped using both protocols to ensure no cross-contamination has occurred. R210 (WT) allele is identified by a 95 bp PCR fragment (top panel) and R210X mutant allele by a 252 bp fragment (middle panel). Genotyping using the *Trp53*$^{R172H/R172H}$ strategy identifies WT allele by a 342 bp PCR fragment and R172H mutant allele by a 410 bp fragment (bottom panel) according to Lang *et. al.*, 2004 (9). All cell lines show clean and expected genotypes (X405, X491 and X547 = *Trp53*$^{R210X/R210X}$; and H661 and H671 = *Trp53*$^{R172H/R172H}$). **(C)** Representative immunofluorescence staining of X405 *Trp53*$^{R210X/R210X}$ T-lymphoma cells after 72h treatment with 200µM aminoglycoside G418. p53 was detected using N1 (N-terminal epitope) and 280aa C-term (C-terminal epitope) antibodies. Rightmost panel show merged DAPI and 280aa C-term p53 staining. This concentration of G418 caused substantial cell death, consistent with the Annexin V data shown in Fig. 7G. Scale bars: 50µm **(B)** and 100µm **(C)**.
