## Supplementary Figure S7 for "Mice carrying nonsense mutant p53 develop frequent multicentric or metastatic tumors"

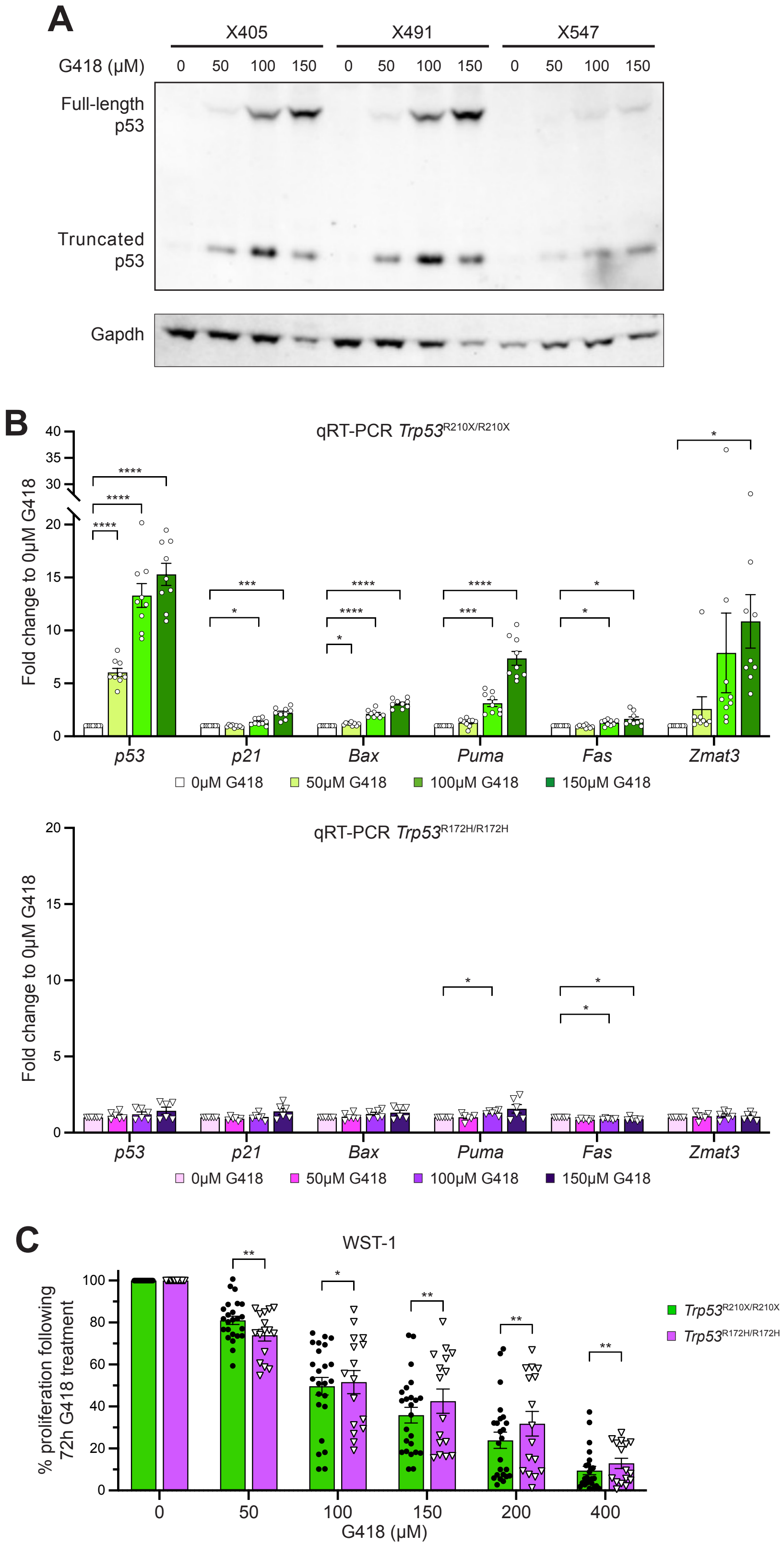


**Supplementary Figure S7. Induction of full-length p53, upregulation of p53 target genes, and inhibition of cell proliferation in *Trp53*$^{R210X/R210X}$ T-lymphoma cells following G418 treatment, related to Figure 7.**

**(A)** Western blot analysis showing dose-dependent induction of full-length p53 in three T-lymphoma cell lines following 72h treatment with G418 at indicated concentrations. p53 was visualized using the anti-p53 antibody 1C12 that recognizes an N-terminal epitope and therefore detects both full-length and C-terminally truncated p53. Gapdh was used as a loading control. **(B)** qRT-PCR results from Figure 7E showing all individual values from each cell line from 3 independent experiments per cell line. Gene expression values are normalized to *Gapdh* expression and compared to untreated (0µM G418) negative control for each gene. Upper panel: *Trp53*$^{R210X/R210X}$ T-lymphoma lines ($n$ = 3); lower panel: *Trp53*$^{R172H/R172H}$ T-lymphoma lines ($n$ = 2). **(C)** WST-1 assay results from Figure 7F showing all individual values from each cell line from 8 independent experiments per cell line. In **B** and **C**, statistical analysis was performed by repeated measures two-way ANOVA followed by Dunnett's multiple comparisons test **(B)**, and by two-way Mixed-effects analysis **(C)**, respectively. Mean ± SEM are indicated. Adjusted P-values: *$p$ < 0.05, **$p$ < 0.01, ***$p$ < 0.001, ****$p$ < 0.0001.
