## Supplementary Figure S8 for "Mice carrying nonsense mutant p53 develop frequent multicentric or metastatic tumors"

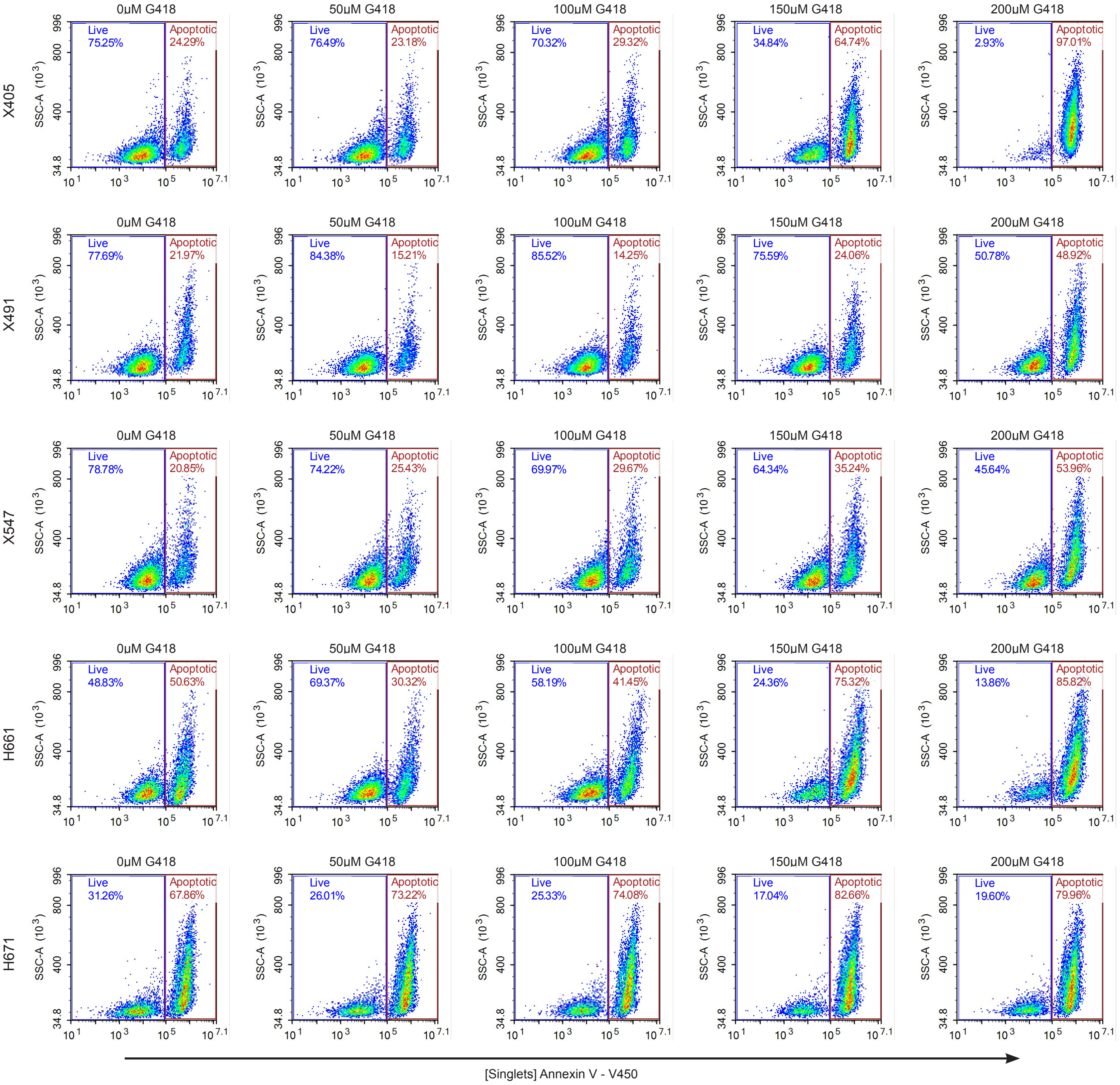


**Supplementary Figure S8. Annexin V flow cytometry of *Trp53*$^{R210X/R210X}$ and *Trp53*$^{R172H/R172H}$ T-lymphoma cell lines following G418 treatment, related to Figure 7G.**

Represenative plots from Annexin V flow cytometry of *Trp53*$^{R210X/R210X}$ (X405, X491 and X547) and *Trp53*$^{R172H/R172H}$ (H661 and H671) mouse T-lymphoma cell lines follwing treatment with G418 for 72h at indicated concentrations; 3 independent experiments per cell line were performed.
