## Supplementary Table S1 for "Mice carrying nonsense mutant p53 develop frequent multicentric or metastatic tumors"

**Supplementary Table S1.** Average litter sizes at 2 weeks of age for different combinations of *Trp53* $^{R210X}$ crosses. Comparisons are made to backcross breeding with C57BL/6J WT mice (top row) using one-way ANOVA followed by Dunnett's multiple comparison test. Number of pups in each litter is plotted in **Supplementary Figure S1.** ns = not significant, *$p < 0.05$.

| **Type of cross** | **No. of litters** | **No. of pups** | **Average litter size** | ***p*-value** |
|---|---|---|---|---|
| **C57BL/6J x *Trp53*** $^{R210X/+}$ | 44 | 309 | 7.02 | |
| ***Trp53*** $^{R210X/+}$ **x *Trp53*** $^{R210X/+}$ | 28 | 179 | 6.39 | 0.6643 (ns) |
| ***Trp53*** $^{R210X/R210X}$ **x *Trp53*** $^{R210X/+}$ | 13 | 86 | 6.62 | 0.9400 (ns) |
| ***Trp53*** $^{R210X/R210X}$ **x *Trp53*** $^{R210X/R210X}$ | 4 | 14 | 3.50 | 0.0304 (*) |
