## Supplementary Table S2 for "Mice carrying nonsense mutant p53 develop frequent multicentric or metastatic tumors"

**Supplementary Table S2.** Gender and genotype distribution for progeny of different combinations of *Trp53* $^{R210X}$ crosses. Intercrosses between *Trp53* $^{R210X/+}$ and/or *Trp53* $^{R210X/R210X}$ resulted in fewer than expected female *Trp53* $^{R210X/R210X}$ offspring, indicated with bold, red text. *$p$ < 0.05, **$p$ < 0.01, ***$p$ < 0.001, ****$p$ < 0.0001 (Chi-square goodness-of-fit test).

| **Backcross** | | **C57BL/6J x *Trp53* $^{R210X/+}$** | | | | |
|---|---|---|---|---|---|---|
| **Progeny Genotype (*n* = 309)** | **Males** | | **Females** | | ***p*-value** | |
| | **Observed no.** | **Expected no.** | **Observed no.** | **Expected no.** | **Both genders** | **Females only** |
| **WT (N = 169)** | 101 (32.69%) | 77.25 (25%) | 68 (22.01%) | 77.25 (25%) | 0.0153 (*) | - |
| ***Trp53* $^{R210X/+}$ (*n* = 140)** | 75 (24.27%) | 77.25 (25%) | 65 (21.04%) | 77.25 (25%) | | |
| **Intercross** | | ***Trp53* $^{R210X/+}$ x *Trp53* $^{R210X/+}$** | | | | |
| **Progeny Genotype (*n* = 179)** | **Males** | | **Females** | | ***p*-value** | |
| | **Observed no.** | **Expected no.** | **Observed no.** | **Expected no.** | **Both genders** | **Females only** |
| **WT (*n* = 48)** | 23 (12.85%) | 22.38 (12.5%) | 25 (13.97%) | 22.38 (12.5%) | | |
| ***Trp53* $^{R210X/+}$ (*n* = 102)** | 48 (26.82%) | 44.75 (25%) | 54 (30.17%) | 44.75 (25%) | 0.0031 (**) | 0.0001 (***) |
| ***Trp53* $^{R210X/R210X}$ (*n* = 29)** | 25 (13.97%) | 22.38 (12.5%) | **4 (2.24%)** | 22.38 (12.5%) | | |
| **Intercross** | | ***Trp53* $^{R210X/R210X}$ x *Trp53* $^{R210X/+}$** | | | | |
| **Progeny Genotype (*n* = 86)** | **Males** | | **Females** | | ***p*-value** | |
| | **Observed no.** | **Expected no.** | **Observed no.** | **Expected no.** | **Both genders** | **Females only** |
| ***Trp53* $^{R210X/+}$ (*n* = 55)** | 24 (27.91%) | 21.5 (25%) | 31 (36.05%) | 21.5 (25%) | 0.0045 (**) | 0.0003 (***) |
| ***Trp53* $^{R210X/R210X}$ (*n* = 31)** | 23 (26.74%) | 21.5 (25%) | **8 (9.30%)** | 21.5 (25%) | | |
| **Intercross** | | ***Trp53* $^{R210X/R210X}$ x *Trp53* $^{R210X/R210X}$** | | | | |
| **Progeny Genotype (*n* = 14)** | **Males** | | **Females** | | ***p*-value** | |
| | **Observed no.** | **Expected no.** | **Observed no.** | **Expected no.** | **Both genders** | **Females only** |
| ***Trp53* $^{R210X/R210X}$** | 12 (85.71%) | 7 (50%) | **2 (14.29%)** | 7 (50%) | 0.0074 (**) | - |
| | | **Summary of *Trp53* $^{R210X/R210X}$ pups from all different types of intercrosses** | | | | |
| **Progeny Genotype (*n* = 74)** | **Males** | | **Females** | | ***p*-value** | |
| | **Observed no.** | **Expected no.** | **Observed no.** | **Expected no.** | **Both genders** | **Females only** |
| ***Trp53* $^{R210X/R210X}$** | 60 (81.08%) | 37 (50%) | **14 (18.92%)** | 37 (50%) | <0.0001 (****) | - |
