## Supplementary Table S3 for "Mice carrying nonsense mutant p53 develop frequent multicentric or metastatic tumors"

**Supplementary Table S3.** Histopathology results, expressed as % of total number of mice with tumors unless otherwise indicated. Data was analyzed using a two-sided Fisher's exact test followed by correction for multiple analyses using the Holm-Šídák method. There were several differences between genotypes, and a difference between sex within genotype was only seen in *Trp53*$^{R210X/+}$ mice, indicated in bold, red text. ns = not significant, *$p$ < 0.05, **$p$ < 0.01, ***$p$ < 0.001, ****$p$ < 0.0001.

| | *Trp53*$^{R210X/R210X}$ (HO) | | | *Trp53*$^{R210X/+}$ (HZ) | | | WT | | | Genotype differences (adjusted $p$-values) | | |
|---|---|---|---|---|---|---|---|---|---|---|---|---|
| | All ($n$ = 49) | Females ($n$ = 8) | Males ($n$ = 41) | All ($n$ = 79) | Females ($n$ = 37) | Males ($n$ = 42) | All ($n$ = 24) | Females ($n$ = 12) | Males ($n$ = 12) | HO vs HZ | HO vs WT | HZ vs WT |
| **Mice with tumors, total** (*no. of mice*) | 98.0 (48) | 100 (8) | 97.6 (40) | 100 (79) | 100 (37) | 100 (42) | 83.3 (20) | 75.0 (9) | 91.7 (11) | 0.3828 (ns) | 0.0736 (ns) | 0.0072 (**) |
| **Multicentric or metastatic tumors,** % mice with tumors *(no. of mice)* | 60.4 (29) | 62.5 (5) | 60.0 (24) | 36.7 (29) | 43.2 (16) | 31.0 (13) | 50.0 (10) | 55.6 (5) | 45.5 (5) | 0.0318 (*) | 0.5912 (ns) | 0.5273 (ns) |
| **Hematopoietic tumors (HT), total** (*no. of mice*) | 79.2 (38) | 87.5 (7) | 77.5 (31) | 40.5 (32) | 37.8 (14) | 42.8 (18) | 70.0 (14) | 77.8 (7) | 63.6 (7) | 0.0003 (***) | 0.5319 (ns) | 0.0474 (*) |
| **HT multicentric,** % of mice with HT (*no. of mice*) | 76.3 (29) | 71.4 (5) | 77.4 (24) | 68.7 (22) | 78.5 (11) | 61.1 (11) | 71.4 (10) | 71.4 (5) | 71.4 (5) | 0.9320 (ns) | 0.9320 (ns) | 0.9999 (ns) |
| **T-cell lymphoma (TCL),** % of mice with HT (*no. of mice*) | 84.2 (32) | 85.7 (6) | 83.9 (26) | 12.5 (4) | 7.1 (1) | 16.7 (3) | 0 | 0 | 0 | 0.0003 (***) | 0.0003 (***) | 0.1507 (ns) |
| **TCL multicentric**, % of mice with TCL (*no. of mice*) | 78.1 (25) | 83.3 (5) | 76.9 (20) | 100 (4) | 100 (1) | 100 (3) | 0 | 0 | 0 | 0.5658 (ns) | - | - |
| **Soft-tissue sarcoma (STS), total** (*no. of mice*) | 37.5 (18) | 37.5 (3) | 37.5 (15) | 40.5 (32) | **27.0 (10)**[a] | **50.0 (21)**[a] | 0 | 0 | 0 | 0.8517 (ns) | 0.0016 (**) | 0.0009 (***) |
| **STS metastasis**, % of mice with STS (*no. of mice*) | 5.6 (1) | 0 | 6.7 (1) | 9.4 (3) | 10 (1) | 9.5 (2) | 0 | 0 | 0 | 0.9999 (ns) | - | - |
| **Osteosarcoma (OS), total** (*no. of mice*) | 0 | 0 | 0 | 25.3 (20) | **51.4 (19)**[b] | **2.4 (1)**[b] | 0 | 0 | 0 | 0.0003 (***) | 0.9999 | 0.0207 (*) |
| **OS metastases**, % of mice with OS (*no. of mice*) | 0 | 0 | 0 | 15.0 (3) | 15.8 (3) | 0 | 0 | 0 | 0 | - | - | - |
| **Carcinoma (C), total** (*no. of mice*) | 0 | 0 | 0 | 12.7 (10) | 13.5 (5) | 11.9 (5) | 30 (6) | 33.3 (3) | 27.3 (3) | 0.0260 (*) | 0.0012 (**) | 0.0860 (ns) |
| **C metastases**, % of mice with C (*no. of mice*) | 0 | 0 | 0 | 10.0 (1) | 20.0 (1) | 0 | 0 | 0 | 0 | - | - | 0.9999 (ns) |
| **Sex cord tumor, total** (*no. of mice*) | 0 | 0 | 0 | 1.3 (1) | 0 | 2.4 (1) | 0 | 0 | 0 | 1.0000 (ns) | 1.0000 (ns) | 1.0000 (ns) |
| **Benign tumors, total** (*no. of mice*) | 0 | 0 | 0 | 12.7 (10) | 8.1 (3) | 16.7 (7) | 40.0 (8) | 33.3 (3) | 45.5 (5) | 0.0175 (*) | 0.0003 (***) | 0.0175 (*) |

[a] $p$ = 0.0416 (*)

[b] $p$ < 0.0001 (****)
