## Supplementary Table S4A for "Mice carrying nonsense mutant p53 develop frequent multicentric or metastatic tumors"

**Supplementary Table S4A.** Results of immunohistochemistry (IHC) analysis of a selection of neoplasias in *Trp53*$^{R210X/R210X}$ (HO) mice. CD3 was used to identify cells of T lymphocyte lineage; vimentin to identify mesenchymal cells and verify sarcoma; CD68 to identify cells of histiocytic lineage; desmin to identify myocytes; and myogenin to identify rhabdomyocytes. - = analysis not performed; 0 = no staining of neoplastic cells; + = few stained neoplastic cells; ++ = moderate numbers of stained neoplastic cells; +++ = numerous stained neoplastic cells; (+) = staining of non-neoplastic cells; inc. = inconclusive IHC; NOS = not otherwise specified; HE = hematoxylin-eosin staining. Diagnoses in italics have not been confirmed with IHC but have the same morphology as the IHC-confirmed diagnosis in the same mouse.

| Mouse ID | Sex | Age (days) | Tissue | CD3 | Vimentin | CD68 | Desmin | Myogenin | Final diagnosis | Histomicrograph shown in Results |
|---|---|---|---|---|---|---|---|---|---|---|
| X340 | f | 145 | thymus | inc. | - | - | - | - | Lymphoma, NOS, solitary | |
| X547 | f | 113 | thymus | +++ | - | (+) | - | - | T-cell lymphoma, multicentric | |
| X405 | f | 118 | thymus | +++ | - | (+) | - | - | T-cell lymphoma, multicentric | |
| X339 | f | 131 | thymus | inc. | - | (+) | - | - | *T-cell lymphoma, multicentric* | |
| X509 | f | 176 | thymus | +++ | - | - | - | - | T-cell lymphoma, multicentric | |
| X319 | f | 225 | thymus | ++ | - | (+) | - | - | T-cell lymphoma, multicentric | |
| X214 | m | 258 | thymus | (+) | - | - | - | - | Histiocytic sarcoma, multicentric | |
| X503 | m | 146 | thymus | inc. | - | (+) | - | - | Lymphoma, NOS, multicentric | |
| X400 | m | 202 | thymus | + | - | (+) | - | - | Lymphoma, pleomorphic, multicentric | |
| X396 | m | 232 | thymus | + | - | (+) | - | - | Lymphoma, pleomorphic, solitary | |
| X315 | m | 70 | thymus | +++ | - | (+) | - | - | T-cell lymphoma, multicentric | |
| X502 | m | 97 | thymus | +++ | - | (+) | - | - | T-cell lymphoma, multicentric | Figure 3: panel A2 HE; panel A3 CD3 |
| X128 | m | 107 | thymus | +++ | - | (+) | - | - | T-cell lymphoma, multicentric | |
| X292 | m | 120 | thymus | +++ | - | (+) | - | - | T-cell lymphoma, multicentric | |
| X165 | m | 135 | thymus | +++ | - | - | - | - | T-cell lymphoma, multicentric | |
| X546 | m | 136 | thymus | +++ | - | (+) | - | - | T-cell lymphoma, multicentric | |
| X501 | m | 176 | thymus | +++ | - | (+) | - | - | T-cell lymphoma, multicentric | |
| X191 | m | 199 | thymus | +++ | - | (+) | - | - | T-cell lymphoma, multicentric | |
| X444 | m | 205 | thymus | +++ | - | (+) | - | - | T-cell lymphoma, multicentric | |
| X299 | m | 211 | thymus | +++ | - | (+) | - | - | T-cell lymphoma, multicentric | |
| X344 | m | 214 | thymus | +++ | - | (+) | - | - | T-cell lymphoma, multicentric | |
| X504 | m | 223 | thymus | +++ | - | (+) | - | - | T-cell lymphoma, multicentric | |
| X491 | m | 254 | thymus | +++ | - | (+) | - | - | T-cell lymphoma, multicentric | |
| X363 | m | 259 | thymus | +++ | - | (+) | - | - | T-cell lymphoma, multicentric | |
| X158 | m | 77 | thymus | +++ | - | (+) | - | - | T-cell lymphoma, solitary | |
| X393 | m | 108 | thymus | +++ | - | (+) | - | - | T-cell lymphoma, solitary | |
| X211 | m | 132 | thymus | +++ | - | (+) | - | - | T-cell lymphoma, solitary | |
| X481 | m | 137 | thymus | +++ | - | - | - | - | T-cell lymphoma, solitary | |
| X294 | m | 141 | thymus | +++ | - | (+) | . | . | T-cell lymphoma, solitary | |
| X152 | m | 188 | thymus | +++ | - | (+) | - | - | T-cell lymphoma, solitary | |
| X509 | f | 176 | subcutis+striated muscle | - | +++ | - | +++ | 0 | Leiomyosarcoma, solitary | |
| X341 | f | 130 | subcutis+striated muscle | - | ++ | (+) | + | 0 | Sarcoma, NOS, solitary | |
| X297 | m | 157 | subcutis+striated muscle | - | +++ | (+) | +++ | 0 | Leiomyosarcoma, solitary | |
| X400 | m | 202 | subcutis+striated muscle | (+) | +++ | (+) | +++ | 0 | Leiomyosarcoma, solitary | |
| X357 | m | 242 | subcutis+striated muscle | - | +++ | (+) | +++ | 0 | Leiomyosarcoma, solitary | |
| X387 | m | 92 | subcutis+striated muscle | - | +++ | (+) | +++ | + | Rhabdomyosarcoma, solitary | Supplementary Figure S2C: panel 5 & 6 HE; panel 7 Desmin; panel 8 Myogenin |
| X327 | m | 123 | subcutis+striated muscle | - | +++ | (+) | +++ | + | Rhabdomyosarcoma, solitary | |
| X339 | f | 131 | spleen | +++ | - | (+) | - | - | T-cell lymphoma, multicentric | |
| X319 | f | 225 | spleen | +++ | - | (+) | - | - | T-cell lymphoma, multicentric | |
| X312 | f | 217 | spleen | +++ | - | (+) | - | - | T-cell lymphoma, solitary | |
| X503 | m | 146 | spleen | - | - | - | - | - | Lymphoma, NOS, multicentric | |
| X488 | m | 207 | spleen | - | - | - | - | - | Lymphoma, NOS, multicentric | |
| X214 | m | 258 | spleen | + | - | +++ | | | Lymphoma, pleomorphic, solitary & Histiocytic sarcoma, multicentric | |
| X128 | m | 107 | spleen | +++ | - | (+) | - | - | T-cell lymphoma, multicentric | |
| X165 | m | 135 | spleen | +++ | - | (+) | - | - | T-cell lymphoma, multicentric | |
| X544 | m | 144 | spleen | +++ | - | (+) | - | - | T-cell lymphoma, multicentric | |
| X216 | m | 174 | spleen | +++ | - | (+) | - | - | T-cell lymphoma, multicentric | |
| X501 | m | 176 | spleen | +++ | - | (+) | - | - | T-cell lymphoma, multicentric | Figure 3: panel B2 HE; panel B3 CD3 |
| X386 | m | 184 | spleen | +++ | - | (+) | - | - | T-cell lymphoma, multicentric | |
| X191 | m | 199 | spleen | +++ | - | (+) | - | - | T-cell lymphoma, multicentric | |
| X444 | m | 205 | spleen | +++ | - | (+) | - | - | T-cell lymphoma, multicentric | |
| X299 | m | 211 | spleen | +++ | - | (+) | - | - | T-cell lymphoma, multicentric | |
| X344 | m | 214 | spleen | +++ | - | - | - | - | T-cell lymphoma, multicentric | |
| X497 | m | 223 | spleen | +++ | - | (+) | - | - | T-cell lymphoma, multicentric | |
| X300 | m | 225 | spleen | +++ | - | (+) | - | - | T-cell lymphoma, multicentric | |
| X363 | m | 259 | spleen | +++ | - | (+) | - | - | T-cell lymphoma, multicentric | |
| X162 | m | 175 | spleen | +++ | - | +++ | - | - | T-cell lymphoma, multicentric & Histiocytic sarcoma, multicentric | |
| X504 | m | 223 | spleen | +++ | - | +++ | - | - | T-cell lymphoma, multicentric & Histiocytic sarcoma, solitary | |
| X312 | f | 217 | soft tissue, hind leg | - | +++ | (+) | +++ | 0 | Leiomyosarcoma, solitary | |
| X498 | m | 118 | soft tissue, hind leg | - | +++ | (+) | 0 | 0 | Sarcoma, NOS, solitary | |
| X211 | m | 132 | soft tissue, front leg | - | +++ | (+) | ++ | 0 | Leiomyosarcoma, solitary | |
| X328 | m | 166 | soft tissue, front leg | - | ++ | (+) | ++ | 0 | Leiomyosarcoma, solitary | |
| X222 | m | 82 | soft tissue, front leg | - | +++ | (+) | 0 | 0 | Sarcoma, NOS, solitary | |
| X543 | m | 129 | skin | - | +++ | (+) | ++ | 0 | Leiomyosarcoma, solitary | |
| X294 | m | 141 | skin | - | +++ | (+) | +++ | 0 | Leiomyosarcoma, solitary | |
| X545 | m | 148 | skin | - | +++ | (+) | +++ | 0 | Leiomyosarcoma, solitary | Figure 3: panel C2 HE; panel C3 Desmin; panel C4 Myogenin |
| X504 | m | 223 | skin | - | +++ | (+) | +++ | 0 | Leiomyosarcoma, solitary | |
| X396 | m | 232 | skin | - | +++ | (+) | +++ | 0 | Leiomyosarcoma, solitary | Supplementary Figure S2C: panel 1 & 2 HE; panel 3 Desmin; panel 4 Myogenin |
| X546 | m | 136 | salivary glands, submandibular | +++ | - | (+) | - | - | T-cell lymphoma, multicentric | Supplementary Figure S2A: panel 9 & 10 HE; panel 11 CD3 |
| X191 | m | 199 | salivary glands, submandibular | +++ | - | (+) | - | - | T-cell lymphoma, multicentric | |
| X344 | m | 214 | salivary glands, submandibular | +++ | - | (+) | - | - | T-cell lymphoma, multicentric | |
| X300 | m | 225 | salivary glands, submandibular | +++ | - | (+) | - | - | T-cell lymphoma, multicentric | |
| X546 | m | 136 | salivary glands, sublingual | +++ | - | (+) | - | - | T-cell lymphoma, multicentric | Supplementary Figure S2A: panel 9 & 10 HE; panel 11 CD3 |
| X191 | m | 199 | salivary glands, parotid | +++ | - | (+) | - | - | T-cell lymphoma, multicentric | Supplementary Figure S2A: panel 5 & 6 HE; panel 7 CD3 |
| X300 | m | 225 | salivary glands, parotid | +++ | - | (+) | - | - | T-cell lymphoma, multicentric | |
| X400 | m | 202 | lymph nodes, submandibular | + | - | (+) | - | - | Lymphoma, pleomorphic, multicentric | |
| X546 | m | 136 | lymph nodes, submandibular | +++ | - | (+) | - | - | T-cell lymphoma, multicentric | |
| X162 | m | 175 | lymph nodes, submandibular | +++ | - | (+) | - | - | T-cell lymphoma, multicentric | |
| X300 | m | 225 | lymph nodes, submandibular | +++ | - | (+) | - | - | T-cell lymphoma, multicentric | Supplementary Figure S2B : panel 7 & 8 HE; panel 9 CD3 |
| X363 | m | 259 | lymph nodes, submandibular | - | - | - | - | - | *T-cell lymphoma, multicentric* | |
| X488 | m | 207 | lymph nodes, mesenteric | - | - | - | - | - | Lymphoma, NOS, multicentric | |
| X400 | m | 202 | lymph nodes, mesenteric | + | - | (+) | - | - | Lymphoma, pleomorphic, multicentric | Supplementary Figure S2A: panel 12 & 13 HE; panel 14 CD3 |
| X544 | m | 144 | lymph nodes, mesenteric | +++ | - | (+) | - | - | T-cell lymphoma, multicentric | Supplementary Figure S2B: panel 10 & 11 HE; panel 12 CD3 |
| X191 | m | 199 | lymph nodes, mesenteric | +++ | - | (+) | - | - | T-cell lymphoma, multicentric | |
| X497 | m | 223 | lymph nodes, mesenteric | +++ | - | (+) | - | - | T-cell lymphoma, multicentric | |
| X363 | m | 259 | lymph nodes, mesenteric | +++ | - | (+) | - | - | T-cell lymphoma, multicentric | |
| X491 | m | 254 | lymph nodes, mediastinum | - | +++ | (+) | ++ | 0 | Leiomyosarcoma in lymph node (metastasis) | |
| X319 | f | 225 | lymph nodes, inguinal | +++ | - | (+) | - | - | T-cell lymphoma, multicentric | |
| X128 | m | 107 | lymph nodes, inguinal | +++ | - | (+) | - | - | T-cell lymphoma, multicentric | |
| X165 | m | 135 | lymph nodes, inguinal | +++ | - | (+) | - | - | T-cell lymphoma, multicentric | |
| X216 | m | 174 | lymph nodes, inguinal | +++ | - | (+) | - | - | T-cell lymphoma, multicentric | |
| X162 | m | 175 | lymph nodes, inguinal | +++ | - | (+) | - | - | T-cell lymphoma, multicentric | Supplementary Figure S2B: panel 1 & 2 HE; panel 3 CD3 |
| X501 | m | 176 | lymph nodes, inguinal | +++ | - | (+) | - | - | T-cell lymphoma, multicentric | |
| X191 | m | 199 | lymph nodes, inguinal | +++ | - | (+) | - | - | T-cell lymphoma, multicentric | |
| X497 | m | 223 | lymph nodes, inguinal | +++ | - | (+) | - | - | T-cell lymphoma, multicentric | |
| X319 | f | 225 | lymph nodes, axillar | +++ | - | (+) | - | - | T-cell lymphoma, multicentric | |
| X488 | m | 207 | lymph nodes, axillar | - | - | - | - | - | Lymphoma, NOS, multicentric | |
| X165 | m | 135 | lymph nodes, axillar | +++ | - | (+) | - | - | T-cell lymphoma, multicentric | |
| X544 | m | 144 | lymph nodes, axillar | +++ | - | (+) | - | - | T-cell lymphoma, multicentric | |
| X162 | m | 175 | lymph nodes, axillar | +++ | - | (+) | - | - | T-cell lymphoma, multicentric | Supplementary Figure S2B: panel 4 & 5 HE; panel 6 CD3 |
| X501 | m | 176 | lymph nodes, axillar | +++ | - | (+) | - | - | T-cell lymphoma, multicentric | |
| X363 | m | 259 | lymph nodes, axillar | +++ | - | (+) | - | - | T-cell lymphoma, multicentric | |
| X547 | f | 113 | lung | - | - | - | - | - | *T-cell lymphoma, multicentric* | |
| X405 | f | 118 | lung | - | - | - | - | - | *T-cell lymphoma, multicentric* | |
| X509 | f | 176 | lung | - | - | - | - | - | *T-cell lymphoma, multicentric* | |
| X319 | f | 225 | lung | - | - | - | - | - | *T-cell lymphoma, multicentric* | |
| X162 | m | 175 | lung | - | - | - | - | - | Histiocytic sarcoma, multicentric | |
| X214 | m | 258 | lung | - | - | - | - | - | Histiocytic sarcoma, multicentric | |
| X503 | m | 146 | lung | - | - | - | - | - | Lymphoma, NOS, multicentric | |
| X488 | m | 207 | lung | - | - | - | - | - | Lymphoma, NOS, multicentric | |
| X315 | m | 70 | lung | +++ | - | (+) | - | - | T-cell lymphoma, multicentric | |
| X502 | m | 97 | lung | - | - | - | - | - | *T-cell lymphoma, multicentric* | Figure 3: panel A4 HE |
| X128 | m | 107 | lung | - | - | - | - | - | *T-cell lymphoma, multicentric* | |
| X292 | m | 120 | lung | - | - | - | - | - | *T-cell lymphoma, multicentric* | |
| X546 | m | 136 | lung | +++ | - | (+) | - | - | T-cell lymphoma, multicentric | |
| X216 | m | 174 | lung | +++ | - | (+) | - | - | T-cell lymphoma, multicentric | |
| X386 | m | 184 | lung | - | - | - | - | - | *T-cell lymphoma, multicentric* | |
| X191 | m | 199 | lung | - | - | - | - | - | *T-cell lymphoma, multicentric* | |
| X299 | m | 211 | lung | - | - | - | - | - | *T-cell lymphoma, multicentric* | |
| X344 | m | 214 | lung | - | - | - | - | - | *T-cell lymphoma, multicentric* | |
| X497 | m | 223 | lung | - | - | - | - | - | *T-cell lymphoma, multicentric* | |
| X504 | m | 223 | lung | - | - | - | - | - | *T-cell lymphoma, multicentric* | |
| X491 | m | 254 | lung | +++ | . | (+) | - | - | T-cell lymphoma, multicentric | |
| X363 | m | 259 | lung | - | - | - | - | - | *T-cell lymphoma, multicentric* & Histiocytic sarcoma, multicentric | |
| X165 | m | 135 | lung | +++ | - | ++ | - | - | T-cell lymphoma, multicentric & Histiocytic sarcoma, solitary | |
| X544 | m | 144 | lung | +++ | - | ++ | - | - | T-cell lymphoma, multicentric & Histiocytic sarcoma, solitary | |
| X501 | m | 176 | lung | +++ | - | ++ | - | - | T-cell lymphoma, multicentric & Histiocytic sarcoma, solitary | |
| X547 | f | 113 | liver | - | - | - | - | - | *T-cell lymphoma, multicentric* | |
| X339 | f | 131 | liver | - | - | - | - | - | *T-cell lymphoma, multicentric* | |
| X319 | f | 225 | liver | - | - | - | - | - | *T-cell lymphoma, multicentric* | |
| X214 | m | 258 | liver | + | - | +++ | - | - | Histiocytic sarcoma, multicentric | |
| X300 | m | 225 | liver | - | - | - | - | - | Histiocytic sarcoma, solitary | |
| X488 | m | 207 | liver | - | - | - | - | - | Lymphoma, NOS, multicentric | |
| X128 | m | 107 | liver | - | - | - | - | - | *T-cell lymphoma, multicentric* | |
| X292 | m | 120 | liver | - | - | - | - | - | *T-cell lymphoma, multicentric* | |
| X165 | m | 135 | liver | +++ | - | (+) | - | - | T-cell lymphoma, multicentric | |
| X544 | m | 144 | liver | - | - | - | - | - | *T-cell lymphoma, multicentric* | |
| X216 | m | 174 | liver | +++ | - | (+) | - | - | T-cell lymphoma, multicentric | |
| X162 | m | 175 | liver | - | - | - | - | - | *T-cell lymphoma, multicentric* | |
| X501 | m | 176 | liver | - | - | - | - | - | *T-cell lymphoma, multicentric* | Figure 3: panel B4 HE |
| X386 | m | 184 | liver | - | - | - | - | - | *T-cell lymphoma, multicentric* | |
| X191 | m | 199 | liver | - | - | - | - | - | *T-cell lymphoma, multicentric* | |
| X299 | m | 211 | liver | +++ | - | (+) | - | - | T-cell lymphoma, multicentric | |
| X344 | m | 214 | liver | +++ | +++ | (+) | - | - | T-cell lymphoma, multicentric | |
| X497 | m | 223 | liver | - | - | - | - | - | *T-cell lymphoma, multicentric* | |
| X504 | m | 223 | liver | - | - | - | - | - | *T-cell lymphoma, multicentric* | |
| X363 | m | 259 | liver | +++ | - | +++ | - | - | T-cell lymphoma, multicentric & Histiocytic sarcoma, multicentric | |
| X319 | f | 225 | kidney | - | - | - | - | - | *T-cell lymphoma, multicentric* | |
| X488 | m | 207 | kidney | - | - | - | - | - | Lymphoma, NOS, multicentric | |
| X128 | m | 107 | kidney | - | - | - | - | - | *T-cell lymphoma, multicentric* | Supplementary Figure S2A: panel 1 & 2 HE |
| X292 | m | 120 | kidney | - | - | - | - | - | *T-cell lymphoma, multicentric* | |
| X165 | m | 135 | kidney | +++ | - | (+) | - | - | T-cell lymphoma, multicentric | |
| X544 | m | 144 | kidney | - | - | - | - | - | *T-cell lymphoma, multicentric* | |
| X216 | m | 174 | kidney | +++ | - | (+) | - | - | T-cell lymphoma, multicentric | |
| X162 | m | 175 | kidney | - | - | - | - | - | *T-cell lymphoma, multicentric* | |
| X501 | m | 176 | kidney | - | - | - | - | - | *T-cell lymphoma, multicentric* | |
| X191 | m | 199 | kidney | - | - | - | - | - | *T-cell lymphoma, multicentric* | |
| X299 | m | 211 | kidney | +++ | - | (+) | - | - | T-cell lymphoma, multicentric | |
| X344 | m | 214 | kidney | - | - | - | - | - | *T-cell lymphoma, multicentric* | |
| X497 | m | 223 | kidney | - | - | - | - | - | *T-cell lymphoma, multicentric* | |
| X504 | m | 223 | kidney | - | - | - | - | - | *T-cell lymphoma, multicentric* | |
| X363 | m | 259 | kidney | - | - | - | - | - | *T-cell lymphoma, multicentric* | |
| X400 | m | 202 | intrathoracal tissue | 0 | ++ | (+) | 0 | 0 | Sarcoma, NOS, solitary | |
| X491 | m | 254 | intraabdominal fat tissue | - | +++ | (+) | ++ | 0 | Leiomyosarcoma, metastatic | |
| X547 | f | 113 | heart | - | - | - | - | - | *T-cell lymphoma, multicentric* | |
| X509 | f | 176 | heart | +++ | - | (+) | - | - | T-cell lymphoma, multicentric | |
| X503 | m | 146 | heart | - | - | - | - | - | Lymphoma, NOS, multicentric | |
| X128 | m | 107 | heart | - | - | - | - | - | *T-cell lymphoma, multicentric* | |
| X292 | m | 120 | heart | - | - | - | - | - | *T-cell lymphoma, multicentric* | |
| X191 | m | 199 | heart | - | - | - | - | - | *T-cell lymphoma, multicentric* | |
| X344 | m | 214 | heart | - | - | - | - | - | *T-cell lymphoma, multicentric* | Supplementary Figure S2A: panel 8 HE |
| X504 | m | 223 | heart | - | - | - | - | - | *T-cell lymphoma, multicentric* | |
| X363 | m | 259 | appendicular skeleton, femur | +++ | - | (+) | - | - | T-cell lymphoma, multicentric | Supplementary Figure S2A: panel 3 HE; panel 4 CD3 |
