## Supplementary Table S4B for "Mice carrying nonsense mutant p53 develop frequent multicentric or metastatic tumors"

**Supplementary Table S4B.** Results of immunohistochemistry (IHC) analysis of a selection of neoplasias in *Trp53*$^{R210X/+}$ (HZ) mice. CD3 was used to identify cells of T lymphocyte lineage; vimentin to identify mesenchymal cells and verify sarcoma; CD68 to identify cells of histiocytic lineage; desmin to identify myocytes; and myogenin to identify rhabdomyocytes. - = analysis not performed; 0 = no staining of neoplastic cells; + = few stained neoplastic cells; ++ = moderate numbers of stained neoplastic cells; +++ = numerous stained neoplastic cells; (+) = staining of non-neoplastic cells; inc. = inconclusive IHC; NOS = not otherwise specified; HE = hematoxylin-eosin staining. Diagnoses in italics have not been confirmed with IHC but have the same morphology as the IHC-confirmed diagnosis in the same mouse.

| Mouse ID | Sex | Age (days) | Tissue | CD3 | Vimentin | CD68 | Desmin | Myogenin | Final diagnosis | Histomicrograph shown in Results |
|---|---|---|---|---|---|---|---|---|---|---|
| X188 | m | 513 | urogenitalia, testicle | (+) | ++ | (+) | - | - | Leydig cell tumor, malignant | |
| X096 | m | 493 | urogenitalia, seminal vesicle | - | (+) | (+) | ++ | 0 | Leiomyosarcoma, metastatic | |
| X188 | m | 513 | urogenitalia, seminal vesicle | (+) | +++ | (+) | +++ | 0 | Leiomyosarcoma, metastatic | Supplementary Figure S3B: panel 5 HE; panel 6 Desmin; panel 7 Myogenin; panel 8 CD68 |
| X212 | m | 413 | urogenitalia, seminal vesicle | (+) | +++ | (+) | ++ | 0 | Leiomyosarcoma, solitary | |
| X329 | m | 476 | urogenitalia, seminal vesicle | (+) | +++ | (+) | ++ | 0 | Leiomyosarcoma, solitary | |
| X215 | m | 489 | urogenitalia, seminal vesicle | - | +++ | (+) | +++ | 0 | Leiomyosarcoma, solitary | |
| X171 | m | 565 | urogenitalia, seminal vesicle | - | +++ | + | ++ | 0 | Leiomyosarcoma, solitary | |
| X096 | m | 493 | urogenitalia, prostate | - | (+) | (+) | + | 0 | Leiomyosarcoma, metastatic | |
| X148 | m | 458 | urogenitalia, prostate | (+) | +++ | (+) | +++ | - | Leiomyosarcoma, solitary | |
| X245 | m | 495 | urogenitalia, preputial gland | (+) | ++ | (+) | (+) | 0 | Carcinoma, adenosquamous, solitary | |
| X188 | m | 513 | urogenitalia, preputial gland | (+) | +++ | (+) | ++ | 0 | Leiomyosarcoma, metastatic | Supplementary Figure S3B: panel 1 HE; panel 2 Desmin; panel 3 Myogenin; panel 4 CD68 |
| X283 | m | 408 | urogenitalia, preputial gland | (+) | +++ | (+) | ++ | 0 | Leiomyosarcoma, solitary | |
| X309 | f | 636 | thymus | ++ | - | + | - | - | Lymphoma, pleomorphic, multicentric & Histiocytic sarcoma, multicentric | |
| X167 | f | 563 | thymus | +++ | ++ | ++ | (+) | 0 | T-cell lymphoma, multicentric & Histiocytic sarcoma, multicentric | |
| X392 | m | 526 | thymus | (+) | +++ | +++ | (+) | 0 | Histiocytic sarcoma, multicentric | |
| X170 | m | 296 | thymus | inc. | - | ++ | - | - | Lymphoma, NOS, multicentric & Histiocytic sarcoma, solitary | |
| X338 | m | 626 | thymus | + | - | (+) | - | - | Lymphoma, pleomorphic, multicentric | |
| X225 | m | 413 | thymus | ++ | - | (+) | - | - | Lymphoma, pleomorphic, solitary | |
| X321 | m | 483 | thymus | inc. | - | ++ | - | - | Lymphoma,NOS, solitary & Histiocytic sarcoma, solitary | |
| X141 | m | 361 | thymus | +++ | - | (+) | - | - | T-cell lymphoma, multicentric | |
| X290 | m | 698 | thymus | +++ | - | (+) | - | - | T-cell lymphoma, multicentric | |
| X349 | m | 621 | thymus | + | - | (+) | - | - | Lymphoma, pleomorphic, multicentric | |
| X404 | f | 461 | subcutis | - | +++ | (+) | +++ | 0 | Leiomyosarcoma, solitary | |
| X247 | f | 472 | subcutis | - | +++ | (+) | +++ | 0 | Leiomyosarcoma, solitary | |
| X326 | f | 567 | subcutis | - | +++ | (+) | +++ | 0 | Leiomyosarcoma, solitary | |
| X194 | f | 567 | subcutis | + | - | (+) | - | - | Lymphoma, pleomorphic, multicentric | |
| X377 | m | 299 | subcutis | - | +++ | (+) | +++ | 0 | Leiomyosarcoma, solitary | |
| X276 | m | 364 | subcutis | (+) | +++ | (+) | +++ | 0 | Leiomyosarcoma, solitary | |
| X379 | m | 376 | subcutis | - | +++ | (+) | ++ | 0 | Leiomyosarcoma, solitary | |
| X338 | m | 626 | subcutis | + | - | (+) | - | - | Lymphoma, pleomorphic, multicentric | |
| X197 | m | 587 | subcutis | (+) | +++ | (+) | ++ | 0 | Sarcoma, NOS, solitary | |
| X099 | f | 548 | spleen | (+) | +++ | +++ | (+) | 0 | Histiocytic sarcoma, solitary | |
| X202 | f | 492 | spleen | inc. | - | (+) | - | - | Lymphoma, NOS, multicentric | |
| X194 | f | 567 | spleen | + | +++ | (+) | (+) | 0 | Lymphoma, pleomorphic, multicentric | |
| X311 | f | 622 | spleen | + | +++ | (+) | (+) | 0 | Lymphoma, pleomorphic, multicentric | |
| X309 | f | 636 | spleen | ++ | - | (+) | - | - | Lymphoma, pleomorphic, multicentric | |
| X354 | f | 571 | spleen | ++ | - | +++ | . | . | Lymphoma, pleomorphic, multicentric & Histiocytic sarcoma, multicentric | |
| X248 | f | 633 | spleen | + | +++ | +++ | (+) | 0 | Lymphoma, pleomorphic, multicentric & Histiocytic sarcoma, multicentric | |
| X173 | f | 559 | spleen | ++ | - | (+) | - | - | Lymphoma, pleomorphic, solitary | |
| X250 | f | 572 | spleen | ++ | - | (+) | - | - | Lymphoma, pleomorphic, solitary | |
| X167 | f | 563 | spleen | +++ | +++ | ++ | (+) | 0 | T-cell lymphoma, multicentric & Histiocytic sarcoma, multicentric | Supplementary Figure S3A: panel 1, 4 & 7 HE; panel 2, 5 & 8 CD68; panel 3, 6 & 9 CD3 |
| X317 | m | 642 | spleen | (+) | +++ | (+) | + | 0 | Hemangiosarcoma, solitary | |
| X103 | m | 660 | spleen | (+) | +++ | +++ | (+) | 0 | Histiocytic sarcoma, multicentric | |
| X079 | m | 533 | spleen | (+) | +++ | +++ | (+) | (+) | Histiocytic sarcoma, solitary | |
| X105 | m | 687 | spleen | - | - | - | - | - | Histiocytic sarcoma, solitary | |
| X149 | m | 505 | spleen | (+) | +++ | (+) | ++ | 0 | Leiomyosarcoma, solitary | |
| X213 | m | 489 | spleen | + | - | (+) | - | - | Lymphoma, pleomorphic, multicentric | |
| X338 | m | 626 | spleen | + | - | (+) | - | - | Lymphoma, pleomorphic, multicentric | |
| X183 | m | 500 | spleen | ++ | +++ | + | (+) | 0 | Lymphoma, pleomorphic, multicentric & Histiocytic sarcoma, multicentric | |
| X392 | m | 526 | spleen | (+) | +++ | (+) | (+) | 0 | Sarcoma, NOS, solitary | |
| X298 | m | 566 | spleen | (+) | +++ | (+) | (+) | 0 | Sarcoma, NOS, solitary | |
| X337 | m | 612 | spleen | +++ | - | (+) | - | - | T-cell lymphoma, multicentric | Figure 4: panel C3 HE |
| X290 | m | 698 | spleen | + | - | (+) | - | - | T-cell lymphoma, multicentric | |
| X349 | m | 621 | spleen | inc. | - | (+) | - | - | *Lymphoma, pleomorphic, multicentric* | |
| X167 | f | 563 | soft tissue, hindleg | (+) | +++ | +++ | ++ | + | Sarcoma, NOS, solitary | |
| X174 | m | 527 | soft tissue, hind leg | - | +++ | (+) | ++ | ++ | Rhabdomyosarcoma, solitary | Figure 4: panel B2 HE; panel B3 Desmin; panel B4 Myogenin |
| X398 | m | 420 | soft tissue, hind leg | - | ++ | inc | inc | 0 | Sarcoma, NOS, solitary | |
| X263 | f | 555 | small intestine | - | - | (+) | - | - | Carcinoma, in situ, solitary | |
| X325 | f | 483 | small intestine | ++ | - | (+) | - | - | Lymphoma, pleomorphic, multicentric | |
| X105 | m | 687 | small intestine | - | +++ | (+) | (+) | (+) | Sarcoma, NOS, solitary | |
| X354 | f | 571 | skin | - | +++ | (+) | ++ | 0 | Leiomyosarcoma, solitary | |
| X239 | f | 358 | skin | - | +++ | (+) | +++ | +++ | Rhabdomyosarcoma, solitary | Supplementary Figure S3B: panel 9 HE; panel 10 Desmin; panel 11 Myogenin; panel 12 CD68 |
| X101 | m | 470 | skin | - | - | (+) | - | - | Carcinoma, squamous cell, solitary | |
| X275 | m | 564 | skin | (+) | +++ | (+) | ++ | 0 | Leiomyosarcoma, solitary | |
| X102 | m | 498 | skin | - | (+) | - | - | - | Sarcoma, NOS, solitary | |
| X149 | m | 505 | skin | (+) | +++ | +++ | ++ | 0 | Sarcoma, NOS, solitary | |
| X194 | f | 567 | salivary gland, submandibular | + | - | (+) | - | - | Lymphoma, pleomorphic, multicentric | |
| X194 | f | 567 | salivary gland, sublingual | + | - | (+) | - | - | Lymphoma, pleomorphic, multicentric | |
| X194 | f | 567 | salivary gland, parotid | + | - | (+) | - | - | Lymphoma, pleomorphic, multicentric | |
| X194 | f | 567 | pancreas | + | - | (+) | - | - | Lymphoma, pleomorphic, multicentric | |
| X354 | f | 571 | pancreas | ++ | +++ | (+) | (+) | 0 | Lymphoma, pleomorphic, multicentric | Supplementary Figure S3C: panel 3 HE; panel 4 CD3 |
| X365 | f | 485 | pancreas | inc. | ++ | + | + | 0 | Sarcoma, NOS, solitary | |
| X392 | m | 526 | pancreas | - | - | +++ | (+) | 0 | Histiocytic sarcoma, multicentric | |
| X096 | m | 493 | pancreas | - | (+) | (+) | + | 0 | Leiomyosarcoma, metastatic | |
| X183 | m | 500 | pancreas | ++ | ++ | ++ | (+) | 0 | Lymphoma, pleomorphic, multicentric & Histiocytic sarcoma, multicentric | |
| X228 | f | 359 | mammary gland | - | - | (+) | - | - | Carcinoma, adenosquamous, solitary | |
| X153 | f | 498 | mammary gland | - | - | (+) | - | - | Carcinoma, ductal, solitary | Supplementary Figure S3C: panel 1 & 2 HE |
| X173 | f | 559 | mammary gland | - | - | (+) | - | - | Carcinoma, metastatic | |
| X235 | f | 590 | mammary gland | - | +++ | + | + | (+) | Hemangiosarcoma, solitary | |
| X372 | f | 583 | mammary gland | - | - | - | - | - | Sarcoma, NOS, solitary | |
| X248 | f | 633 | lymph nodes, submandibular | - | +++ | +++ | (+) | 0 | Histiocytic sarcoma, multicentric | |
| X202 | f | 492 | lymph nodes, submandibular | - | - | - | - | - | Lymphoma, NOS, multicentric | |
| X194 | f | 567 | lymph nodes, submandibular | + | - | (+) | - | - | Lymphoma, pleomorphic, multicentric | |
| X167 | f | 563 | lymph nodes, submandibular | ++ | +++ | +++ | (+) | 0 | T-cell lymphoma, multicentric & Histiocytic sarcoma, multicentric | |
| X213 | m | 489 | lymph nodes, submandibular | - | - | - | - | - | *Lymphoma, pleomorphic, multicentric* | |
| X167 | f | 563 | lymph nodes, mesenteric | (+) | +++ | +++ | (+) | 0 | Histiocytic sarcoma, multicentric | |
| X202 | f | 492 | lymph nodes, mesenteric | - | - | - | - | - | Lymphoma, NOS, multicentric | |
| X325 | f | 483 | lymph nodes, mesenteric | ++ | - | (+) | - | - | Lymphoma, pleomorphic, multicentric | |
| X263 | f | 555 | lymph nodes, mesenteric | ++ | - | (+) | - | - | Lymphoma, pleomorphic, multicentric | |
| X194 | f | 567 | lymph nodes, mesenteric | + | - | (+) | - | - | Lymphoma, pleomorphic, multicentric | |
| X194 | f | 567 | lymph nodes, mesenteric | + | - | (+) | - | - | Lymphoma, pleomorphic, multicentric | |
| X354 | f | 571 | lymph nodes, mesenteric | ++ | +++ | (+) | (+) | 0 | Lymphoma, pleomorphic, multicentric | |
| X311 | f | 622 | lymph nodes, mesenteric | ++ | - | (+) | - | - | Lymphoma, pleomorphic, multicentric | |
| X248 | f | 633 | lymph nodes, mesenteric | - | - | ++ | - | - | *Lymphoma, pleomorphic, multicentric* & Histiocytic sarcoma, multicentric | |
| X365 | f | 485 | lymph nodes, mesenteric | inc. | ++ | + | + | 0 | Sarcoma, NOS, solitary | |
| X392 | m | 526 | lymph nodes, mesenteric | - | - | +++ | (+) | 0 | Histiocytic sarcoma, multicentric | |
| X103 | m | 660 | lymph nodes, mesenteric | - | +++ | +++ | (+) | 0 | Histiocytic sarcoma, multicentric | |
| X096 | m | 493 | lymph nodes, mesenteric | - | (+) | (+) | + | 0 | Leiomyosarcoma, metastatic | |
| X402 | m | 447 | lymph nodes, mesenteric | inc. | - | (+) | - | - | Lymphoma, NOS, multicentric | |
| X213 | m | 489 | lymph nodes, mesenteric | - | - | - | - | - | *Lymphoma, pleomorphic, multicentric* | |
| X338 | m | 626 | lymph nodes, mesenteric | + | - | (+) | - | - | Lymphoma, pleomorphic, multicentric | |
| X183 | m | 500 | lymph nodes, mesenteric | ++ | ++ | ++ | (+) | 0 | Lymphoma, pleomorphic, multicentric & Histiocytic sarcoma, multicentric | |
| X290 | m | 698 | lymph nodes, mesenteric | +++ | - | (+) | - | - | T-cell lymphoma, multicentric | |
| X349 | m | 621 | lymph nodes, mesenteric | + | - | (+) | - | - | Lymphoma, pleomorphic, multicentric | |
| X309 | f | 636 | lymph nodes, mediastinal | ++ | - | + | - | - | Lymphoma, pleomorphic, multicentric & Histiocytic sarcoma, multicentric | |
| X392 | m | 526 | lymph nodes, mediastinal | (+) | +++ | +++ | (+) | 0 | Histiocytic sarcoma, multicentric | |
| X213 | m | 489 | lymph nodes, inguinal | - | - | - | - | - | *Lymphoma, pleomorphic, multicentric* | |
| X141 | m | 361 | lymph nodes, inguinal | +++ | - | (+) | . | . | T-cell lymphoma, multicentric | |
| X337 | m | 612 | lymph nodes, inguinal | +++ | - | (+) | - | - | T-cell lymphoma, multicentric | Figure 4: panel C2 HE |
| X354 | f | 571 | lymph nodes, axillar | ++ | +++ | (+) | (+) | 0 | Lymphoma, pleomorphic, multicentric | |
| X311 | f | 622 | lymph nodes, axillar | + | - | (+) | - | - | Lymphoma, pleomorphic, multicentric | |
| X213 | m | 489 | lymph nodes, axillar | - | - | - | - | - | *Lymphoma, pleomorphic, multicentric* | |
| X338 | m | 626 | lymph nodes, axillar | + | - | (+) | - | - | Lymphoma, pleomorphic, multicentric | |
| X337 | m | 612 | lymph nodes, axillar | +++ | - | (+) | - | - | T-cell lymphoma, multicentric | |
| X318 | f | 414 | lung | - | - | (+) | - | - | Adenocarcinoma, pulmonary, solitary | |
| X167 | f | 563 | lung | (+) | +++ | +++ | (+) | 0 | Adenoma, pulmonary & Histiocytic sarcoma, multicentric | |
| X173 | f | 559 | lung | - | - | (+) | - | - | Carcinoma, metastasis | |
| X389 | f | 517 | lung | - | - | - | - | - | Histiocytic sarcoma, multicentric | |
| X252 | f | 542 | lung | - | - | - | - | - | Lymphoma, NOS, multicentric | |
| X354 | f | 571 | lung | - | - | - | - | - | *Lymphoma, pleomorphic, multicentric* | |
| X311 | f | 622 | lung | - | - | - | - | - | *Lymphoma, pleomorphic, multicentric* | |
| X309 | f | 636 | lung | ++ | - | + | - | - | Lymphoma, pleomorphic, multicentric & Histiocytic sarcoma, multicentric | |
| X325 | f | 483 | lung | + | +++ | +++ | +++ | 0 | Mesothelioma, metastatic | |
| X314 | f | 462 | lung | (+) | +++ | (+) | + | 0 | Osteosarcoma, metastasis | Figure 4: panel A3 HE |
| X195 | f | 485 | lung | (+) | +++ | ++ | (+) | 0 | Osteosarcoma, metastasis | |
| X360 | f | 495 | lung | (+) | - | (+) | - | - | Osteosarcoma, metastasis | |
| X337 | m | 612 | lung | - | - | - | - | - | *T-cell lymphoma, multicentric* | Figure 4: panel C4 HE |
| X268 | m | 413 | lung | - | - | (+) | - | - | Adenocarcinoma, pulmonary, solitary | |
| X189 | m | 464 | lung | - | - | (+) | - | - | Adenocarcinoma, pulmonary, solitary | |
| X330 | m | 558 | lung | - | - | - | - | - | Adenocarcinoma, pulmonary, solitary | |
| X290 | m | 698 | lung | - | - | - | - | - | Adenoma, pulmonary & *T-cell lymphoma, multicentric* | |
| X170 | m | 296 | lung | - | - | - | - | - | Lymphoma, NOS, multicentric | |
| X402 | m | 447 | lung | - | - | - | - | - | Lymphoma, NOS, multicentric | |
| X171 | m | 565 | lung | - | - | - | - | - | Lymphoma, NOS, solitary | |
| X213 | m | 489 | lung | - | - | - | - | - | *Lymphoma, pleomorphic, multicentric* | |
| X338 | m | 626 | lung | - | - | - | - | - | *Lymphoma, pleomorphic, multicentric* | |
| X183 | m | 500 | lung | ++ | +++ | ++ | (+) | 0 | Lymphoma, pleomorphic, multicentric & Histiocytic sarcoma, multicentric | |
| X349 | m | 621 | lung | - | - | - | - | - | *Lymphoma, pleomorphic, multicentric* | |
| X389 | f | 517 | liver | - | +++ | +++ | (+) | 0 | Histiocytic sarcoma, multicentric | |
| X202 | f | 492 | liver | - | - | - | - | - | Lymphoma, NOS, multicentric | |
| X252 | f | 542 | liver | - | - | - | - | - | Lymphoma, NOS, multicentric | |
| X311 | f | 622 | liver | - | - | - | - | - | *Lymphoma, pleomorphic, multicentric* | |
| X354 | f | 571 | liver | - | - | - | - | - | *Lymphoma, pleomorphic, multicentric* & Histiocytic sarcoma, multicentric | |
| X248 | f | 633 | liver | + | +++ | +++ | (+) | 0 | Lymphoma, pleomorphic, multicentric & Histiocytic sarcoma, multicentric | |
| X309 | f | 636 | liver | ++ | +++ | ++ | + | 0 | Lymphoma, pleomorphic, multicentric & Histiocytic sarcoma, multicentric | |
| X365 | f | 485 | liver | inc. | ++ | + | + | 0 | Sarcoma, NOS, solitary | |
| X337 | m | 612 | liver | - | - | - | - | - | *T-cell lymphoma, multicentric* | Figure 4: panel C5 HE |
| X290 | m | 698 | liver | ++ | +++ | +++ | (+) | 0 | *T-cell lymphoma, multicentric* & Histiocytic sarcoma, solitary | |
| X225 | m | 413 | liver | - | - | (+) | - | - | Adenoma, hepatocellular | |
| X215 | m | 489 | liver | - | - | (+) | - | - | Adenoma, hepatocellular | |
| X171 | m | 565 | liver | - | - | (+) | - | - | Adenoma, hepatocellular | |
| X317 | m | 642 | liver | - | - | (+) | - | - | Adenoma, hepatocellular | |
| X227 | m | 506 | liver | - | - | +++ | - | - | Adenoma, hepatocellular & Histiocytic sarcoma, solitary | |
| X392 | m | 526 | liver | (+) | +++ | +++ | (+) | 0 | Histiocytic sarcoma, multicentric | |
| X103 | m | 660 | liver | (+) | +++ | +++ | (+) | 0 | Histiocytic sarcoma, multicentric | |
| X350 | m | 624 | liver | (+) | +++ | +++ | (+) | 0 | Histiocytic sarcoma, solitary | |
| X402 | m | 447 | liver | - | - | - | - | - | Lymphoma, NOS, multicentric | |
| X213 | m | 489 | liver | - | - | - | - | - | *Lymphoma, pleomorphic, multicentric* | |
| X183 | m | 500 | liver | ++ | +++ | ++ | (+) | 0 | Lymphoma, pleomorphic, multicentric & Histiocytic sarcoma, multicentric | |
| X298 | m | 566 | liver | (+) | +++ | + | + | 0 | Sarcoma, NOS, solitary | |
| X311 | f | 622 | kidney | - | - | - | - | - | *Lymphoma, pleomorphic, multicentric* | |
| X309 | f | 636 | kidney | ++ | +++ | ++ | + | 0 | Lymphoma, pleomorphic, multicentric & Histiocytic sarcoma, multicentric | |
| X167 | f | 563 | kidney | - | - | - | - | - | *T-cell lymphoma, multicentric* | |
| X337 | m | 612 | kidney | - | - | - | - | - | *T-cell lymphoma, multicentric* | Figure 4: panel C6 HE |
| X290 | m | 698 | kidney | - | - | - | - | - | *T-cell lymphoma, multicentric* | |
| X183 | m | 500 | kidney | ++ | +++ | ++ | (+) | 0 | Lymphoma, pleomorphic, multicentric & Histiocytic sarcoma, multicentric | |
| X325 | f | 483 | intrathoracal tissue | + | +++ | +++ | +++ | 0 | Mesothelioma, metastatic | |
| X389 | f | 517 | intraabdominal fat tissue | - | - | - | - | - | Lipoma | |
| X096 | m | 493 | intraabdominal fat tissue | - | (+) | (+) | + | 0 | Leiomyosarcoma, metastatic | |
| X290 | m | 698 | heart | - | - | (+) | - | - | *T-cell lymphoma, multicentric* | |
| X213 | m | 489 | heart | - | - | - | - | - | *Lymphoma, pleomorphic, multicentric* | |
| X099 | f | 548 | Harderian gland | - | - | (+) | - | - | Adenoma, Harderian gland | |
| X314 | f | 462 | axial skeleton, thoracic spine | - | +++ | + | + | 0 | Osteosarcoma, axial, metastatic | Figure 4: panel A2 HE |
| X069 | f | 412 | axial skeleton, thoracic spine | - | ++ | ++ | + | 0 | Osteosarcoma, axial, solitary | |
| X342 | f | 446 | axial skeleton, thoracic spine | - | - | + | - | - | Osteosarcoma, axial, solitary | |
| X403 | f | 468 | axial skeleton, skull | - | - | (+) | - | - | Osteosarcoma, axial, solitary | |
| X099 | f | 548 | axial skeleton, sacrum | - | +++ | ++ | - | 0 | Osteosarcoma, axial, solitary | |
| X091 | f | 553 | axial skeleton, sacrum | - | - | ++ | - | - | Osteosarcoma, axial, solitary | |
| X406 | f | 453 | axial skeleton, rib cage | - | +++ | + | (+) | 0 | Osteosarcoma, axial, solitary | |
| X287 | f | 359 | axial skeleton, lumbar spine | - | +++ | + | + | + | Osteosarcoma, axial, solitary | |
| X485 | f | 408 | axial skeleton, lumbar spine | - | +++ | + | + | 0 | Osteosarcoma, axial, solitary | |
| X318 | f | 414 | axial skeleton, lumbar spine | - | +++ | + | + | 0 | Osteosarcoma, axial, solitary | |
| X302 | f | 426 | axial skeleton, lumbar spine | - | +++ | + | + | 0 | Osteosarcoma, axial, solitary | |
| X153 | f | 498 | axial skeleton, lumbar spine | - | - | + | - | - | Osteosarcoma, axial, solitary | |
| X252 | f | 542 | axial skeleton, lumbar spine | - | +++ | + | (+) | 0 | Osteosarcoma, axial, solitary | |
| X248 | f | 633 | axial skeleton, lumbar spine | - | +++ | + | (+) | 0 | Osteosarcoma, axial, solitary | |
| X167 | f | 563 | axial skeleton, lumbar spine | (+) | +++ | +++ | ++ | + | Sarcoma, NOS, solitary | |
| X370 | m | 514 | axial skeleton, lumbar spine | - | +++ | (+) | + | + | Osteosarcoma, axial, solitary | |
| X360 | f | 495 | appendicular skeleton, hind leg | - | +++ | + | + | 0 | Osteosarcoma, appendicular, metastatic | |
| X147 | f | 361 | appendicular skeleton, hind leg | - | +++ | ++ | (+) | 0 | Osteosarcoma, appendicular, solitary | |
| X155 | f | 540 | appendicular skeleton, hind leg | - | - | + | - | - | Osteosarcoma, appendicular, solitary | |
| X195 | f | 485 | appendicular skeleton, front leg | - | +++ | + | ++ | + | Osteosarcoma, appendicular, metastatic | |
| X250 | f | 572 | appendicular skeleton, front leg | - | +++ | + | + | 0 | Osteosarcoma, appendicular, solitary | |
| X365 | f | 485 | appendicular skeleton, front leg | - | ++ | (+) | + | 0 | Sarcoma, NOS, solitary | |
| X099 | f | 548 | appendicular skeleton, femur | - | +++ | ++ | - | 0 | Osteosarcoma, appendicular, solitary | |
| X167 | f | 563 | appendicular skeleton, femur | (+) | +++ | +++ | inc | 0 | Sarcoma, NOS, solitary | |
| X103 | m | 660 | appendicular skeleton, femur | - | +++ | +++ | (+) | 0 | Histiocytic sarcoma, multicentric | |
| X370 | m | 514 | adrenal gland | (+) | +++ | (+) | (+) | 0 | Adenoma, cortical | |
