## Supplementary Table S4C for "Mice carrying nonsense mutant p53 develop frequent multicentric or metastatic tumors"

**Supplementary Table S4C.** Results of immunohistochemistry (IHC) analysis of a selection of neoplasias in *Trp53* $^{+/+}$ (WT) mice. CD3 was used to identify cells of T lymphocyte lineage; vimentin to identify mesenchymal cells and verify sarcoma; CD68 to identify cells of histiocytic lineage; desmin to identify myocytes; and myogenin to identify rhabdomyocytes. - = analysis not performed; 0 = no staining of neoplastic cells; + = few stained neoplastic cells; ++ = moderate numbers of stained neoplastic cells; +++ = numerous stained neoplastic cells; (+) = staining of non-neoplastic cells; inc. = inconclusive IHC; NOS = not otherwise specified; HE = hematoxylin-eosin staining. Diagnoses in italics have not been confirmed with IHC but have the same morphology as the IHC-confirmed diagnosis in the same mouse.

| Mouse ID | Sex | Age (days) | Tissue | CD3 | Vimentin | CD68 | Desmin | Myogenin | Final diagnosis | Histomicrograph shown in Results |
|---|---|---|---|---|---|---|---|---|---|---|
| X156 | f | 693 | spleen | - | - | - | - | - | *Lymphoma, pleomorphic, multicentric* | |
| X131 | f | 769 | spleen | ++ | +++ | (+) | (+) | 0 | Lymphoma, pleomorphic, multicentric | |
| X291 | f | 810 | spleen | ++ | .- | (+) | - | - | Lymphoma, pleomorphic, multicentric | Supplementary Figure S4A 1 & 3; HE; 2 & 4; CD3 |
| X193 | f | 934 | spleen | ++ | - | (+) | - | - | Lymphoma, pleomorphic, multicentric | |
| Bx3 | f | 869 | spleen | ++ | - | ++ | - | - | Lymphoma, pleomorphic, multicentric & Histiocytic sarcoma, multicentric | |
| X172 | m | 724 | spleen | ++ | - | (+) | - | - | Lymphoma, pleomorphic, solitary | |
| X156 | f | 693 | small intestine | ++ | - | (+) | - | - | Lymphoma, pleomorphic, multicentric | |
| X156 | f | 693 | skin | (+) | - | (+) | - | - | Papilloma, solitary | |
| X088 | m | 791 | seminal vesicle | + | - | + | - | - | Lymphoma, pleomorphic, multicentric & Histiocytic sarcoma, multicentric | |
| X291 | f | 810 | pancreas | ++ | - | (+) | . | . | Lymphoma, pleomorphic, multicentric | |
| Bx3 | f | 869 | pancreas | ++ | - | ++ | - | - | Lymphoma, pleomorphic, multicentric & Histiocytic sarcoma, multicentric | |
| X088 | m | 791 | lymph nodes, submandibular | + | - | + | - | - | Lymphoma, pleomorphic, multicentric & Histiocytic sarcoma, multicentric | |
| X156 | f | 693 | lymph nodes, mesenteric | ++ | - | (+) | - | - | Lymphoma, pleomorphic, multicentric | |
| X131 | f | 769 | lymph nodes, mesenteric | ++ | - | (+) | - | - | Lymphoma, pleomorphic, multicentric | |
| X291 | f | 810 | lymph nodes, mesenteric | ++ | . | (+) | . | . | Lymphoma, pleomorphic, multicentric | |
| X193 | f | 934 | lymph nodes, mesenteric | ++ | - | (+) | - | - | Lymphoma, pleomorphic, multicentric | |
| Bx3 | f | 869 | lymph nodes, mesenteric | ++ | - | ++ | - | - | Lymphoma, pleomorphic, multicentric & Histiocytic sarcoma, multicentric | |
| X210 | m | 790 | lymph nodes, mesenteric | (+) | +++ | +++ | (+) | 0 | Histiocytic sarcoma, multicentric | |
| X352 | m | 781 | lymph nodes, mesenteric | ++ | - | (+) | - | - | Lymphoma, pleomorphic, multicentric | |
| X291 | f | 810 | lymph nodes, mediastinal | ++ | . | (+) | . | . | Lymphoma, pleomorphic, multicentric | |
| Bx3 | f | 869 | lymph nodes, mediastinal | ++ | - | ++ | - | - | Lymphoma, pleomorphic, multicentric & Histiocytic sarcoma, multicentric | |
| X210 | m | 790 | lymph nodes, mediastinal | (+) | +++ | +++ | (+) | 0 | Histiocytic sarcoma, multicentric | |
| X088 | m | 791 | lymph nodes, mediastinal | + | - | (+) | - | - | Lymphoma, pleomorphic, multicentric | |
| X088 | m | 791 | lymph nodes, inguinal | + | - | + | - | - | Lymphoma, pleomorphic, multicentric & Histiocytic sarcoma, multicentric | |
| X352 | m | 781 | lymph nodes, axillar | - | - | - | - | - | *Lymphoma, pleomorphic, multicentric* | |
| X088 | m | 791 | lymph nodes, axillar | + | - | + | - | - | Lymphoma, pleomorphic, multicentric & Histiocytic sarcoma, multicentric | |
| X193 | f | 934 | lung | - | - | - | - | - | Carcinoma, pulmonary, solitary | |
| X291 | f | 810 | lung | ++ | - | - | - | - | Lymphoma, pleomorphic, multicentric | Supplementary Figure S4A: panel 5 & 7 HE; panel 6 & 8 CD3 |
| X352 | m | 781 | lung | - | - | - | - | - | Adenoma, pulmonary | |
| X289 | m | 782 | lung | - | - | (+) | - | - | Adenoma, pulmonary | |
| X237 | m | 909 | lung | - | - | (+) | - | - | Adenoma, pulmonary | |
| X391 | m | 711 | lung | - | - | - | - | - | Adenoma, pulmonary, solitary | |
| X221 | m | 737 | lung | - | - | (+) | - | - | Carcinoma, pulmonary, solitary | Supplementary Figure S4C: panel 3 & 4 HE |
| X414 | m | 788 | lung | - | - | - | - | - | Histiocytic sarcoma, multicentric | |
| X210 | m | 790 | lung | ++ | +++ | +++ | (+) | 0 | Histiocytic sarcoma, multicentric | |
| X140 | m | 793 | lung | - | +++ | +++ | (+) | 0 | Histiocytic sarcoma, multicentric | Supplementary Figure S4B: panel 5 & 7 HE; panel 6 & 8 CD68 |
| B10 | f | 911 | liver | - | - | +++ | - | - | Adenoma, hepatocellular & Histiocytic sarcoma, solitary | |
| X193 | f | 934 | liver | - | - | (+) | - | - | Carcinoma, hepatocellular, solitary | |
| Bx5 | f | 841 | liver | (+) | + | +++ | (+) | 0 | Carcinoma, hepatocellular, solitary & Histiocytic sarcoma, solitary | |
| X291 | f | 810 | liver | - | - | - | - | - | Histiocytic sarcoma, solitary | |
| X172 | m | 724 | liver | - | - | (+) | - | - | Adenoma, hepatocellular | |
| X097 | m | 701 | liver | - | - | (+) | - | - | Carcinoma, hepatocellular, solitary | Supplementary Figure S4C: panel 1 & 2 HE |
| X414 | m | 788 | liver | (+) | +++ | +++ | (+) | 0 | Histiocytic sarcoma, multicentric | |
| X210 | m | 790 | liver | (+) | +++ | +++ | (+) | 0 | Histiocytic sarcoma, multicentric | |
| X088 | m | 791 | liver | - | - | +++ | - | - | Histiocytic sarcoma, multicentric | |
| X140 | m | 793 | liver | (+) | +++ | +++ | (+) | 0 | Histiocytic sarcoma, multicentric | Supplementary Figure S4B: panel 1 & 3 HE; panel 2 & 4 CD68 |
| X237 | m | 909 | liver | (+) | +++ | +++ | (+) | 0 | Histiocytic sarcoma, solitary | |
| X291 | f | 810 | kidney | - | - | - | - | - | Lymphoma, pleomorphic, multicentric | |
| X352 | m | 781 | kidney | - | - | (+) | - | - | Carcinoma, renal, solitary | |
| X310 | f | 758 | intraabdominal fat tissue | - | - | - | - | - | Lipoma | |
| X229 | f | 640 | brain | - | - | (+) | - | - | Carcinoma, pituitary gland, pars distalis, solitary | |
